# A matrikine organizes the dendritic cell–T cell triads that build tertiary lymphoid structures

**DOI:** 10.64898/2026.09.24.753585

**Authors:** Daniel J. Lagal, Duncan Hong, Athanasios Papadas, George S. Yacu, Emma Geatches, Nicholas Leschinsky, Yun Huang, Yaling Dou, Matteo Fields, Alicia Gibbons, Elsa Molina, Kersi Pestonjamasp, Peter T. Toth, Alexander Cicala, Joao Mamede, Jeffrey Schneider, Kristina A. Matkowskyj, Dustin Deming, Surabhi Naik, Fotis Asimakopoulos

## Abstract

Tertiary lymphoid structures (TLS) predict checkpoint-inhibition response, and conventional dendritic cells (cDC) are required to build and maintain them. What licenses tumor niches is not fully understood: chemokines recruit cells but do not specify TLS-archetypal DC:CD4⁺T:CD8⁺T triad composition. We show that the organizing signal is a matrix fragment. Proteoglycan versican (VCAN) proteolysis releases the matrikine versikine constitutively in CXCL13-replete germinal center light zones. In tumors, versikine is released inducibly within CXCL13-rich stromal niches. Versikine activates CD11b⁺ cDC, expands cDC1, and suffices for triad assembly. Versikine-conditioned DC engage progenitor-exhausted CD8⁺ T-cells (Tpex) and Tpex-sustaining, Th1-polarized follicular helper CD4⁺ T-cells. Excess unproteolyzed VCAN inverts this toward DC–Treg crosstalk and Tpex loss. Versikine (mRNA or recombinant protein) plus anti-PD-L1 eradicated experimental tumors reverse-translated to model adverse-prognosis, VCAN-proteolysis-weak patients from a prospective PD-1 blockade trial in oligometastatic colorectal cancer. The extracellular matrix instructs, rather than merely scaffolds, the multicellular units of antitumor immunity.

## INTRODUCTION

Ectopic multicellular structures resembling organized secondary lymphoid tissue have long been recognized as functional immunity hubs at sites of inflammation and cancer ^1–5^. These formations, termed tertiary lymphoid structures (TLS), have been an intense subject of scrutiny as their importance in orchestrating and fine-tuning anti-tumor immunity is better understood ^6^. Tumors bearing abundant TLS have generally been associated with favorable prognoses and improved responses to cancer immunotherapy ^2,7–10^. Therapeutic TLS induction has been postulated to overcome the efficacy barriers of modern immunotherapy ^11,12^.

However, several conceptual and practical challenges remain. First, various degrees of organization of lymphoid tissue along the tumor margin have been described ^13^, ranging from loose collections of antigen-presenting cells and lymphocytes, to complex lymph node-like architectures with mature follicles and germinal centers. Although it is tempting to consider these patterns as stepwise stages of a linear pathway, this view may be oversimplified; paradoxically, lesser-complexity formations may correlate more closely with anti-tumor immunity ^14–18^. Second, TLS presence may not always be favorable ^19^. Third, the foundational events during TLS formation are not yet fully elucidated and as a result, TLS have been challenging to therapeutically induce *in vivo*, particularly in human subjects ^20^.

Most attempts to understand the earliest stages of TLS formation and replicate them have focused on cellular actors and the roles of chemokines and cytokines that orchestrate cellular crosstalk. The role of extracellular matrix (ECM) and matrix remodeling in TLS organization is much less well-understood and has been confounded by the concept of tumor stroma acting as an “immune barrier” ^18,21^. Therapeutically, this notion has fostered attempts to disrupt tumor stroma which have generally been unsuccessful and, in some cases, harmful ^22,23^.

The lack of broad success of targeting tumor stroma can be attributed in part to the fact that prior approaches have ignored stromal heterogeneity and the complexity of signaling networks that operate within tumor stroma. Wound healing can provide an instructive conceptual framework in this regard ^24^. The early stages of wound healing are characterized by immune trafficking and intense cellular activity that aims to restore homeostasis. A key anatomic and functional framework for this activity is a permissive provisional ECM, rich in proteoglycans and glycoproteins ^25^. Later in wound evolution, provisional matrix is replaced by fibrotic stroma, signaling permanent wound closure. The transition from provisional to fibrotic ECM is facilitated by signals that are known to promote immunosuppression and tumor immune evasion, such as TGF-β ^26,27^. Cancers have been likened to “wounds that do not heal” and immunologically “cold” tumors would be expected to mirror the cessation of immune activity, immune evasion and immune exclusion associated with wound closure and fibrosis ^24^.

Conversely, the persistence of stromal ECM composition similar to wound provisional ECM would be expected to characterize “hot” tumors that would be susceptible to modern immunotherapies, such as immune checkpoint inhibition (ICI). Indeed, published work from our laboratory and others has shown that the regulated proteolysis of the large matrix proteoglycan versican (VCAN) ^28^, a cardinal event in provisional ECM remodeling in development and wound healing ^29–34^, is associated with T-cell inflammation. This association is conserved among multiple human solid and liquid tumor types ^35–41^. VCAN proteolysis at a site predicted to release a 441-aa N-terminal fragment, versikine (hereafter abbreviated to Vkine) ^42,43^, is essential for the development of circulatory and skeletal systems ^44,45^. Mice with a disrupted cleavage site demonstrate developmental abnormalities ^46,47^ and accelerated wound closure ^46^. We have previously demonstrated that Vkine promotes the abundance and activation of antigen-presenting cells, resulting in T-cell infiltration in multiple experimental models ^39^. Thus, we hypothesized that VCAN proteolysis results in T-cell inflammation at least in part through the activities of the released fragment, Vkine.

Recently, we demonstrated that VCAN proteolysis carries predictive value with respect to clinical outcomes following ICI, beyond its well-established correlation with histological T-cell infiltration. In a Phase 1b trial of mismatch repair-proficient oligometastatic colorectal cancer ^41^, we stratified patients according to their VCAN proteolysis status. We have previously described opposing activities of Vkine (immunostimulatory) and its parent macromolecule VCAN (immunosuppressive) ^30,43,48^. Patients with robust VCAN proteolysis and low accumulation of intact VCAN (termed VCAN-proteolysis-predominant, VPP) were in the most favorable prognostic group ^41^: all VPP patients were alive at 4.1-year median follow-up, with median recurrence-free survival of 3.78 years against 0.83 years for patients displaying low rates of VCAN proteolysis (termed VCAN-proteolysis-weak, VPW). Within the VPW subgroup, patients demonstrating excess parental unprocessed VCAN (VPW-VCAN^hi^) had particularly adverse outcomes.

In the current manuscript we highlight VCAN proteolysis, and by extension, provisional ECM remodeling, at the crossroads of lymphoid neogenesis, wound healing, and adaptive immunity. We show that constitutive VCAN proteolysis in germinal center (GC) light zones (LZ), colocalizing with CXCL13^+^ gradients ^49^, is mirrored by inflammatory VCAN proteolysis within tumor stromal CXCL13-rich niches. We further demonstrate that Vkine organizes tri-partite immune hubs (“immune triads”) critical for TLS assembly and maintenance ^6,13^ and immunotherapy responses ^50^. Therapeutic Vkine renders experimental tumors exquisitely sensitive to ICI. A matrix fragment organizes chemokine-recruited immune cells into TLS-foundational multicellular units.

## MATERIALS AND METHODS

### Lead contact

Further information and requests for resources and reagents should be directed to, and will be fulfilled by, the lead contact, Fotis Asimakopoulos.

### Data availability

Single-cell RNA sequencing data generated in this study are available through GEO (GSE348233). Publicly available scRNA-seq data from NCT02837263 were obtained from GEO (GSE316301).

### Human tissue microarrays (TMA)

Human non-small cell lung cancer (NSCLC) TMA (#BC041115e) and lymph node and tonsil TMA (#LY241i) were commercially obtained from US Biomax (Derwood, MD). Freshly cut tissue sections were obtained and used for immunohistochemistry analysis.

### Animal strains and animal research ethics

C57BL/6J (JAX, #000664), BALB/cJ (JAX, #000651), B6.129S(C)-Batf3tm1Kmm/J (*Batf3*-/-, JAX #103755) mouse strains were purchased from the Jackson Laboratory (Bar Harbor, ME, USA). Mice were housed, cared for and used in accordance with the Guide for Care and Use of Laboratory Animals (NIH Publication 86-23) under IACUC-approved protocols #S19109 at the University of California San Diego and #25-010 at Rush University Medical Center.

### Cell lines

LLC (ATCC, #CRL-1642), CT26 (ATCC, #CRL2638), MC38 (Kerafast, #ENH204-FP), and HEK293T (ATCC, #CRL-3216) cells were cultured in complete DMEM medium (Corning, #10-013) with 10% Fetal Bovine Serum (FBS) (Gibco, #A5209401), 50 μM 2-mercaptoethanol (Gibco, #21985023), 100 U/mL Penicillin, 100 μg/mL Streptomycin and 25 ng/mL Amphotericin B cocktail (Gibco, #15240096) and 1X GlutaMAX supplement (Gibco, #35050061). Cells were cultured for 5-20 passages, maintained in sterile conditions and grown in a controlled atmosphere at 37°C and 5% CO2.

### Constructs

pLenti6-UbC-Vkine-HA and pSecTag2-Vkine-Myc-His have been previously described ^35,45^. Empty vector (EV) backbones without Vkine open reading frames were used as controls. For lentivirus transduction, psPAX2 (Addgene, #12260) and pCMV-VSVg (Addgene, #8454) were used.

For Vkine mRNA synthesis, DNA encoding Vkine-HA or Vkine-Myc was PCR-amplified from pLenti6-UbC-VKine-HA or pSecTag2-Vkine-Myc-His, respectively, and infusion-cloned into the template plasmid provided with the Takara IVTpro mRNA Synthesis System (Cat. #6141). A control mRNA was generated in parallel by cloning DNA encoding Thy1.1 into the same template. Constructed vectors were transformed into NEB Stable competent cells (NEB, #C3040H) for amplification. mRNA was synthesized according to protocol from IVTpro mRNA Synthesis Systems.

All plasmids were prepped and purified using QIAprep Spin Miniprep Kit (Qiagen, #27104).

### Lentiviral transduction

HEK293T cells in the logarithmic growth phase were seeded in 12-well plates. The next day, cells were transfected with a mixture of 0.9 μg of ps-PAX2 (packaging plasmid), 0.6 μg of pVSV-G (envelope plasmid), and 1.5 μg of transfer plasmid encoding open reading frames in 100 μL of optiMem containing 0.075 mg/mL PEI reagent per well. After 30 hours, medium was collected, filtered using 0.45 μm filters, and added to 2×10^5 target cells, plated on 6-well plates the day before. Target cells were transduced for 72 hours, then cells were trypsinized and plated in 10 cm dishes. Cells were then selected through several passages with fresh medium containing antibiotics for mammalian cell selection, e.g. blasticidin 10 μg/mL (InvivoGene, #ant-bl-05). HA-tagged Vkine expression was confirmed by western blotting using anti-HA antibody.

### *In vitro* proliferation assay

*In vitro* proliferation was assessed using the CellTiter-Glo® Luminescent Cell Viability Assay (Promega, #G7570). CT26 and MC38 cells were seeded at a density of 1×10^4 cells per well in white 96-well plates (Nunc™ MicroWell™, Thermo Fisher, #136101). Luminescence was measured at 24, 48, 72, and 96 hours post-seeding according to the manufacturer’s protocol. For each cell line, a standard curve was generated by plating known cell numbers (0-1.5×10^5) and correlating cell number with luminescent signal by linear regression, allowing conversion of luminescence values to absolute cell numbers. Cell numbers were normalized to the growth surface area of each well.

### Tumor cell inoculation and tumor growth measurements

At roughly 80% confluency, cultured tumor cell lines were harvested by trypsinization and washed/resuspended in PBS. Unless stated otherwise, 5×10^5 cells (LLC, MC38, or CT26) were injected subcutaneously (s.c.) into the right flank of recipient C57BL/6 (for LLC and MC38) or BALB/cJ (for CT26) mice. Tumor growth was measured every three days using a digital caliper, and tumor volumes were approximated with the formula: Tumor volume = (largest tumor diameter) x (perpendicular tumor diameter)^2 divided by 2.

### Tissue processing

Unless stated otherwise, tumors were excised at 21 days after implantation. For subsequent analysis by flow cytometry, tumors were cut into small pieces and digested using the mouse tumor dissociation kit (Miltenyi Biotec, #130-096-730) in an automatic motion with the gentle MACS dissociator (Miltenyi Biotec, #130-093-235). Tissue was passed through a 70μm cell strainer (Fisher Scientific, #08-771-2) with FACS buffer (PBS with 2% FBS). Spleens were excised, disaggregated using a syringe plunger and filtered through a 70 μm cell strainer with FACS buffer. Cell suspensions were centrifuged for 7 minutes at 300 x g and red blood cells were lysed using ACK buffer (Quality Biological, #118-156-101) before proceeding with antibody staining.

For RNA isolation, whole tissue was mechanically homogenized in QIAzol Lysis Reagent (Qiagen, #79306) and processed per manufacturer recommendations.

### Antibodies

Antibodies are listed in Supp. Table S1.

### Flow cytometry analysis

Single cell suspensions were stained with viability probe Ghost Dye 780 (Tonbo Biosciences, #13-0865) for 10 minutes following manufacturer’s guidelines. Cells were washed and blocked with FACS buffer (PBS with 2% FBS) containing Fc block (BioLegend, #156604) in a 1:50 proportion for 10 minutes at room temperature. Subsequently, cells were incubated with desired antibodies for extracellular markers (Supp. Table S1) in 1:100 concentration with FACS buffer for at least 30 minutes at 4°C. Cells were then washed and resuspended in FACS buffer. For absolute quantitation, 10 μL of Precision Count beads (Biolegend, #424902) were added to samples. All flow cytometry data were collected using a LSR Fortessa flow cytometer (BD Biosciences) and analyzed using the FlowJo software (v10.10.1).

### SDS-PAGE and western blotting

Protein lysates were generated using NP-40 lysis buffer (Thermo Fisher, #J60766.AP). Lysates were shaken on an automatic wheel for 15 minutes and centrifuged for 10 minutes at 10,000 x g. Protein concentration was measured through absorbance at 592 nm in the TECAN plate reader following Pierce BCA protein assay (Thermo Fisher, #23225). 70 μg of protein were used in combination of 4x Laemmli buffer (Bio-rad, #161-0747) (supplemented with 2-mercaptoethanol (Bio-rad, #1610710) in a 1:10 proportion). Samples were heated at 95°C for 5 minutes. Prepared samples were loaded onto 4–15% Mini-PROTEAN® TGX™ Precast Protein polyacrylamide gels (Bio-rad, #4561083DC). Electrophoresis was performed for 90 minutes at 120V with Tris-Glycine-SDS buffer (Bio-rad, #1610732). Proteins were transferred to 0.2 μm PVDF membrane using transfer pack (Bio-rad, #1610732) and using a wet system with 25 mM Tris and 192 mM Glycine buffer for 2 hours and 30 minutes at 0.25 Amp at 4°C. Membrane was blocked with TBS-T containing 5% blot-grade blocker (Bio-rad, #1706404). Antibodies (Supp. Table S1) were incubated overnight at 4°C in 1:1000 dilution in TBS-T containing 1% blot-grade blocker. Membrane was washed and incubated with anti-IgG rabbit antibody conjugated with horseradish peroxidase in TBS-T for 90 minutes at room temperature. Specific proteins were detected using Pierce™ ECL Western Blotting Substrate (Thermo Fisher, #32106). Chemiluminescence intensity was quantified with ImageJ (software 1.54p).

### Volumetric immunofluorescence microscopy

Subcutaneous tumors were harvested on day 21 post-implantation. Tumors were excised, skin and fibroadipose tissue removed, and fixed using 4% PFA overnight. Tumors were dehydrated in a sucrose gradient (10% and 20% for 2 hours each and 30% overnight) and embedded in TissueTek OCT. Embedded tumor samples were stored at –80 °C. Consecutive sections of 25 µm thickness were cut. Sections were permeabilized and blocked with 0.25% Triton X100, 2% BSA and 3% goat serum (Thermo Fisher, #31872) dissolved in PBS overnight. Sections were stained with desired primary antibodies (see Supp. Table S1) in 0.2% Triton X100 and 3% goat serum in PBS overnight followed by washes (0.2% Triton X-100) and incubated with desired fluorochrome-conjugated secondary antibodies (Supp. Table S1) for 3 hours. Sections were washed and stained with 1 µg/mL DAPI (Thermo Fisher, #D3571). Stained sections were mounted with prolong glass anti-fade mounting medium (Thermo Fisher, #P36982) and analyzed on Zeiss Laser Scanning Confocal (LSM) 980. Immunofluorescence was analyzed on images acquired using a Zeiss Laser Scanning Confocal (LSM) 980 equipped with a 63X LD Plan-Apochromat 1.2 NA objective. For each field of view, XY plane was conserved between acquisitions and z-stack comprising 25 optical sections was collected at a step size of 1 µm using sequential laser excitation at 405, 488, 561 and 639 nm wavelength. Pinhole was set to 1 Airy unit, and images were captured at a resolution of 1024×1024 pixels with 8-bit depth.

### Immunofluorescence and immunohistochemistry of formalin-fixed tissue

Paraffin-embedded unstained 5 μm-thick human TMA and human tonsil sections were deparaffinized and rehydrated using standard methods. Sections for VCAN staining were treated with a solution of 125 mU/mL Chondroitinase ABC (Sigma, #C2905) in PBS for 45 minutes at 37°C. Antigen retrieval was carried out in citrate buffer, pH 6.0 (Vector Laboratories, #H-3300). Endogenous peroxidase activity was quenched using BLOXALL (Vector, #SP-6000). Primary antibodies are listed in Supp. Table S1. For immunohistochemistry, secondary antibodies conjugated to biotin (listed in Supp. Table S1) were incubated with avidin-conjugated HRP (Vector, #PK-4000) and chromogenic staining was generated using DAB substrate (Vector, #SK-4100). Slides were counterstained with hematoxylin (Vector, #H-3404-100). For immunofluorescence, secondary antibodies containing fluorochromes and nuclear staining with DAPI were used as described in the previous section. Stained slides were examined using an Echo Revolve microscope with an attached digital camera. Immunofluorescence sections were examined using Zeiss Laser Scanning Confocal (LSM) 980, Evident VS200 or Keyence BZX 710 microscope.

### Triad quantification in human NSCLC tissue microarray

Following acquisition and stitching, individual tissue scans were extracted for the specific cell markers based on hue using the Hybrid Cell Count feature in the Keyence BZX analyzer. Color masks were assigned to each cell type (red for XCR1, green for CD8 and blue for CD4), and mask images were analyzed using the GA3 feature in Elements (Nikon) software. The number of intersecting points between masked areas was counted either as such or following dilation of the masked areas by 0.7 µm. Results were expressed as XCR1+ touching either CD4+ or CD8+ or both. This analysis was performed on single tissue sections in two dimensions.

### Triad quantification in murine tumors by volumetric microscopy

Raw z-stacks acquired at 63X were imported into Imaris (Oxford Instruments, v9.7) and Fiji (ImageJ v1.54p). In Imaris, 3D surfaces were generated for nuclei and membrane using intensity-based thresholding, with segmentation parameters optimized empirically and held constant across all samples. Imaris reconstructions were used for three-dimensional visualization and for object-level verification of candidate assemblies (Fig. 1K). Intercellular distances were determined in Fiji. Candidates were identified in Fiji by scrolling through the full z-stack. For each candidate heterologous cell pair, a line was drawn across the apposed cell borders, beginning and ending within the marker-negative interior of each cell. Fluorescence intensity profiles for both markers were generated along the line and scaled to its length, and the separation between the two marker intensity maxima was recorded. Because CD11c, CD4 and CD8a are surface antigens, this peak-to-peak separation corresponds to the distance between the apposed plasma membranes. Immune triad formation was defined as an individual CD11c⁺ cell for which all three pairwise membrane-to-membrane distances (CD11c⁺–CD8⁺, CD11c⁺–CD4⁺ and CD8⁺–CD4⁺) did not exceed 5 µm. Where the nearest CD11c⁺ membrane maximum corresponded to a dendritic process rather than the cell body, the contact was retained. Triad counts (# of CD11c^+^ in triads) were normalized to total CD11c⁺ counted in the same manner and extrapolated to 1 mm³ volume.

**Fig. 1.**
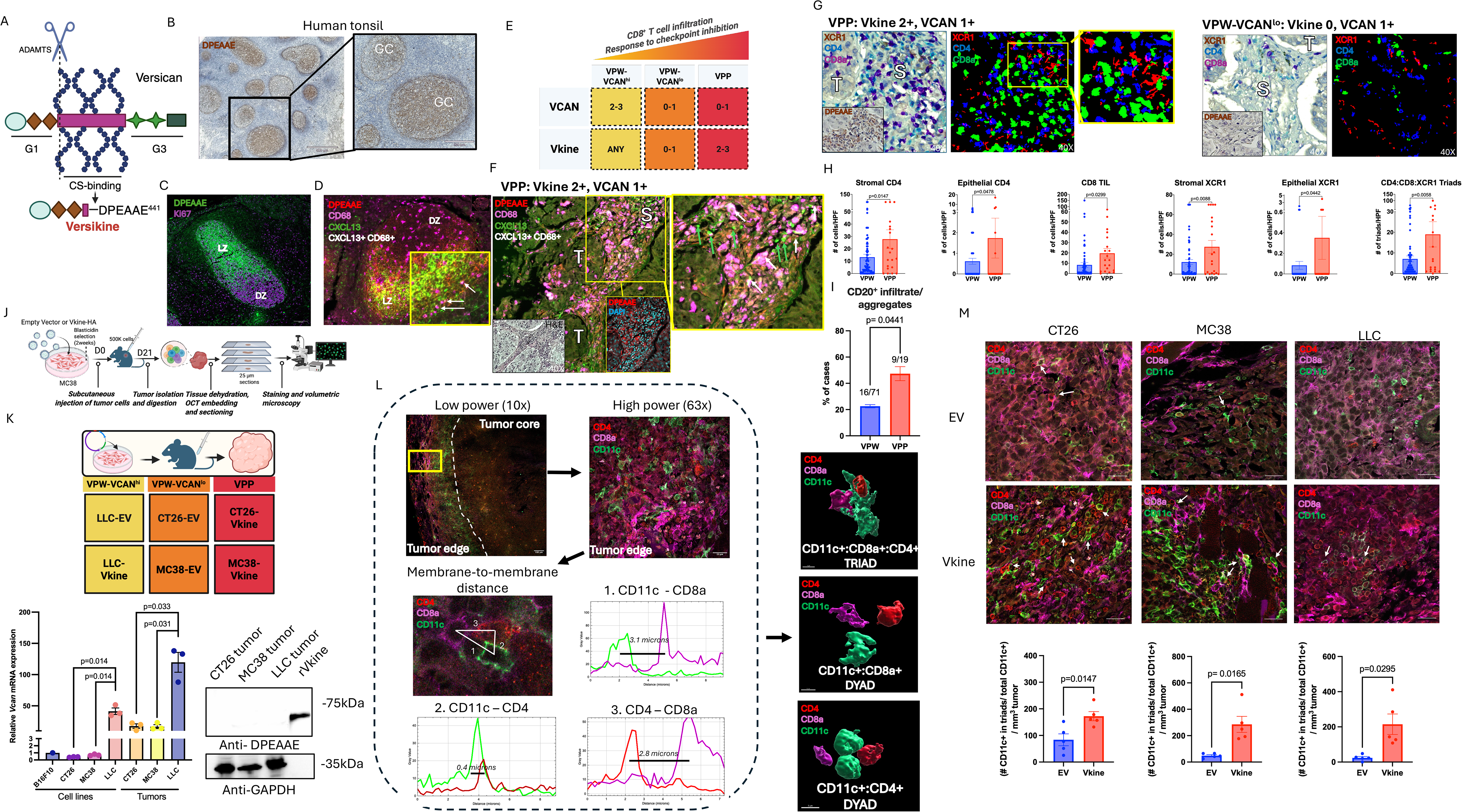
Versican (VCAN) proteolysis localizes within CXCL13 niches in secondary lymphoid tissue and tumor stroma. Its bioactive product, Vkine, promotes immune triad formation in tumors. A: Regulated VCAN proteolysis at position 441-442 of the VCAN-V1 isoform generates a fragment, Vkine, with distinct bioactivity. The proteolysis event unmasks a neoepitope (DPEAAE) that can be immunohistochemically detected. B: Immunohistochemistry of human tonsillar tissue with an anti-DPEAAE antibody reveals polarized staining within germinal centers (GC) (chromogen: DAB, counterstain: hematoxylin). 4X objective: scalebar 500 μm, 10X objective: scalebar 230 μm. C: Immunofluorescence (IF) staining with a Ki67 antibody demonstrates that anti-DPEAAE staining correlates with the Ki67-sparse light zone (LZ). Ki67 is a marker of centroblast division within the dark zone (DZ). 20X objective: scalebar 100 μm. D: Staining for DPEAAE, CD68 and CXCL13 demonstrates proximity of DPEAAE staining with CXCL13 staining within the GC-LZ. CD68 marks tingible body macrophages within the germinal center. A subset of LZ CD68^+^ macrophages is CXCL13^+^ (white arrows) but the majority of the CXCL13 signal derives from non-macrophage populations. 40X objective: scalebar 50 μm, inset scalebar 20 μm. E: Human tumor stratification as VPP, VPW-VCAN^lo^, and VPW-VCAN^hi^ followed established criteria that were prospectively validated ^36,41^. F: Staining for DPEAAE, CD68 and CXCL13 in a non-small cell lung cancer tissue microarray (NSCLC TMA) ^39^ is shown to have a stromal distribution. Both CD68^+^CXCL13^+^ (blue arrows) and CD68-CXCL13^+^ (white arrows) cells are demonstrated. T, tumor core. S, stroma. 40X objective: scalebar 50 μm, inset scalebar 20 μm. G: Immunohistochemical staining for XCR1, CD8 and CD4 in NSCLC TMA ^39^ was analyzed as detailed in Materials and Methods to enumerate immune cell populations and frequency of triad interactions. Tumors were categorized as VPP or VPW per established criteria based on VCAN and DPEAAE staining ^36,41^. 40X objective. H: Density of individual immune cell types per HPF as well as XCR1:CD4:CD8 triad configurations compared between VPP (n=20) and VPW (n=70) tumors in the NSCLC TMA ^39^. I: Proportion of tumor cores in the NSCLC TMA characterized by CD20+ infiltrate/aggregates (versus cases characterized by lack of CD20+ staining or few individual dispersed CD20+ cells) in VPW (n=71) vs. VPP (n=19) tumors. J: Workflow for IF analysis of TME in animal models. K: Animal models were engineered to replicate VPP and VPW human cohorts (K, top). Endogenous *Vcan* production was measured by RT-PCR relative to endogenous *Sdha* expression (K, bottom left). Endogenous Vcan proteolysis was interrogated by DPEAAE probing of total tumor lysates (K, bottom right). B16F10 is provided as a previously published endogenous Vcan production negative control ^107^. L: Workflow for IF and volumetric microscopy of animal model tumors after staining for CD11c (DC), CD4 and CD8. For each candidate heterologous cell pair, a line was drawn across the apposed cell borders and the separation between the two marker intensity maxima was recorded; because CD11c, CD4 and CD8a are surface antigens, this peak-to-peak separation reports the distance between the apposed plasma membranes. Representative profiles are shown for CD11c–CD8a (3.1 µm), CD11c–CD4 (0.4 µm) and CD4–CD8a (2.8 µm). Immune triad calling required all three pairwise membrane-to-membrane distances to be equal to or less than 5 µm. Representative Imaris images are shown for a triad, and two dyads where CD11c^+^ are apposed to a single lymphocyte partner. Where the nearest CD11c⁺ membrane maximum corresponded to a dendritic process rather than the cell body, the contact was retained, since DC–T cell interactions are initiated at processes. 10X objective: scalebar 100 µm, 63X objective: scalebar 20 µm. M: Top, IF microscopy delineates CD11c+, CD8+ and CD4+ cells in stromal neighborhoods for each model. Heterologous three-cell aggregates shown in white arrows. Bottom, density of immune triads in each model (defined as number of CD11c^+^ in triad normalized to total CD11c⁺ counted in the same manner and extrapolated to 1 mm³ volume). 63X objective: scalebar 20 μm. Data are representative as mean ± SEM. Statistics were performed using unpaired Student’s t test. Statistical significance is provided as the exact p value or denoted by an asterisk, *p < 0.05; **p < 0.01; ***p < 0.001, ****p < 0.0001. p values less than 0.05 were considered statistically significant.

### Cell quantification and proximity analysis

Individual raw z-stacks were imported into the Indica Labs HALO image analysis platform (v4.3). Nuclear segmentation was performed using DAPI counterstaining to define individual cell boundaries. Fluorescence intensity within each segmented cell was quantified across all channels, and positive cells were identified using intensity-based thresholding, with threshold values held constant across all samples. The algorithm classified cells within the XY plane into discrete phenotypic populations according to their fluorescence intensity profiles, enabling quantification of cell counts for each phenotype. Spatial relationships between phenotypes were assessed using the HALO Proximity Analysis module within the XY plane, quantifying the number of cells of a given phenotype located within defined radial distance bins (5 µm increments) up to a maximum distance of 100 µm from cells of a reference phenotype (CD11c). This proximity analysis is two-dimensional and is reported separately from the volumetric triad quantification above. For cell quantification within tumor stroma, stromal boundaries were defined manually as regions of interest and stromal area was quantified by HALO.

### Therapeutic efficacy studies

Recipient syngeneic mice were inoculated with 5×10^5 cells of either LLC-EV, LLC-Vkine, MC38-EV, MC38-Vkine, CT26-EV, or CT26-Vkine. Anti-PD-L1, anti-CTLA-4, anti-IL4 antibodies or isotype control (control IgG, Sigma-Aldrich) were administered at 100µg in 100µL PBS per dose through the intraperitoneal route (IP), starting on Day 3 post-inoculation and every three days thereafter. Tumor growth was monitored using a digital caliper. Tumor measurements were taken every three days post-inoculation. Mice were considered to reach survival endpoints when they were in clinical distress above IACUC-mandated threshold, when their tumors reached 20 mm in any dimension, or when they were found deceased. For tumor rechallenge studies, mice were allowed to rest for at least 45 days following clearance of the original, palpable tumor. 5×10^5 cells of the native, unmanipulated cell line (LLC, MC38, CT26) were injected into the opposite flank of mice and their tumor burdens were subsequently monitored in the same manner for demonstration of immune memory.

### LNP-mRNA formulation and characterization

Thy1.1 and Vkine-myc mRNA were synthesized *in vitro* using the Takara IVTpro mRNA synthesis system from the DNA templates described above. Lipid mixtures for LNPs consisted of the ionizable lipid SM-102 (Cayman Chemical, #2089251-47-6), 1,2-DSPC (Avanti Polar Lipids, #816-94-4), cholesterol (Sigma, #C8667), and DMG-PEG2000 (Avanti Polar Lipids, #160743-62-4), dissolved in pure ethanol at a molar ratio of 50:10:38.5:1.5, respectively, and a total lipid molar concentration of 10 mM. This “organic phase” was mixed with an aqueous phase of 50 mM sodium acetate buffer containing mRNA at a 3:1 aqueous to organic ratio, using the PreciGenome Nanogenerator Flex-M particle synthesis system. For the synthesis, the system’s total flow rate (TFR) and flow rate ratio (FRR) were set to 3 mL/min and 3:1, respectively. The resultant mRNA-LNP formulations were dialyzed against PBS using the 10K MWCO Slide-A-Lyzer G3 Dialysis Cassettes (Thermo Scientific, #A52971), concentrated with Amicon Ultra Centrifugal filters (Merck Millipore), and passed through a 0.2µm filter. mRNA-LNPs were then tested for particle size, polydispersity, and zeta potential using the Zetasizer Nano (Malvern Panalytical), and tested for concentration and encapsulation efficiency (typically 85%-92%) using the Quant-it Ribogreen assay (Invitrogen, # R11490). Formulations were then diluted in PBS to a concentration of 200 ng mRNA/µL.

### *In vivo* LNP-mRNA administration and monitoring

Recipient WT C57BL/6J and/or *Batf3* -/- mice were inoculated with 2.5 x 10^5 MC38 cells (10 mice per arm). One week post tumor inoculation, 5 µg of LNP-formulated mRNA (Thy1.1 or Vkine-myc) was delivered intratumorally. mRNA treatments were administered on Days 7, 9, and 11 post-inoculation. For experiments evaluating therapeutic synergy, mice were also treated with either 200 µg of either isotype or anti-PD-L1 antibody in 200 µL PBS, on Days 9 and 11 post-inoculation through the intraperitoneal route (IP). For T-cell depletion experiments, 150 µg of anti-CD8 or isotype was administered one day prior to treatment and every three days afterwards. Tumor growth was monitored using a digital caliper. Measurements were taken every three days post-inoculation. Mice were considered to reach survival endpoints when they were in clinical distress above IACUC-mandated threshold, when their tumors reached 20 mm in any dimension, or when they were found deceased. Mice that were tumor-free following treatment were monitored for the absence of a palpable tumor for at least 30 days, then they were euthanized and their spleens were harvested. Pan-T splenocytes were isolated using the Pan-T Cell Isolation Kit (Miltenyi Biotec, #130-095-130). 1.5×10^6 pan-T cells were adoptively transferred to naive recipients via retroorbital inoculation and 2.5 x 10^5 MC38 cells were subcutaneously inoculated the following day. Tumor burden was monitored in the same manner as mentioned above.

### Recombinant Vkine (rVkine) production and purification

HEK293T-Vkine producer cells were seeded in T-175 cell culture flasks and cultured in DMEM 10% FBS media. At 75 to 80% confluency, cell media were changed to DMEM 1% FBS and collected after 48 hours of incubation. Collected supernatant was centrifuged to remove debris and filtered with a 0.22 μm PES syringe filter. Protein purification was performed using a nickel-based immobilized metal affinity chromatography (IMAC) (Bio-rad) equipped with UV, conductivity, and pH monitors controlled via ChromLab software. Cell media containing rVkine was loaded onto 5 mL His Trap^TM^ HP column (Cytiva, #17524801) at a flow rate 2 mL/min. Chromatography system and column were washed with 20% ethanol and 1 M NaOH and rinsed with DMEM medium. Unbound material was removed by washing with DMEM containing 10 mM imidazole until UV absorbance (280 nm) returned to baseline. Bound protein was eluted using elution buffer (DMEM containing 500 mM imidazole) into 2.5 mL fractions which were collected.

SDS-PAGE and Coomassie staining of the gel (GelCode™ Blue Safe Protein Stain, Thermo Fisher, # 24594) was used to visualize the total protein content in each fraction. Fractions containing eluted protein were concentrated using Sartorius Vivaspin 20, 10,000 MWCO PES concentrator (#VS2001). During this step, buffer exchange to PBS was performed following manufacturer recommendations. Endotoxin assay was performed using Genscript ToxinSensor Chromogenic LAL Endotoxin Assay Kit (#L00350C). Endotoxin was removed using Pierce™ High Capacity Endotoxin Removal Spin Columns (Thermo Fisher, #88274). rVkine identity was confirmed by western blot using c-myc, DPEAAE and G1-domain (VCAN) antibodies and concentration was determined by DPEAAE ELISA, as detailed below.

### Vkine ELISA

rVkine ELISA was performed with modifications from a published protocol ^51^. DPEAAE antibody was used to coat Biolegend 96-well uncoated Nunc™ MaxiSorp™ ELISA plates (Biolegend, #423501) in coating buffer (Na_2_CO_3_ 15 mM and NaHCO_3_ 35mM pH 9.6) overnight. The rest of the protocol was performed as detailed in the reference above.

### rVkine *in vivo* administration

C57BL/6J mice were inoculated with 5 x 10^5 MC38 cells. On Days 7, 10 and 13 post-inoculation, incremental doses (50, 75 or 125 μg) of rVkine were delivered intratumorally in a 100 μL volume. Mice were also treated with either 200 µg of either isotype (IgG control) or anti-PD-L1 antibody in 200 µL PBS on Days 9 and 11 post-inoculation through the intraperitoneal route (IP). Tumor growth was monitored using a digital caliper. Measurements were taken every three days post-inoculation. Mice were considered to reach survival endpoints when they were in clinical distress above IACUC-mandated threshold, when their tumors reached 20 mm in any dimension, or when they were found deceased.

### RNA isolation and Real-Time PCR

RNA was isolated using QIAGEN RNeasy Mini Kit and cDNA was synthesized using the iScript Reverse Transcription Supermix (Biorad, #1708840). Quantitative real-time (qRT-PCR) analysis was performed using SsoAdvanced Universal SYBR Green Supermix (Biorad #1725272) according to the manufacturer’s instructions on an CFX96 Touch Real Time PCR detection (Bio-rad, # 1845097) using the relative standard curve method. PCR conditions were 2 min at 50°C, 10 min at 95°C followed by 40 2-step cycles of 15 s at 95°C and 1 min at 60°C. QuantiTec Primers (Qiagen), for *Vcan* as well as *Sdha* for normalization control, were used to assess relative gene expression.

### scRNAseq library preparation and sequencing

Single-cell suspensions were prepared from CT26-EV (n= 3 mice), CT26-Vkine (n= 3 mice), MC38-EV (n= 4 mice), MC38-Vkine (n= 4 mice), LLC-EV (n= 4 mice), and LLC-Vkine (n= 3 mice) tumors harvested on Day 21 post-subcutaneous inoculation as described in the tissue processing section. Dissociated tumors were stained with the viability probe, Ghost Dye 780 for 10 minutes. Cells were washed, Fc-blocked and stained with anti-CD45 antibody. Around 2×10^5 live CD45^+^ cells were sorted using FACSymphony S6 cell sorter (BD Biosciences). FACS-sorted single cell suspensions were washed with ice cold FACS buffer and resuspended in PBS containing 10% FBS in a concentration of 1400 cells/μL. Single cells were immediately processed for individual transcriptomic profiling using Chromium GEM-X Single Cell 3’ v4 Gene Assay (10x Genomics, #CG000731). A pool of 3.6 M barcodes was used to index around 16,000 cells individually through partitioning cells in nanoliter scale Gel Beads-in-emulsion (GEM). Barcoded and full-length cDNAs from polyadenylated-mRNAs were amplified via PCR to generate sufficient mass for library construction. The final libraries contained specific sequences for paired-end Illumina sequencing. Sequencing was carried out using sequencer-reader library Illumina NovaSeq 6000. Sequences are publicly accessible through GEO (GSE348233).

### Analysis of scRNAseq data

#### Pre-processing

Raw sequencing data were demultiplexed and aligned to the mouse reference genome (mm10) to generate gene-barcode count matrices using Cell Ranger v7 ^52^. Because CT26, MC38, and LLC tumors arise in different genetic backgrounds and exhibit distinct tumor microenvironments, we chose to process each dataset independently rather than integrate to avoid batch-correction artifacts that could obscure biologically meaningful differences between models. Count matrices were imported into Seurat v5 ^53^ for quality control, normalization, and downstream analysis. Cells with less than 200 genes, greater than 10% mitochondrial reads, or with total UMI counts below the 1^st^ or above the 99^th^ percentile were excluded from downstream analysis. RNA expression data were normalized and scaled, and the top 3,000 variable genes were used for principal component analysis. For each dataset, the number of principal components was selected at the elbow of the scree plot and corrected for potential batch effects using Harmony (CT26) or CCA (MC38, LLC) on the sample ID. Batch-corrected components were then used to generate k-nearest neighbor (kNN) graphs and uniform manifold approximation and projection (UMAP) embeddings (CT26: PCs = 30, resolution = 0.8; MC38: PCs = 20, resolution = 0.5; LLC: PCs = 30; resolution = 0.5). Major immune lineages were identified by *de novo* markers (Supp. Table S2) and expression of well-established immune lineage markers: T cell (*Cd3d, Themis, Trac*), B cell (*Cd79a, Pax5, Ebf1*), NK cell (*Ncr1, Klrb1c, Car2*), dendritic cell (*Itgax, Flt3, H2-Ab1*), monocyte/macrophage (*Adgre1, Msr1, Lyz2*), neutrophil (*Csf3r, Cxcr2, S100a9),* and mast cell (*Mcpt4*, *Cma1*), allowing for more precise cell type identification during subsequent sub-clustering. Individual Seurat objects were created after subsetting cell types of interest.

#### Sub-clustering of T cells

Established markers were used to identify and subset T cells (Cd3d, Themis, Trac). PCA, UMAP, and Louvain clustering steps were performed as above. T cell sub-populations were annotated by reference mapping and label transfer using the ProjecTILs mouse TIL atlas v1 ^54^. Clusters mapping to multiple predicted subpopulations were iteratively subclustered until ProjecTILs assigned a single predicted cell type to each resulting cluster. If a single identity could not be resolved, the cluster was designated as unresolved. This process was repeated across a range of prediction confidence score thresholds, and cell identity assignments remained consistent. Final annotations were validated by confirming that the top 25 *de novo* marker genes identified for each cluster -- using FindAllMarkers (two-sided Wilcoxon rank-sum test with Benjamini-Hochberg correction; min.pct = 0.1; min.diff.pct = 0.25) -- were consistent with the identity of the annotated T cell subpopulation.

Sub-clustering of Dendritic cells.

Dendritic cells were identified and subset based on expression of well-established markers (*Itgax*, *Flt3*, *H2-Ab1*). PCA, UMAP, and Louvain clustering steps were performed as above. Dendritic cell subpopulations were annotated by module scoring using subtype-specific marker gene sets ^55^: cDC1 (*Xcr1, Clec9a, Cadm1*); cDC2 (*Cd209a, H2-DMb2, Sirpa, Tlr2, Myd88*); mregDC (*Ccr7, Il12b, Fscn1, Cd200, Cd274*); pDC (*Siglech, Tlr7, Tlr9, Ifna1, Ifnb1, Il6, Tnf*); and moDC (*Adgre1, Cd69, Mertk, Fcgr1*). Final annotations were assigned based on module scores and canonical gene expression, in addition to verifying the top 25 *de novo* marker genes for each cluster were consistent with the assigned identity. Conventional and inflammatory DC sub-populations (cDC1, cDC2, moDC, and mregDC) were retained for downstream analysis.

#### Differential gene expression analysis

Differential gene expression between conditions was assessed using FindMarkers (two-sided Wilcoxon rank-sum test with Benjamini-Hochberg correction) on the RNA assay. Genes with absolute log2 fold-change ≥ 0.25, adjusted p-value < 0.05 and expression detected in at least 10% cells were considered significantly differentially expressed. GSEA ^56^ was performed on the complete ranked gene list using clusterProfiler against Gene Ontology Biological Processes and Hallmark mouse gene sets retrieved via msigdbr (v25.1.1), with genes ranked by sign(log2FC) ×-log10(p-value). Gene sets containing between 15 and 500 genes were retained for testing. Module scores were calculated using Seurat’s AddModuleScore or UCell ^57^.

#### Cell-cell communication analysis

Ligand/receptor interactions were inferred in CellChat using the mouse CellChatDB v2 database ^58^. Interaction probabilities were computed using the trimean method, weighted by the proportion of cells expressing each ligand/receptor within a population, and aggregated at the signaling pathway level. Interactions involving fewer than 10 cells were excluded. Objects were created independently for EV and Vkine and subsequently merged using the mergeCellChat function for cross-condition comparison.

### Reanalysis of publicly available scRNAseq data from NCT02837263

#### Pre-processing

Publicly available scRNA-seq data ^41^ profiling peripheral blood mononuclear cells collected at baseline and following stereotactic body radiation therapy, pembrolizumab, and surgical resection were obtained through NCBI GEO (accession number GSE316301) for independent analysis. Count matrices and metadata were used to generate Seurat objects. Cells with fewer than 200 genes, more than 10% mitochondrial reads, more than 40% ribosomal reads, or with total UMI counts below the 1st or above the 99th percentile were excluded. RNA expression data were log-normalized and scaled, and the top 4,000 variable genes were used for principal component analysis. Harmony-corrected components were then used to generate uniform manifold approximation and projection (UMAP) embeddings. Cell type annotations were derived using *de novo* markers and canonical gene expression ^59^ and validated by label transfer and reference mapping to the Azimuth human PBMC reference ^60^.

#### Th7R identification and functional analysis

The CD4^+^ T cell compartment was subset and reclustered. A previously reported gene signature derived from sorted Th7R, Th1, and Th17 cells ^59^ was applied to SELLlow CD4^+^ clusters. For each cluster, UCell was used to compute per-cell enrichment scores for the Th7R, Th1, and Th17 signatures using their respective positive and negative marker genes, and the proportion of cells scoring highly for each signature was calculated per cluster. Clusters were assigned the identity of their dominant signature; clusters in which two or more signatures scored highly, or none scored highly, were annotated as unresolved. Differential gene expression and transcription factor activity were compared between VPP and VPW patients at baseline and post-treatment using a two-sided Wilcoxon rank-sum test with Benjamini-Hochberg correction. Differential gene expression was assessed using FindMarkers on the RNA assay, and transcription factor (TF) activity was inferred using the univariate linear model method implemented in decoupleR ^61^ with regulons from the collecTRI database ^62^.

#### Tpex identification

The CD8^+^ T cell compartment was subset and reclustered. Clusters were examined for expression of GZMK and GZMB to distinguish circulating precursor-exhausted (Tpex), exhausted (Tex), and cytotoxic (CTL) CD8^+^ T cells, consistent with Takei et al ^59^. GZMK^+^ GZMB-clusters were further evaluated for expression of PDCD1 (PD-1), CXCR5, CCR7, IL7R, and TCF7 (adapted from ^59^). Differential gene expression between VPP and VPW patients was performed using FindMarkers as described above.

### Statistical analysis

Statistical analysis was performed using GraphPad Prism software (GraphPad version 9.0.0). Statistical significance was determined using unpaired two-tailed Student’s t-test, unpaired non-parametric Mann-Whitney test or Wilcoxon rank sum test with continuity correction as indicated in figure legends. The log rank (Mantel-Cox) test was used to determine statistical significance for overall survival *in vivo*. Data are shown as mean ± SEM. Significance was indicated with asterisks *p < 0.05; **p < 0.01; ***p < 0.001, ****p < 0.0001. p values less than 0.05 were considered statistically significant.

### Graphics

Graphics and diagrams were created using BioRender.

## RESULTS

### Constitutive VCAN proteolysis in human germinal centers localizes within CXCL13-rich light zone gradients

Regulated proteolysis of the large matrix proteoglycan versican (VCAN) by ADAMTS versicanases (ADAMTS-1,4,5,9,15 and 20) at position 441-442 of the V1 isoform releases a bioactive fragment (matrikine), Vkine (Fig. 1A). The cleavage event unveils the neoepitope DPEAAE at the C-terminal end of Vkine which can be immunochemically detected ^63^. We and others have demonstrated critical roles for Vkine in antigen presentation and tumor immunosurveillance when released in the peritumoral stroma ^35,36,63^, in addition to known roles in development, wound healing and non-tumoral inflammation ^29,30,43,64^. However, a role for Vkine in normal lymphoid tissue architecture and lymphoid neogenesis has not been previously reported. We used immunohistochemical staining for DPEAAE to detect robust VCAN proteolysis in germinal centers of human normal tonsillar tissue (Fig. 1B). DPEAAE appeared to stain polar ends of germinal center structures, suggesting that VCAN proteolysis and Vkine release may be restricted to either the dark zone (DZ) or light zone (LZ). To determine Vkine’s anatomical distribution, we used staining for Ki67, a marker delineating active proliferation of cycling DZ centroblasts. Using this criterion, DPEAAE staining appeared to be limited to the LZ (Fig. 1C and Supp. Fig. S1A). Staining for CD23, a marker delineating the extent of the LZ follicular dendritic cell (FDC) network, confirmed co-localization of DPEAAE in the LZ (Supp. Fig. S1B). Indeed, FDCs have been reported to express ADAMTS versicanases both constitutively and inducibly ^65^. Staining for intact VCAN demonstrated individual producer cells located mostly in the LZ and marginal sinus that also co-stained for CD68, identifying them as LZ tingible body macrophages (Supp. Fig. S1C/D).

CXCL13 gradients define and position the GC light zone ^66^. We sought to determine whether CXCL13 gradients corresponded with the extent of constitutive VCAN proteolysis in the GC. Indeed, we found that VCAN proteolysis and CXCL13 gradients precisely overlapped within tonsillar LZ (Fig. 1D and Supp. Fig. S1E). Within the LZ, we detected a subset of CXCL13^+^ CD68^+^ macrophages, however most CXCL13 appeared to derive from CD68^neg^ cells, consistent with known secretion of the chemokine by follicular dendritic cells and T follicular helper cells ^67^ (Fig. 1D and Supp. Fig. S1E). LZ-restriction of VCAN proteolysis was consistent across human secondary lymphoid tissue locations, with a representative pattern from a normal lymph node GC shown in Supp. Fig. S1F.

### VCAN proteolysis in human tumors co-localizes with CXCL13^+^ stromal niches

We then sought to determine whether VCAN proteolysis overlaps with CXCL13 gradients in tumors. VCAN proteolysis in tumors is inflammation-driven, dynamic and inducible (not constitutive) and mainly localized to the stroma ^36,39^. To determine whether VCAN proteolysis overlaps with CXCL13 activity, we stratified a previously reported TMA of human non-small cell lung cancer ^39^ into 3 cohorts using Vkine/VCAN staining criteria previously used to classify the independent clinical trial cohort ^41^ (depicted in Fig. 1E). The most immune active VCAN-proteolysis-predominant (VPP) group included tumors with *both* an endogenously active VCAN proteolysis pathway *and* low levels of residual, unprocessed (unproteolyzed) parental VCAN (DPEAAE 2-3+ AND VCAN staining 0-1+). These patients (40% of overall in ^41^) were shown to respond most robustly to pembrolizumab ICI and demonstrated the best survival outcomes. Two other groups (collectively termed VCAN-proteolysis-weak, VPW) fared clinically much worse. The VPW-VCAN^lo^ subgroup constituted approximately 50% of the overall cohort in the clinical trial ^41^ and were relatively resistant to single-agent ICI. These patients have low/undetectable rates of VCAN proteolysis and low levels of unproteolyzed VCAN, i.e., low pathway activity overall. Patients whose tumors demonstrated excess unproteolyzed VCAN (VPW-VCAN^hi^) were a relatively small group (app. 10%) with particularly adverse outcomes. In VPW-VCAN^hi^ patients, VCAN proteolysis occurs at rates that are not sufficient to deplete accumulated parental VCAN, therefore the VCAN:Vkine ratio is inadequate to unmask the beneficial effects of the released matrikine. In these VPW-VCAN^hi^ tumors, parental VCAN actively antagonizes the immunostimulatory effects of released Vkine ^32^.

In VPP patient samples, VCAN proteolysis was observed in the tumor stroma in close proximity with CXCL13^+^ cells (Fig. 1F and summarized in Supp. Fig. S1G). CXCL13^+^ macrophages, previously reported to be a major source of CXCL13 in inflammatory lymphoid neogenesis settings ^68^ were detected in these areas (Fig. 1F, white arrows). However, CD68^neg^ CXCL13^+^ cells were predominant (Fig. 1F, green arrows), consistent with prior reports demonstrating CXCL13 derivation from infiltrating Tfh cells or follicular dendritic cells (FDC) in nascent TLS ^69,70^. These data suggest that VCAN proteolysis co-localizes with CXCL13 niches and may synergize with CXCL13 chemotactic gradients to promote TLS assembly.

### VCAN proteolysis is associated with prominent DC:CD4^+^ T:CD8^+^ T immune triad formation in human tumors

Tripartite antigen-dependent cross-talk in physical “triads” consisting of a DC, a CD4^+^ T cell and a CD8^+^ T cell constitutes an early event in TLS organization, is required for TLS maintenance and is critical for effector T cell differentiation and response to ICI ^6,50,71–76^. To determine whether VCAN proteolysis is associated with enhanced rates of immune triad formation in human samples, we used immunohistochemistry to detect XCR1, a marker of antigen-cross-presenting cDC1, as well as CD4 and CD8 on the lung cancer samples (Fig. 1G). Following staining, the tissue microarray samples were scanned and processed as described in the Materials and Methods.

The VPP phenotype demonstrated a significantly higher number of intratumoral CD8^+^ T cells, suggesting T-cell inflammation, as previously reported (Fig. 1H). A higher proportion of stromal XCR1^+^ DC was also seen, consistent with prior reported activities of stromal Vkine in expanding cDC1 through enhanced survival ^39^. This analysis also demonstrated a higher number of both stromal and intraepithelial CD4^+^T cells in VPP tumors, reminiscent of the independent clinical trial VPP cohort ^41^. A higher number of immune triads consisting of XCR1^+^ DC, CD4^+^ and CD8^+^ T cells was detected in VPP tumors compared to VPW tumors (Fig. 1H). Immune triads are an early and necessary step in TLS organization, but the structures they seed are defined by a B cell compartment. We therefore asked whether VCAN proteolysis status was associated with B cell infiltration in the same tumors. CD20+ infiltrate/aggregates were present in 47.4% of VPP tumors (9 of 19) compared with 22.5% of VPW tumors (16 of 71; two-sided Fisher’s exact test, p = 0.044; OR 3.09) (Fig. 1I). The association tracked the extent of proteolysis rather than its mere presence. Across ascending strata of stromal Vkine intensity, the proportion of tumors containing CD20+ infiltrate/aggregates rose from 6.2% (1 of 16) at intensity 0, to 20.6% (7 of 34) at intensity 1, to 41.5% (17 of 41) at intensity 2–3 (exact Cochran–Armitage test for trend, p = 0.0047) (Supp. Fig. S1H). Notably, in tumors with no detectable stromal Vkine, B cell infiltrate/aggregates were almost entirely absent. These data indicate that VCAN proteolysis status in human tumors is associated not only with the tricellular units that initiate lymphoid organization, but with the B cell compartment that defines its product.

### The product of VCAN proteolysis, Vkine, suffices for tumor immune triad formation

We hypothesized that VCAN proteolysis acts through the released matrikine, Vkine, in promoting immune triads. To test this hypothesis, we engineered a series of experimental tumor models replicating each of the human patient cohorts in the clinical trial ^41^. To replicate VPW-VCAN^lo^ and VPP conditions, we ectopically expressed Vkine in two models of colorectal cancer (CT26, MC38) with undetectable tumor cell-derived VCAN and low VCAN expression in implanted subcutaneous tumors (Fig. 1J/K). Thus CT26-Vkine and MC38-Vkine phenocopy VPP tumors and CT26-EV and MC38-EV phenocopy the corresponding VPW-VCAN^lo^ tumors.

To model VPW-VCAN^hi^ tumors, we utilized the Lewis Lung Carcinoma (LLC) model which robustly overexpresses VCAN from the tumor cells themselves but also tumor-infiltrating leukocytes (Fig. 1K). LLC-Vkine and LLC-EV models thus phenocopy human VPW-VCAN^hi^ tumors, with or without VCAN proteolysis, respectively. We confirmed that subcutaneous tumors from all three models lack endogenous VCAN proteolysis that could be confounding our modeling and predictions (Fig. 1J/K). Moreover, we confirmed that the expression levels of ectopic Vkine were comparable between each engineered cell line (Supp. Fig. S1I).

CT26-EV, CT26-Vkine, MC38-EV, MC38-Vkine, LLC-EV and LLC-Vkine tumors were implanted in immunocompetent mice and harvested on Day 21 post implantation. Tumors were fixed and sectioned in 25μm sections and subjected to IF staining using anti-CD11c, anti-CD4 and anti-CD8 antibodies. Across all three models, Vkine promoted broad immune infiltration which was more pronounced for CD8^+^ T cells in the CT26 model and CD4^+^ T cells in the MC38 model (Supp. Fig. S1J). Ectopic expression of Vkine resulted in increased CD11c^+^ density in the more immunogenic models by IF (Supp. Fig. S1J).

To enumerate intratumoral triads in three dimensions, volumetric immunofluorescence imaging was performed using confocal microscopy. For each field of view, XY plane was conserved between acquisitions and a z-stack comprising 25 optical sections was collected at a step size of 1 µm using sequential laser excitation at 405, 488, 561 and 639 nm wavelengths. To reconstruct and visualize 3D objects, raw z-stack images were imported to Imaris and analyzed as described in Materials and Methods. Triad formation was defined as the co-localization of an individual CD11c⁺ cell with both a CD8⁺ cell and a CD4⁺ cell, such that each pairwise inter-membrane distance (CD11c⁺–CD8⁺, CD11c⁺–CD4⁺, and CD8⁺–CD4⁺) did not exceed 5 µm. The number of CD11c^+^ cells in triads identified within a 25 µm z-depth (XY-plane analysis) was normalized to total CD11c⁺ counted in the same manner and extrapolated to 1 mm³ volume (Fig. 1L). This analysis demonstrated that Vkine promoted immune triads in all 3 models irrespective of the genetic background and the presence or absence of excess unproteolyzed VCAN (Fig. 1M). Thus Vkine, the product of VCAN proteolysis, suffices for immune triad formation across variable immunogenicity TMEs.

### Vkine delays tumor growth in a cDC1-dependent manner

Ectopic expression of Vkine by tumor cells *in vitro* does not result in appreciable growth or proliferation differences in culture, with EV tumor cells demonstrating *in vitro* kinetics comparable to their Vkine-expressing counterparts (Supp. Fig. S2A). By contrast, a profound growth defect of Vkine-expressing tumors is observed in immunocompetent hosts in the immunogenic CT26 and MC38 models engineered to express Vkine ectopically (Fig. 2A/B). As previously demonstrated, Vkine has no “monotherapy” activity in the VCAN-replete LLC context (Fig. 2C), consistent with human clinical trial data for the ICI-resistant VPW-VCAN^hi^ cohort of patients ^41^.

**Fig. 2.**
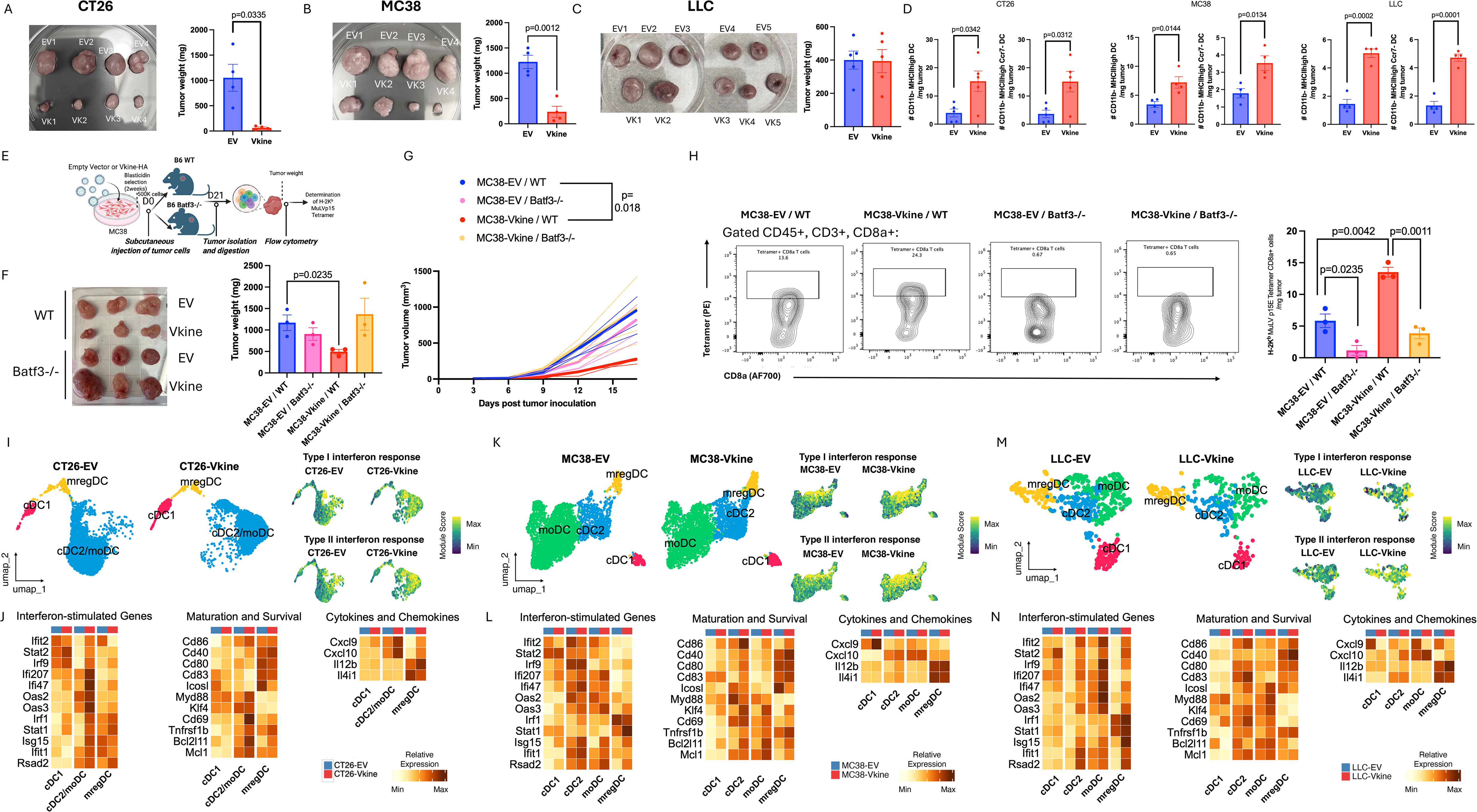
Vkine expands tumor cDC1 and activates CD11b^+^ DC. Tumor volume and weight endpoints on Day 21 post-implantation demonstrate a growth defect of CT26-Vkine (A) and MC38-Vkine tumors (B) compared to EV counterparts but no significant difference between LLC-Vkine tumors and LLC-EV controls (C). Expansion of total CD11b^neg^ DC (D, left panels) and non-migratory CD11b^neg^Ccr7^neg^ DC (cDC1) (D, right panels) across experimental models. E: MC38-EV and MC38-Vkine tumors were implanted in WT or *Batf3*-null syngeneic recipients. D21 endpoints shown in gross morphology (F, left) and tumor weights (F, right). G: Individual tumor growth spider plots (thin lines) and means (thick lines) shown for tumor growth rates of MC38-EV and MC38-Vkine tumors implanted in WT or *Batf3*-null syngeneic recipients (n=3 per arm). H: Tetramer staining for MC38-intrinsic MuLV-derived tumor antigen (p15E) reveals that an increase in tumor-antigen-specific CD8^+^ T cells in MC38-Vkine conditions compared to-EV (H, left two flow cytometry panels) is abrogated by *Batf3* loss (H, right two flow cytometry panels). Absolute density of tetramer^+^CD8^+^ T cells per unit tumor mass reveals lack of increase of tumor antigen-specific T cells in *Batf3*-null recipients of MC38-Vkine tumors (H, right). Uniform manifold approximation and projection (UMAP) plots of scRNAseq of DC subsets reveal a dramatic shift in transcriptomic profiles of cDC2 (a CD11b^+^ DC subset) in CT26-Vkine compared to CT26-EV as well as an increase in cDC1 density (I, left panels). Feature plots showing module scores for Type I interferon response and Type II interferon response gene signatures in EV- and Vkine-conditions (I, right panels). Gene sets were obtained from Gene Ontology Biological Processes. J: Heatmaps showing relative gene expression for each DC cluster reveal induction of interferon-stimulated genes (ISG), maturation and survival genes as well as cytokines and chemokines. Corresponding patterns for MC38 (K/L) and LLC (M/N) are shown. Panels A, B, C, D, F, and H designate mean ± SEM and were analyzed using unpaired Student’s t test. Growth curves in G were compared using Mann-Whitney test. Statistical significance is provided as the exact p value or denoted by an asterisk, *p < 0.05; **p < 0.01; ***p < 0.001, ****p < 0.0001. p values less than 0.05 were considered statistically significant.

We hypothesized that Vkine exerts anti-tumor activity via cDC1 as this antigen-presenting-cell subtype has been shown to have critical roles in peri-tumoral stimulation of incoming antigen-experienced T cells as well as TLS formation and maintenance ^6,77^. We have previously demonstrated that Vkine instigates an NK-dependent loop that promotes survival of cDC1 ^39^. We specifically sought to determine whether Vkine regulated the tumor-resident Ccr7^neg^ cDC1 fraction in our models, based on prior evidence for critical roles of this fraction in peritumoral restimulation of LN-primed incoming T cells ^77^. To do so, we specifically enumerated CD11b^neg^ DC and CD11b^neg^Ccr7^neg^ DC based on our previously published flow-cytometry-based protocol ^78^. Indeed, across all three models, Vkine promoted expansion of CD11b^neg^Ccr7^neg^ intratumoral DC (Fig. 2D). Additional lines of evidence have demonstrated a central role for Ccr7^+^ DC in immune triad formation ^6,73^. We detected a trend towards an expansion of the Ccr7^+^ migratory (mregDC) fraction (approximately 2-3 fold), however, small numbers of intratumoral Ccr7^+^ DC precluded statistical significance for this DC subset using flow cytometry (Supp. Fig. S2B).

We then asked whether cDC1 were required for Vkine’s anti-tumor activity. To this end, we implanted MC38-EV and MC38-Vkine cells in *Batf3*-/- hosts that lack endogenous cDC1 ^79^ (Supp. Fig. S2C). *Batf3* loss completely abrogated Vkine’s growth-inhibitory activity, suggesting that antigen cross-presenting cDC1 were essential to mediate Vkine’s effects (Fig. 2E/F/G). Moreover, *Batf3* loss abrogated tumor antigen-specific T cell responses facilitated by Vkine, as shown by tetramer staining of MC38 tumor-specific CD8^+^ T cells recognizing endogenous MuLV-derived p15E antigen ^80^ (Fig. 2H and Supp. Fig. S2D). Thus, *Batf3* loss abrogated the increase in tumor antigen-specific CD8^+^ T cells induced by Vkine.

### Vkine-replete TME promotes broad CD11b^+^ DC activation

To obtain granular insights into the immune TME composition of each VCAN status at a single-cell resolution, subcutaneous tumors from each model were derived and analyzed by scRNAseq. 500K cells were subcutaneously inoculated into the corresponding syngeneic recipients, BALB/c (CT26) or C57BL/6 (MC38, LLC) and tumors were harvested on Day 21. Single-cell suspensions were prepared from dissociated tumors, and CD45^+^ cells were purified via flow cytometry and subjected to scRNAseq (Chromium GEM-X Single Cell 3’ Gene Expression v4, 10x Genomics). Raw sequencing reads were aligned and quantified using Cell Ranger (10x Genomics) and processed in Seurat. After filtering and quality control, we recovered 35,382 cells from CT26-EV, 26,770 from CT26-Vkine, 33,707 from MC38-EV, 26,163 from MC38-Vkine, 14,871 from LLC-EV, and 9,204 from LLC-Vkine (Supp. Fig. S2E/F/G). Clusters were defined using an unsupervised approach (see Materials and Methods) and annotated by lineage based on *de novo* markers and canonical gene expression (Supp. Fig. S2H/I/J). This analysis identified six major immune lineages across the three models: T cell (*Cd3d, Themis, Trac*), B cell (*Cd79a, Pax5, Ebf1*), NK cell (*Ncr1, Klrb1c, Car2*), dendritic cell (DC) (*Itgax, Flt3, H2-Ab1*), monocyte/macrophage (*Adgre1, Msr1, Lyz2*), and neutrophil (*Csf3r, Cxcr2, S100a9),* alongside small populations of mast cells and non-immune cells (Supp. Fig. S2H/I/J and Supp. Table S2). As expected, *Ptprc* (Cd45) was broadly expressed across clusters, while *Vcan* was enriched in the monocyte/macrophage lineage (Supp. Fig. S2K-P).

DC were annotated based on expression of modules associated with published DC markers (Supp. Fig. S3A/G/M) and *de novo* gene expression (Supp. Fig. S3B/H/N). Across the three models, Vkine appeared to result in transcriptional changes consistent with broad DC activation (Supp. Fig. S3D/J/P). scRNAseq confirmed Vkine-induced cDC1 expansion in the more immune-infiltrated models CT26 and MC38 (Fig. 2I/K and Supp. Fig. S3C/I), consistent with the flow cytometry data and prior findings ^39^. cDC1 demonstrated enhancements in antigen presentation machinery (*Tap1, Tap2, MhcII*) as well as co-ordinate alterations in co-stimulatory (*Slamf7, Slamf8*) and co-inhibitory receptors (*Cd274, Pvr*)(Supp. Fig. S3E/K/Q). The most significant transcriptional profile changes, however, were detected in the CD11b^+^ DC (cDC2 and moDC) compartment. Vkine caused a dramatic shift in the transcriptional profile of CT26 cDC2/moDC with strong evidence of active Type I and Type II interferon incoming-(Fig. 2I) and outgoing signaling (Supp. Fig. S3F). Several ISGs were coordinately upregulated in cDC2/moDC in the presence of Vkine (Fig. 2J). Additional profile changes included an increase in activation (*Cd69*), co-stimulation (*Cd80*, *Cd83*, *Cd86*), survival (*Mcl1*, *Bcl2l11*) and IFN-dependent cytokine release (*Cxcl9*, *Cxcl10*)(Fig. 2J). Similar changes were also seen in the MC38-Vkine DC (Fig. 2L) and even the less immunogenic LLC-Vkine model (Fig. 2N). These data indicate that Vkine promotes profound alterations in CD11b^+^ DC transcriptional states, resulting in activation and enhanced antigen presentation. Regarding mregDC, despite the relative paucity of this crucial intratumoral population ^72^, analysis of transcriptional profiles promoted by Vkine indicates a redress in the balance between immunogenic (*Cd40, Il12b*) and tolerogenic (*Il4i1*) markers (Fig. 2J/L/N). Thus, Vkine may promote mregDC immunogenic repolarization.

### Tpex engagement by DC is enhanced in the presence of Vkine

Tpex have been the objects of intense scrutiny due to their role as the main mediators of response to checkpoint inhibitors ^81^. Tpex maintain their stem cell attributes through interactions with cDC1 in both reactive and tumor settings ^77,82^. We therefore asked whether enhanced tumor T cell infiltration and response to ICI in the presence of Vkine may be supported by an expanded Tpex reservoir and whether DC played a role in this process.

T cell annotations on the scRNAseq datasets were carried out based on the ProjecTILs mouse tumor-infiltrating T cell reference atlas ^54^ and canonical marker genes (Supp. Fig. 4A-L). The CT26 model proved the most useful platform in delineating Tpex-Tex dynamics in the presence of Vkine. In CT26, CD8^+^ T effector responses are known to be brisk and dynamic and result in rapid T cell exhaustion ^83^. Ectopic expression of Vkine in CT26 led to a significant expansion of the Tex and Tpex compartments (Supp. Fig. S4A). Tex in CT26-Vkine expressed a profound array of cytolytic molecules (*Gzmb*, *Prf1*) as well as activation/exhaustion markers (*Pdcd1*, *Lag3*, *Havcr2*) (Supp. Fig. S5A/B). In CT26, Vkine resulted in an overrepresentation of Tpex among *Tcf7*-expressing CD8^+^ T cells (Supp. Fig. S5C/D).

The CD8^+^ T cell dynamics in the MC38 model are known to be more blunted and sustained ^83^. Interestingly, Vkine in MC38 appeared to prevent terminal exhaustion with preservation of a Teff/mem phenotype even at a late stage of tumor evolution (Day 21) (Supp. Fig. S4E). These effector cells did express more cytolytic molecules in the presence of Vkine (Supp. Fig. S5E/F). A Tpex cluster was not clearly discernible by scRNAseq in this model under the conditions utilized (Supp. Fig. S5G/H). T cells represented a smaller proportion of CD45^+^ cells in LLC compared to CT26 or MC38, consistent with lower immunogenicity of this model (Supp. Fig. S4I and Supp. Fig. S2G) and Vkine demonstrated a minimal effect in T cell activation or expansion, reminiscent of VPW-VCAN^hi^ patients (Supp. Fig. S5I/J).

Consistent with the scRNAseq profiles, the CT26 model was the most informative in delineating Tpex spatial dynamics by IF analysis. Vkine was associated with an increase in CD8^+^Tcf1^+^ T cells, mostly located in the tumor periphery (Fig. 3A). HALO proximity analysis of IF images demonstrated that physical interactions between DC and CD8^+^Tcf1^+^ T cells were enhanced in the presence of Vkine, globally in the less than 30μm range (which indicates proximity) but specifically within the 5-10 μm range that suggests physical contact (Fig. 3B). This effect was additionally observed in MC38, despite the poor delineation of a distinct Tpex population by scRNAseq in the latter model (Fig. 3C/D). By contrast, no increase in CD8^+^Tcf1^+^ proximity to CD11c^+^ DC was seen in the LLC model which could be confounded by the paucity of this progenitor population in this model (Fig. 3E/F).

**Fig. 3.**
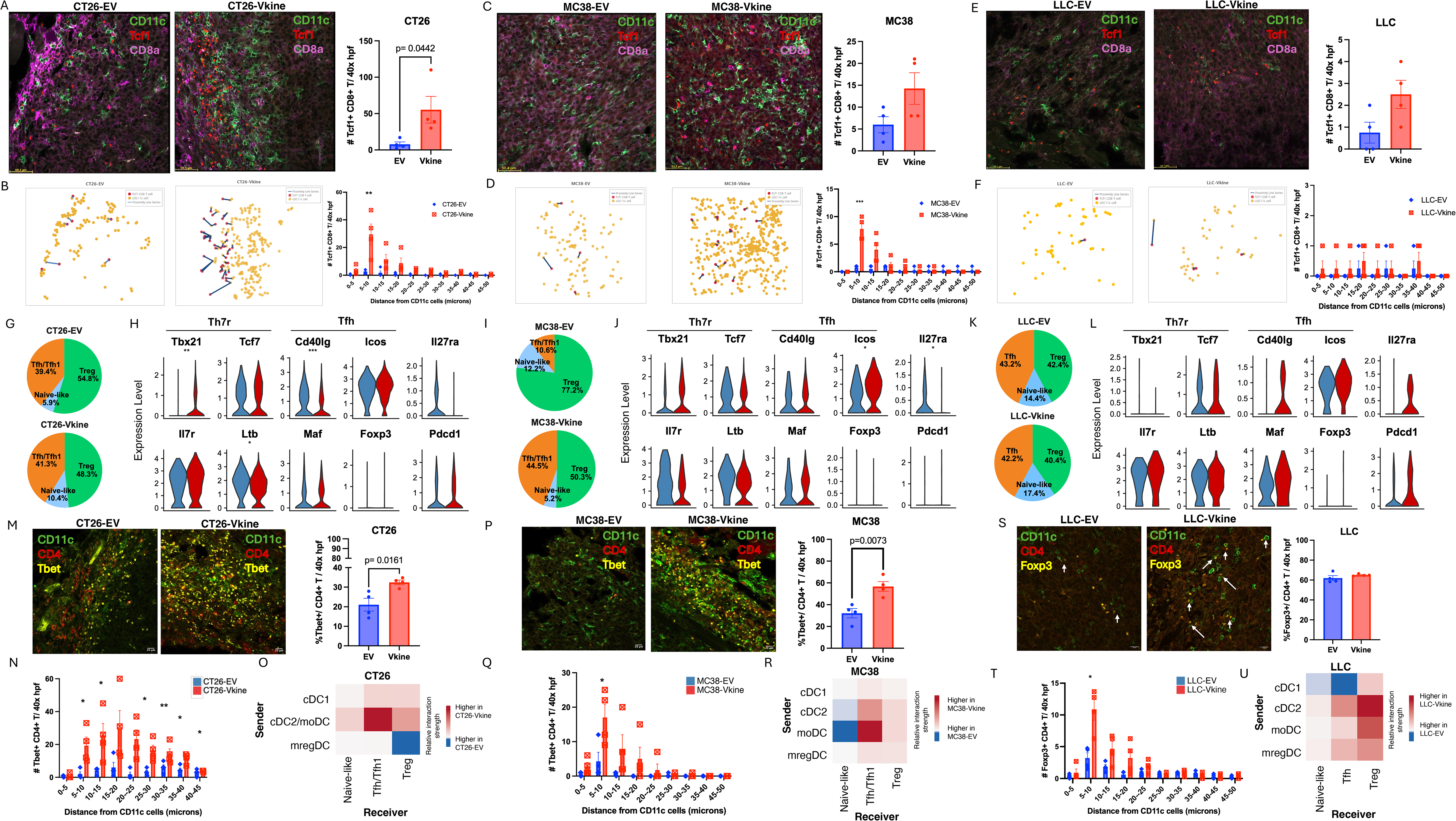
Th1-polarized Tfh (Th7R-like) and Tpex maintain physical proximity with DC in Vkine-replete TME. A: Confocal imaging analysis by immunofluorescence in CT26-EV and Vkine tumors using anti-CD11c, anti-CD8 and anti-Tcf1 antibodies reveals spatial distribution of DC and CD8^+^ Tcf1^+^ along the tumor rim. 40X objective: scalebar 50μm (A, left). The analysis confirms an increase in the density of CD8^+^Tcf1^+^ cells in CT26-Vkine vs CT26-EV conditions. B: HALO proximity analysis reveals an increase in the number of CD8^+^ Tcf1^+^ located within 30 μm to a CD11c^+^ DC peaking at the 5-10 μm range, suggesting physical interactions. Similar analysis of MC38-EV and -Vkine tumors (C and D) also reveals an increase in the rates of physical proximity between CD11c^+^ DC and CD8^+^Tcf1^+^ T cells despite an overall paucity of CD8^+^Tcf1^+^ cells. 40X objective: scalebar 50μm. E: By contrast, CD8^+^Tcf1^+^ are rare in both LLC-EV and LLC-Vkine. 40X objective: scalebar 50μm. F: Vkine does not increase the rate of proximity between CD11c^+^ and CD8^+^Tcf1^+^ T cells in LLC-Vkine compared to LLC-EV. G: Vkine promotes an expansion of Tfh cells in the CT26 TME, as determined by scRNAseq. H: Violin plots for gene expression of Tfh cells demonstrate expression of Th7R marker genes (*Tbx21, Tcf7, Il7r, Ltb*) with retention of key components of a core Tfh program (*Cd40lg, Icos, Maf*, negative for *Foxp3*). Il27r receptor downregulation is consistent with active signaling. I and J: In MC38-Vkine, the expansion of Tfh/Tfh1 cells is dramatic. K and L: In LLC, Tfh-like cells do not expand and lack key drivers of the Tpex-supportive Th7R program (lack of *Tbx21* expression). Because Tbx21 is essential for the Tpex-supportive functions of Th7R cells ^89^, these Tfh-like cells in LLC are likely dysfunctional. M: Immunofluorescence analysis confirms the transcriptomic data in CT26, demonstrating an increase in CD4^+^T-bet^+^ T cells in Vkine-replete TME that is reminiscent of the increase in T-bet-expressing CD4^+^ T cells in VPP clinical trial patients ^41^. 40X objective: scalebar 20μm. N: HALO proximity analysis demonstrates an increase in physical proximity between CD4^+^T-bet^+^ and CD11c^+^ DC in Vkine-replete TME. O: Heatmap depicting differential interaction strength between DC and CD4 subpopulations as inferred by CellChat. Red indicates increased signaling in Vkine-replete conditions and blue indicates decreased signaling. This demonstrates a shift in active signaling flow between CD11b^+^ DC and Tfh/Tfh1 in Vkine-replete TME compared to EV. P: Similar changes are seen in the MC38-Vkine context, compared to EV, including an increase in CD4^+^T-bet^+^ T cells in MC38-Vkine tumor rim (40X objective: scalebar 20μm), increase in physical interactions between CD11c^+^ DC and CD4^+^T-bet^+^ T cells (Q) and active signaling flow between CD11b^+^ DC and Tfh/Tfh1 (R). LLC demonstrates no increase in CD4^+^Foxp3^+^ Tregs in Vkine-replete conditions (40X objective: scalebar 20μm) (S); however, CD11c^+^ DC and CD4^+^Foxp3^+^ Tregs gain physical proximity in the presence of Vkine (T). U: Functional signaling flow occurs between CD11b^+^ DC and Treg as well as dysfunctional Tfh in LLC-Vkine. Data depict mean ± SEM. Statistical comparisons by unpaired Student’s t-test. Statistical significance is provided as the exact p value or denoted by an asterisk, *p < 0.05; **p < 0.01; ***p < 0.001, ****p < 0.0001. p values less than 0.05 were considered statistically significant.

### DC engage distinct CD4^+^ partners in Vkine-permissive and non-permissive microenvironments

In experimental models, Vkine promotes triad formation both in permissive (VCAN^lo^: CT26, MC38) and non-permissive (VCAN^hi^: LLC) contexts with starkly different functional outcomes (Fig. 1L), suggesting that triads in non-permissive contexts are functionally impaired or alternatively polarized. We hypothesized that the CD4^+^ partners may be Tpex-supportive in permissive contexts but non-Tpex-supportive in Vkine-nonpermissive contexts.

In Vkine-replete TME, CT26 and MC38 tumors demonstrated an expansion of CD4^+^ T cells bearing a Tfh phenotype *(Cd40lg, Icos, Maf)* that was modest in CT26 (Fig. 3G) but significant in the MC38 model (Fig. 3I/J). In LLC, the proportion of Tfh-like cells over total CD4^+^ remained relatively constant (Fig. 3K), however CD4^+^ cells were increased in absolute terms in Vkine-replete TME (Supp. Fig. S1I).

Importantly, we observed that in the immunogenic MC38 and CT26 models, Vkine promoted the transition towards a more activated, effector-like and Th1-polarized phenotype in Tfh cells (hence our designation as Tfh/Tfh1 in MC38-Vkine and CT26-Vkine). Transcriptomic analysis demonstrated an increase in T-bet (*Tbx21*) expression in MC38-Vkine and CT26-Vkine Tfh that was not seen in LLC-Vkine Tfh (Fig. 3H/J/L). Accordingly, IF staining confirmed the results of transcriptomic profiling demonstrating a relative increase in CD4^+^T-bet^+^ in CT26-Vkine and MC38-Vkine (Fig. 3M/P). Global transcriptomic analysis also demonstrated the acquisition of a Th1-polarized program in MC38 and CT26 Tfh/Tfh1 but not in LLC-Vkine Tfh-like cells (Supp. Fig. S6A/G/M). Indeed, GSEA analysis showed that Vkine promoted Tfh transcriptional programs consistent with effector and Th1-responses and corresponding reduction in gene sets supporting B cell homeostasis and humoral responses that are associated with a core Tfh identity: these changes were consistent in MC38- and CT26-Vkine models but not observed in LLC-Vkine tumors (Supp. Fig. S6B/H/N).

The fact that Vkine promoted Th1-like gene sets in Tfh cells was reminiscent of the clinical trial data, demonstrating an increase in T-bet^+^ CD4^+^ T cells in VPP patients ^41^. A T-bet expressing CD4^+^ lineage (termed Th7R) was recently reported to nurture Tpex in triads ^59^. Indeed, our expanded Tfh populations in Vkine conditions expressed many of the lineage markers associated with Th7R (*Il7r*, *Ltb*, *Tcf7*)(Fig. 3H/J/L). To determine whether our Tfh/Tfh1 transcriptional profile was similar to the reported Th7R program, we evaluated expression of a module score and confirmed that the transcriptional identity of Tfh/Tfh1 cells overlapped with the reported Th7R gene signature (Supp. Fig. S6C/I/O). This finding strengthens our hypothesis that Vkine results in expansion of a Tpex-supportive Th7R CD4^+^ subset in permissive microenvironments. By contrast, Tfh-like cells in VCAN^hi^ TME (LLC model) lack evidence of maturation towards a functional Th7R program (Supp. Fig S6D/J/P and Supp. Fig. S6E/K/Q).

Taken together, the above observations suggest that CD4^+^ Tfh cells recruited to Vkine-permissive TME engage in interactions with Vkine-activated cDC2 that impart or amplify a Th7R transcriptional profile. Recruitment of CD4^+^ Tfh cells to intratumoral triads may be facilitated by Vkine-induced CXCL9/10-CXCR3 signaling (Supp. Fig. S6F/M/S) which has been shown to be critical for TLS networking and anti-tumor immunity ^84^. We hypothesize that these “second-touch” interactions in the TME ^85^ expand Tfh/Tfh1 cells that can support Tpex in triads with cDC1. HALO proximity analysis confirmed that in CT26-Vkine and MC38-Vkine, DC physically interacted with T-bet-expressing CD4^+^ T cells (Fig. 3N/Q). CellChat analysis corroborated our hypothesis, demonstrating robust crosstalk between cDC2 and Tfh/Tfh1 in CT26-Vkine and MC38-Vkine (Fig. 3 O/R). In LLC, characterized by excess parental VCAN, DC physically interacted mostly with Treg (Fig. 3S/T) but not CD4+Tbet+ cells (Supp. Fig. S7A). CellChat analysis demonstrated signaling flow between cDC2 and Treg as well as dysfunctional (non-Th1 polarized) Tfh (Fig. 3U). Interestingly, CellChat analysis also demonstrated that in MC38 tumors (characterized by robust expansion of Tfh/Tfh1 cells), Vkine diverted ICOSL-ICOS signaling away from Treg and towards Tfh/Tfh1 (Supp. Fig. S6L and Supp. Table S3).

### Tumors replicating the VPW-VCAN^lo^ human phenotype are eradicated by Vkine plus single-agent checkpoint inhibition with resistance to re-challenge

Vkine’s role in assembly of Tpex-supporting immune triads would be expected to result in exquisite sensitivity to, and synergy with, ICI. To test this hypothesis, we implanted CT26- and MC38-Vkine tumors subcutaneously and allowed tumors to grow until Day 3, when EV controls were approximately 4mm in diameter. Anti-PD-L1 antibody was administered according to the schedule shown in Fig. 4A. Both CT26 and MC38 are moderately responsive to anti-PD-L1 immunotherapy at baseline, and indeed anti-PD-L1 immunotherapy alone resulted in 2/16 complete responders in the CT26-EV (Fig. 4B) and 3/10 complete responders in the MC38-EV condition (Fig. 4C). Complete responses were dramatically increased in the presence of Vkine (10/17 and 6/11, respectively) (Fig. 4B/C). MC38-Vkine complete responders re-challenged with MC38 tumor cells at Day 60 after primary inoculation rejected the challenge, suggesting immune memory to MC38 tumor antigens (Fig. 4D). Re-challenged mice were subjected to a tertiary challenge with a heterologous tumor expressing human Vkine (LLC). As expected, LLC tumors grew without impediment in MC38-Vkine memory mice demonstrating that cured mice had developed immune memory against MC38-specific tumor antigens but not heterologous LLC antigens or putative human Vkine epitopes (Fig. 4E).

**Fig. 4.**
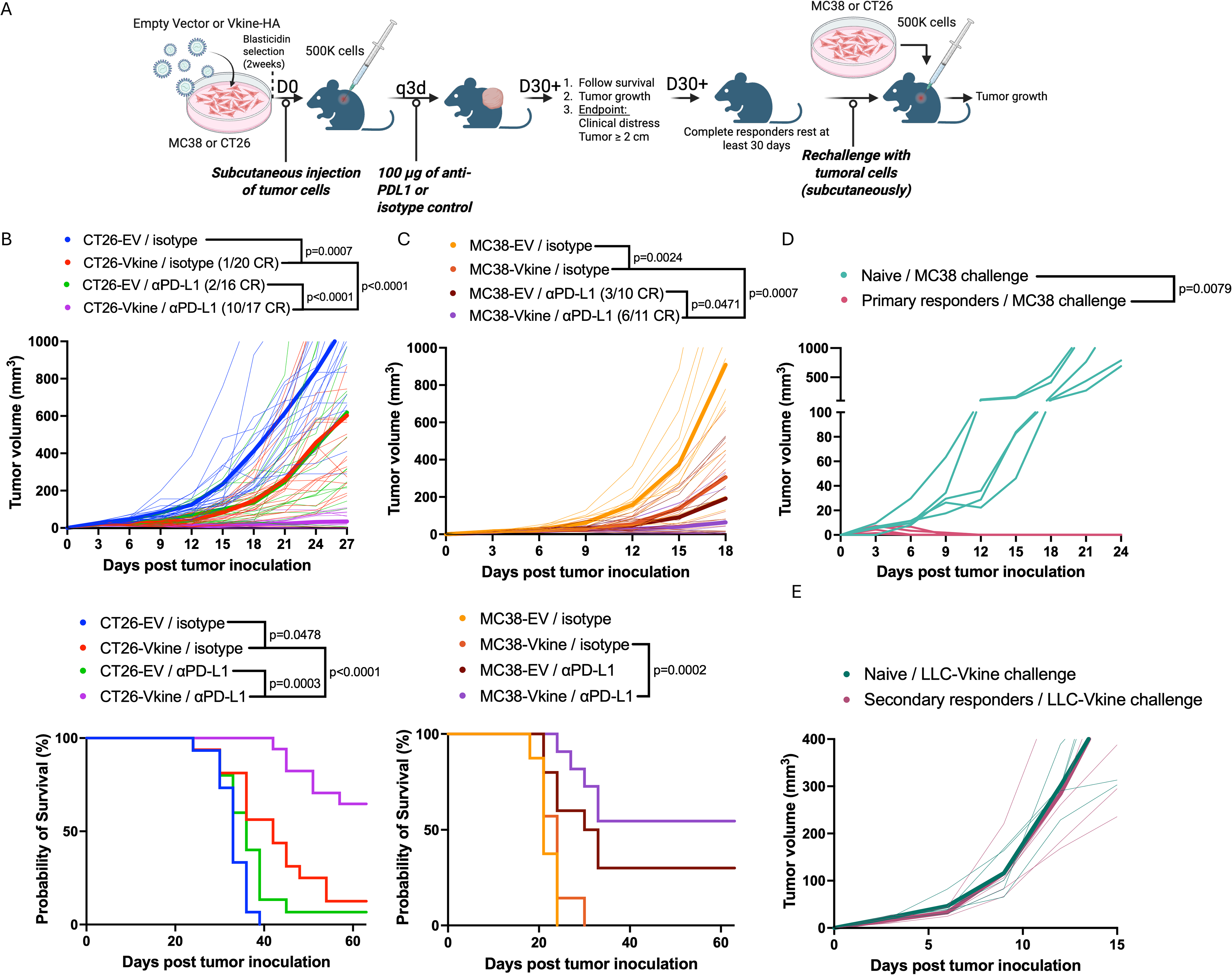
Vkine synergy with PD-1 pathway checkpoint inhibition cures a subset of tumors with immune memory to rechallenge. A: Schematic layout of *in vivo* experiments demonstrating workflows of tumor implantation, treatment and tumor harvesting. B: Anti-PD-L1 treatment of CT26-Vkine demonstrates robust responses with 10/17 complete responders and prolongation of overall survival. Tumor growth curves shown as spider plots (thin lines) and means (thick lines). Corresponding Kaplan-Meir survival curves are shown to the bottom (n=16-20 per arm). Interestingly, 1/20 CT26-Vkine tumors regressed spontaneously without further therapy. Similar results were seen in MC38 tumors where Vkine synergy with anti-PD-L1 therapy resulted in complete responses (6/11) demonstrated as growth curve spider plots (C, top) and Kaplan Meir survival curves (n=10-11 per arm)(C, bottom). D: Rechallenge of cured MC38-Vkine mice with native MC38 cells (not engineered to express EV or Vkine) demonstrated a complete resistance to re-challenge, indicating long-term immune memory to MC38 tumor antigens (n=5). E: Mice that survived the secondary rechallenge received a tertiary challenge with LLC-Vkine tumor cells (n=5). These mice were unable to reject LLC-Vkine tumors demonstrating that their immune memory was specific to MC38 tumor antigens, not those of a heterologous tumor (LLC) or putative antigens encoded by human Vkine sequences. Tumor growth curves were analyzed by Mann-Whitney test and survival curves by logrank (Mantel-Cox) test. Statistical significance is provided as the exact p value or denoted by an asterisk, *p < 0.05; **p < 0.01; ***p < 0.001, ****p < 0.0001. p values less than 0.05 were considered statistically significant.

### Tumors replicating the poor-prognosis VPW-VCAN^hi^ human phenotype respond to combination immunotherapy targeting Treg or immunoregulatory DC

We engineered LLC-based models in order to replicate the poor-prognosis VPW-VCAN^hi^ subset. LLC tumors are known to be resistant to PD-1 pathway ICI and Vkine synergy with single-agent PD-1 inhibition is modest in this model ^39^. Based on our mechanistic understanding of Vkine’s activities in non-permissive microenvironments, we sought to devise rational combinatorial immunotherapy regimens that would synergize with Vkine in the extremely unfavorable TME replete with high levels of residual unprocessed VCAN (Fig. 5A).

**Fig. 5.**
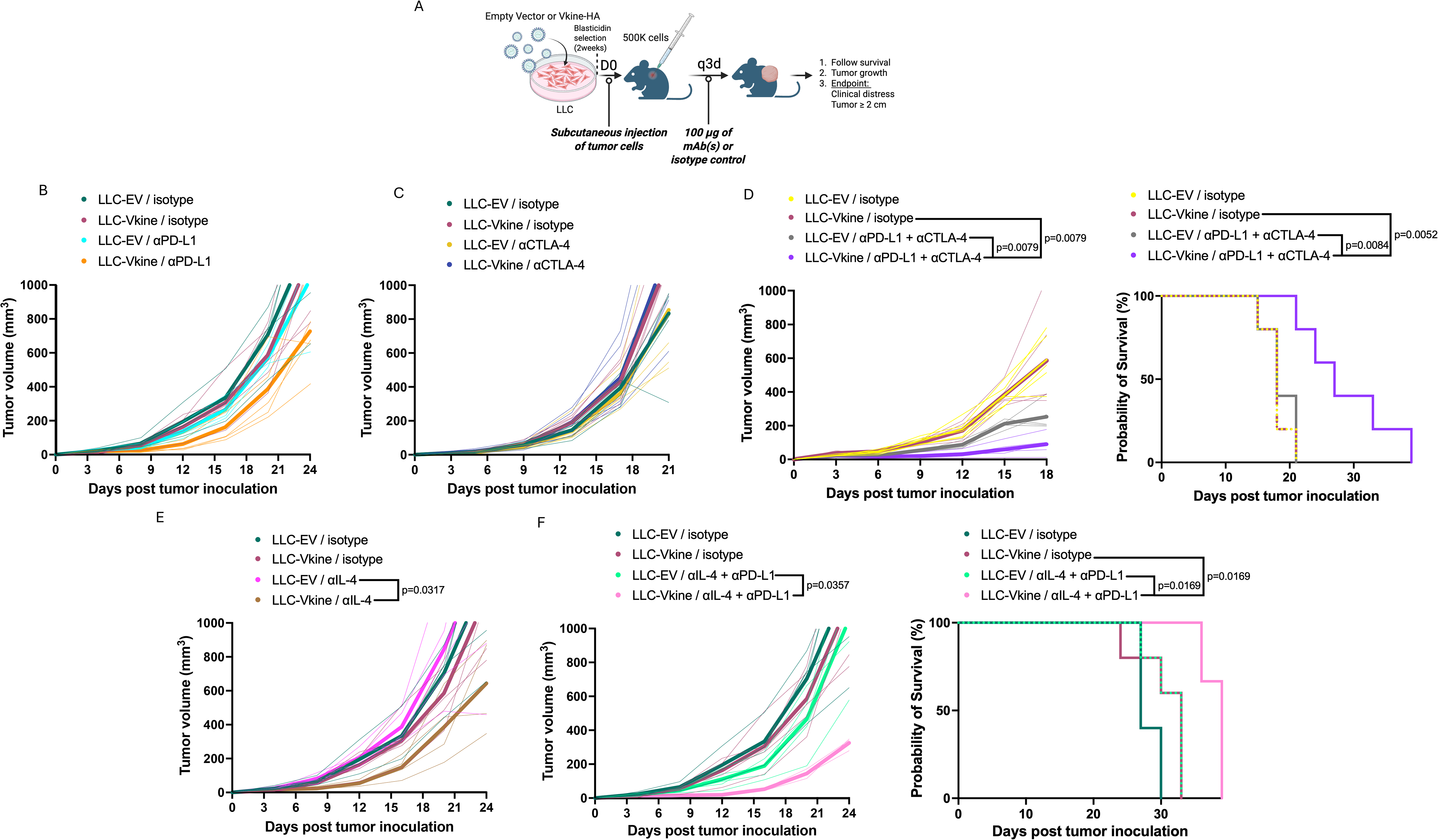
The adverse effect of unproteolyzed parental VCAN can be overcome through rational immunotherapy combinations targeting Treg or immunoregulatory DC. A: Schematic of *in vivo* experiments demonstrating workflows of tumor implantation, treatment and tumor harvesting. B: Single-agent anti-PD-L1 monotherapy in the LLC model (modeling VPW-VCAN^hi^ human cancers) has modest activity in both control (EV) and -Vkine replete conditions (n=3-5 per arm). C: Single-agent anti-CTLA4 monotherapy in the LLC model has modest activity even in Vkine-expressing tumors (n=5 per arm). D: Combination anti-PD-L1/anti-CTLA4 immunotherapy reveals synergy with Vkine, resulting in prolongation of overall survival (n=5 per arm). E: Single-agent anti-IL4 monotherapy has modest synergy with Vkine in LLC tumors (n=4-5 per arm). F: Combination anti-PD-L1/anti-IL4 immunotherapy targeting mregDC ^87^ reveals robust synergy with Vkine, resulting in prolongation of overall survival (n=3-5 per arm). Tumor growth curves were analyzed by Mann-Whitney test and survival curves by logrank (Mantel-Cox) test. Statistical significance is provided as the exact p value or denoted by an asterisk, *p < 0.05; **p < 0.01; ***p < 0.001, ****p < 0.0001. p values less than 0.05 were considered statistically significant.

We demonstrated earlier that in the presence of excess unprocessed VCAN, cDC2 engage Treg preferentially. We therefore hypothesized that Treg-targeting immunotherapy may rationally synergize with Vkine and anti-PD-L1 (Fig. 5B). Among several experimental approaches to deplete Treg, one of the most efficient in mouse models employs anti-CTLA4 antibodies ^86^. Whereas anti-CTLA4 antibodies were ineffective as monotherapy in Vkine-expressing LLC (Fig. 5C), the combination of anti-CTLA4 and anti-PD-L1 resulted in clinically meaningful responses with improved animal survival (Fig. 5D).

A second approach was based on our observation that Vkine promoted repolarization of mregDC towards an enhanced immunogenic profile (Fig. 2J/L/N). Interestingly, mregDC repolarization was evident in LLC-Vkine, i.e., the effect was unaffected by the presence of excess unproteolyzed VCAN (Fig. 2N). We demonstrated a smaller, albeit reproducible, benefit of single-agent anti-IL4 in Vkine-expressing LLC tumors (Fig. 5E). Combination immunotherapy with anti-IL4 and anti-PD-L1 previously shown to reprogram mregDC ^87^ results in meaningful synergy with Vkine, again resulting in significant benefit in the experimental animal cohort (Fig. 5F).

The choice of immunotherapy combinations in Vkine-nonpermissive TME was based on mechanistic understanding of Vkine’s activities. Indeed, Vkine did not synergize with anti-LAG3 ^+^ anti-PD-L1 double immunotherapy in the LLC model (Supp. Fig. S7B/C). The results demonstrate that rational, mechanism-based, Vkine-incorporating immunotherapy combinations can at least partially overcome the inhibitory effect of excess unproteolyzed VCAN in the TME (VPW-VCAN^hi^ phenotype).

### Vkine delivered as LNP-mRNA or recombinant protein has monotherapy activity dependent on CD8^+^ T cells and cDC1 and synergizes with checkpoint inhibition

Immunocompetent syngeneic models expressing ectopic Vkine have provided useful tools to study the steady-state effects of Vkine in the TME. To determine the efficacy and relevance of Vkine as immunotherapy, we sought to determine whether Vkine delivered as mRNA in lipid nanoparticles or recombinant protein would produce similar effects, as predicted from the ectopic Vkine-expressing models.

We incorporated human Vkine sequences in an *in vitro* transcription vector and generated Vkine mRNA (Supp. Fig. S8A) that was subsequently packaged in lipid nanoparticles. Nanoparticles were typically 60-70nm in diameter and were optimized for *in vivo* delivery (Supp. Fig. S8B). Control mRNA delivered in LNP resulted in efficient translation in the TME (Supp. Fig. S8C). MC38 tumors were implanted and allowed to grow to a size of approximately 4 mm diameter (Fig. 6A). Starting on Day 7, LNP-Vkine mRNA or control LNP-Thy1.1 mRNA (Supp. Fig. S8C) were delivered intratumorally on days 7, 9 and 11. In some animals, anti-PD-L1 antibodies were administered alone or in combination with LNP-Vkine mRNA. As shown in Fig. 6B, LNP-Vkine mRNA demonstrated monotherapy activity with 1/10 mice becoming complete responders with LNP-Vkine mRNA alone. As discussed earlier, MC38 demonstrated partial sensitivity to anti-PD-L1 monotherapy with 2/10 complete responders in that arm. The combination produced striking synergy with 5/10 complete responders after a limited course of 3 LNP doses. LNP-Vkine mRNA plus anti-PD-L1 synergy resulted in an overall survival advantage (Fig. 6B). Adoptive transfer of pan-T cells from complete responders conferred a significant degree of immunological memory to secondary tumor recipients (Fig. 6C). Lastly, we demonstrated that Vkine mRNA response required cDC1 and CD8^+^ T cells, as the response was abrogated in the absence of either cDC1 or CD8^+^ T cells (Fig. 6D/E).

**Fig. 6.**
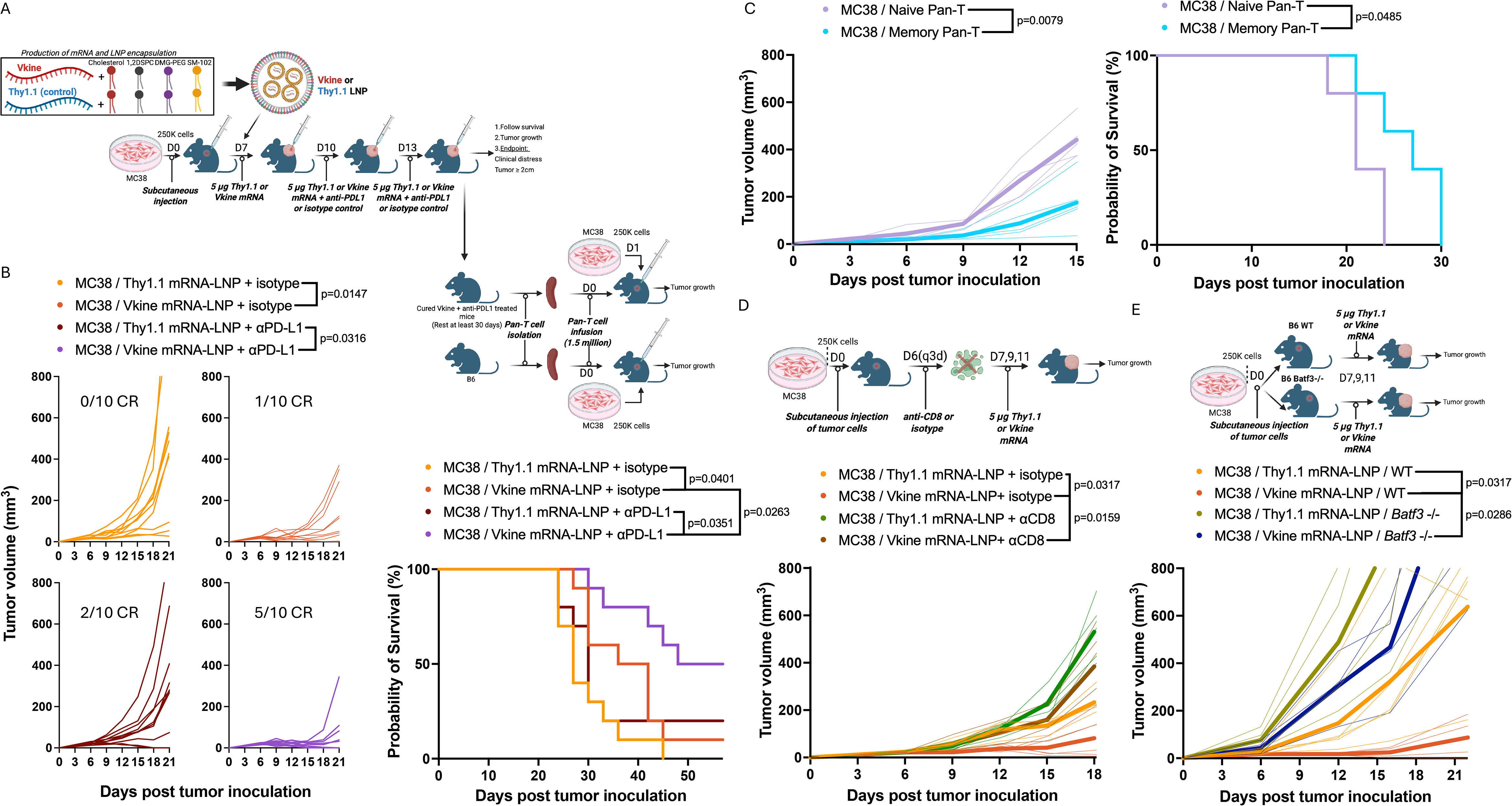
Vkine mRNA delivered in lipid nanoparticles has monotherapy activity and displays synergy with anti-PD-L1 checkpoint inhibition that is dependent on cDC1 and CD8^+^ T cells. A: Schematic layout of *in vivo* experiments demonstrating workflows of tumor implantation, treatment and tumor harvesting. B: Spider plots of individual tumor growth curves following intratumoral injection of LNP-Vkine mRNA vs Thy1.1 mRNA control, with and without concurrent anti-PD-L1 immunotherapy (B, left). Corresponding survival curves are shown in B, right (n=10 per arm). C: Pan-T cells from mice cured after LNP-Vkine mRNA administration were adoptively transferred into naïve syngeneic recipients. Following adoptive transfer, mice were challenged with MC38 tumor and demonstrated partial resistance to the rechallenge (n=5 per arm)(growth curves to the left and Kaplan Meir survival curves to the right). D: CD8 depletion abrogated the responses to LNP-Vkine mRNA. Spider plots (individual tumors, thin lines) and mean growth curves (thick lines), shown for each experimental arm (n=5 per arm). E: *Batf3*-null mice (deficient in cDC1^79^) were resistant to LNP-Vkine mRNA. Spider plots of individual tumors (thin lines) and mean growth curves (thick lines) shown for each experimental arm (n=4-5 per arm). Tumor growth curves were analyzed by Mann-Whitney test and survival curves by logrank (Mantel-Cox) test. Statistical significance is provided as the exact p value or denoted by an asterisk, *p < 0.05; **p < 0.01; ***p < 0.001, ****p < 0.0001. p values less than 0.05 were considered statistically significant.

The encouraging results with LNP-Vkine mRNA prompted us to test the effect of recombinant Vkine (rVkine) protein injected intratumorally (Fig. 7A). We followed the process for propagating Vkine-producer cells and rVkine purification from the supernatant delineated in ^88^. Briefly, rVkine purification was accomplished directly from HEK-293T cell culture supernatant (Fig. 7B) through Ni^2+^ immobilized metal affinity chromatography (IMAC), as delineated in Materials and Methods (Supp. Fig. S9A). The elution was collected in fractions (Supp. Fig. S9B). Following collection of the eluate and flow through, initial validation of rVkine recovery was performed using SDS-PAGE and Coomassie blue staining (Supp. Fig. S9C). Fractions exhibiting a predominant band at 75kDa, typically around the 200mM imidazole range, were pooled for further recovery (Fig. 7C and Supp. Fig. S9C). rVkine was buffer exchanged into PBS and tested for endotoxin contamination. Final validation was performed through western blotting of rVkine with several antibodies targeting distinct Vkine domains (Fig. 7D) as well as Ponceau staining of the membranes to identify any degradation products.

**Fig. 7.**
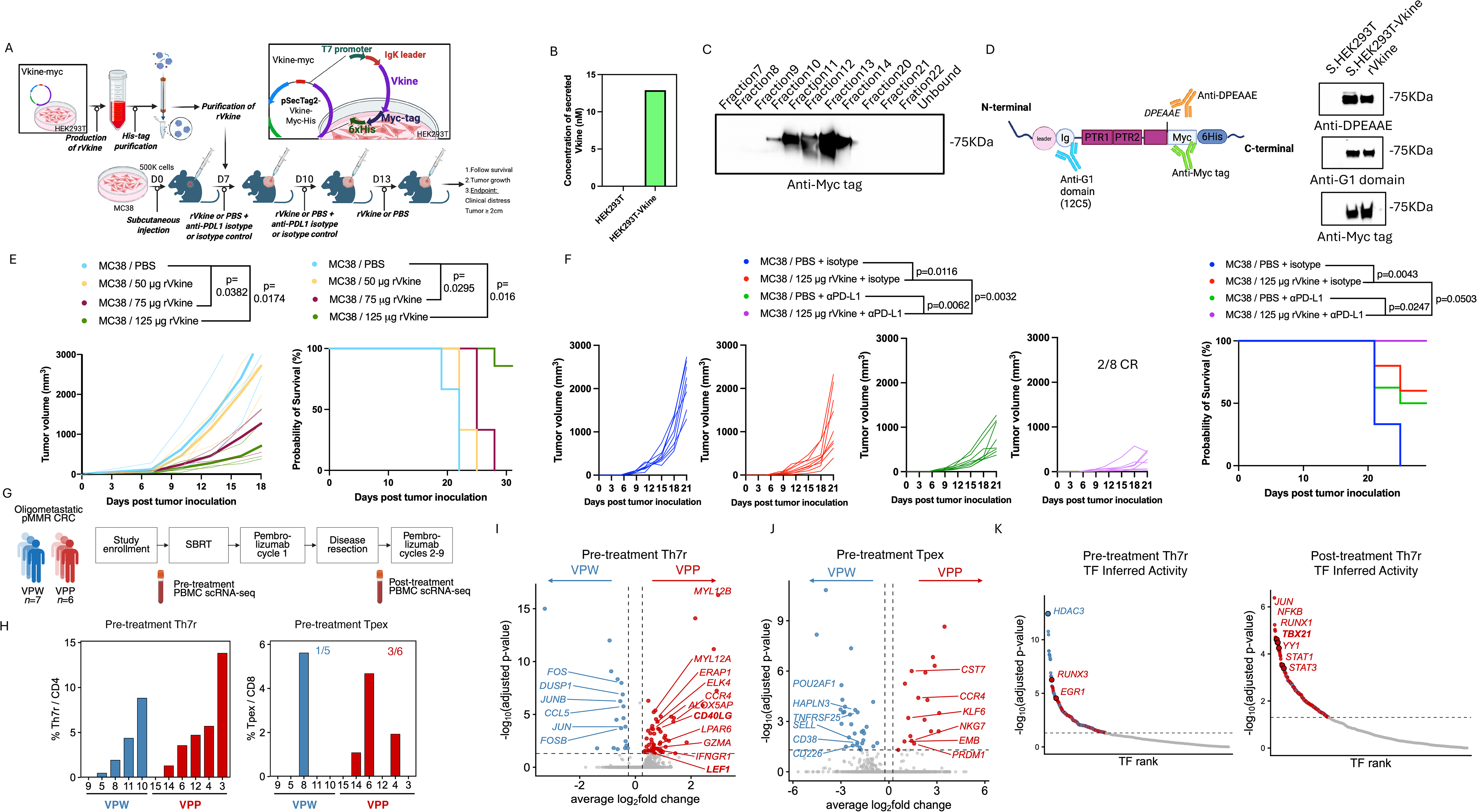
Recombinant Vkine (rVkine) reverses the immunodeficiency associated with the VPW phenotype and synergizes with PD-1 pathway checkpoint inhibition. A: Schematic of *in vivo* experiments demonstrating workflows of tumor implantation, treatment and tumor harvesting. B: rVkine secreted into the culture supernatant of producer 293T cultures. C: Chromatography fractions demonstrating elution of rVkine migrating at app. 75kD (anti-myc tag immunoblot). D: Schematic demonstrating the Vkine domain structure with targets for each antibody depicted. Immunoblots showing Vkine migrating at 75kD using each detection antibody. E: *In vivo* dose-response curves for IT administration of escalating doses of rVkine. To the left are shown growth curves and to the right are shown Kaplan-Meir survival curves (n=3 per arm). F: Spider plot growth curves (F, left) and survival plots (F, right) demonstrating synergy between rVkine and anti-PD-L1 checkpoint inhibition immunotherapy (n=8 per arm). G: Clinical trial schema and sample collection timepoints ^41^. H: scRNAseq data for each of the timepoints obtained from GEO (GSE316301) and were analyzed as in Methods. Percentages of Th7R CD4^+^ and Tpex CD8^+^ are shown for each patient with pre-treatment data, stratified per VPP and VPW cohorts. I and J: Volcano plots demonstrating differential gene expression in VPP vs VPW pre-treatment Th7R and Tpex cell populations. Red, higher in VPP; blue, higher in VPW. K: Transcription-factor inferred activity is shown in pre-and post-treatment Th7R populations. Red, higher in VPP; blue, higher in VPW. Tumor growth curves were analyzed by Mann-Whitney test and survival curves by logrank (Mantel-Cox) test. Statistical significance is provided as the exact p value or denoted by an asterisk, *p < 0.05; **p < 0.01; ***p < 0.001, ****p < 0.0001. p values less than 0.05 were considered statistically significant.

rVkine demonstrated anti-tumor effects when injected intratumorally, in a dose-dependent manner (Fig. 7E). In a limited course-experiment (3 intratumoral doses), rVkine synergized with anti-PD-L1 and the synergy produced 2/8 complete responders (Fig. 7F). rVkine synergy with anti-PD-L1 produced a significant survival benefit in the combined treatment cohort (Fig. 7F).

### Circulating Th7R in VPP human patients display enhanced Tpex-supportive profiles

To further confirm the human clinical relevance of our findings, we analyzed the results of the first clinical trial employing VCAN proteolysis as a prospective biomarker to predict outcomes to PD-1 ICI (pembrolizumab) in oligometastatic colorectal cancer ^41^. Briefly, the clinical trial scheme is summarized in Fig. 7G. Patients with oligometastatic colorectal cancer underwent SBRT to hepatic deposits followed by one round of pembrolizumab. The disease was then resected and patients received additional adjuvant pembrolizumab to progression. As previously shown ^41^, VPP patients had the most favorable outcomes with 100% patients alive at last follow-up and median RFS of 3.78 years. Patients with VPW phenotype fared much more poorly (median RFS at 0.83 years). Peripheral blood samples were collected at enrollment and following disease resection and CD45^+^ fractions were subjected to scRNAseq (data available under GSE316301). Following a pre-processing, filtering, and quality control pipeline adapted from the original study, broad immune lineages were identified using *de novo* markers and canonical gene expression (Supp. Fig. S9D/E). To identify Th7R cells, the CD4^+^ compartment was re-clustered, and a 58-gene discriminatory signature derived from sorted Th7R, Th1, and Th17 cells ^59,89^ was applied to SELL^low^ CD4^+^ clusters (Supp. Fig. S9F/G). Clusters 12 and 16_0 were annotated as Th7R based on signature enrichment and expression of Th7R-specific markers such as *IL7R*, *TCF7*, *NELL2*, *LTB*, *GZMK*, and *CCL5* (Supp. Fig. S9H). To identify Tpex, the CD8^+^ compartment was re-clustered and examined for expression of signature genes of Tpex. Two clusters demonstrated a *PDCD1*^high^ (PD-1^high^) *IL7R*^+^ *TCF7*^+^ *SLAMF6^+^ GZMK*^+^ *GZMB*-profile consistent with a Tpex identity; however, one cluster was excluded due to absence of *CXCR5* and *CCR7* expression, which are required to support the annotation as Tpex (Supp. Fig. S9I).

Despite the small number of patients in each cohort (VPP/VPW), we detected a trend towards an overrepresentation of Th7R as a percentage of the total circulating CD4 compartment in VPP vs VPW. There was a stronger trend in the representation of Tpex cells among circulating CD8^+^ T cells, with 3/6 patients in the VPP cohort demonstrating sizeable Tpex compartments vs 1/5 in the VPW cohort (Fig. 7H). Gene expression analysis demonstrated unique Th7R profiles among VPP patients compared to VPW patients. In the former, Th7R expressed significantly higher levels of *CD40LG*, stemness markers (*LEF1*) and effector markers (*GZMA*), suggesting an enhanced functional capacity to provide Tpex support within immune triads (Fig. 7I). Interestingly, both Th7R and Tpex in VPP patients expressed non-canonical *CCR4*, suggesting enhanced ability to compete with Treg for DC-attraction and retention into triads (Fig. 7I/J). Transcriptional factor activity inference analysis revealed enhanced *TBX21* pathway activity post-treatment in VPP patients, again underscoring enhanced Tpex-supportive roles in VPP Th7R (Fig. 7K). Taken together, the data suggest that VPP conditions may promote the tumoral infiltration, retention and organization of Th7R and Tpex into functional intratumoral immune triads.

## DISCUSSION

Apposed DC:CD4^+^T:CD8^+^T units are now established as a requirement for anti-tumor immunity: intratumoral triads are necessary for immunotherapy-mediated tumor elimination ^50^, three-cell clusters are a conserved feature of endogenous anti-tumor responses ^90^ and cDC1 presenting on both MHC classes are required for TLS development and maintenance ^6^. What makes these units assemble *in situ* is unknown. Here we identify a matrix fragment that does: proteolytic release of the versican matrikine versikine (Vkine) assembles the DC:CD4⁺T:CD8⁺T triads that constitute the founding units of TLS, and supplying Vkine therapeutically confers sensitivity to checkpoint blockade in tumors that lack endogenous pathway activity. We discuss below how this places provisional matrix remodeling upstream of tumor lymphoid neogenesis, and why it argues for harnessing stromal signals rather than removing them.

The peritumoral stroma constitutes the first theater of encounter between the developing tumor and the host, and homeostatic responses mounted there (immunoediting: elimination/equilibrium) require a milieu permissive to immune infiltration, such as the provisional matrix of wound healing ^48,91^. Within provisional matrix, the regulated proteolysis of the large matrix proteoglycan versican (VCAN) plays a cardinal role in regulating cell-fate decisions in both development and adulthood. Adult mice bearing targeted disruption of the Vkine-generating cleavage site show accelerated wound healing, with faster progression to fibrosis and wound closure ^46^, as well as resolution of inflammatory colitis ^92^. These results strongly suggest that VCAN proteolysis prolongs inflammation whereas its attenuation is associated with resolution of inflammation and healing ^29,92^. Consistent with this, VCAN proteolysis associates with T-cell inflammation across solid and hematologic malignancies.

VCAN proteolysis acts through two effects that are related by a protease but functionally separable. Intact VCAN has a significant adverse effect on T-cell trafficking and activity in both cancer and non-malignant inflammation ^93^. Tumors that are rich in intact VCAN demonstrate robust immune exclusion, an effect likely dependent on signaling from both the protein core and the chondroitin-sulfate GAG chains ^37^. Independently, Vkine engages adaptive immunity by regulating the abundance and function of tumor-antigen cross-presenting, Batf3-dependent, conventional dendritic cells (cDC1) ^32,43,64^. That these constitute two activities rather than two ends of a single lever is indicated by patient stratification: VPP tumors outperform VPW-VCAN tumors ^41^. Were Vkine’s contribution merely the relief of intact VCAN, tumors with little VCAN would have nothing to relieve and should fare at least as well as those with active proteolysis. Vkine therefore supplies something that the absence of VCAN does not.

In this manuscript we demonstrate that Vkine assembles the immune “triads” that constitute the units of TLS ^6^ and the functional orchestrators of immunotherapy responses ^50,73^. The data argue for a “niche-first” model in which a permissive milieu is pre-requisite for functional organization of chemokine-recruited cells; in this regard, tumors may borrow a playbook from normal lymphoid organ architecture, as VCAN proteolysis is a regular component of light zone structure. Vkine’s roles in promoting immune triads appear to be exerted mainly through its actions on DC: on the one hand, Vkine promotes cDC1 survival through an NK-dependent mechanism ^39^. On the other hand, Vkine strongly activates CD11b+ DC (cDC2 and moDC) to acquire an ISG+ activation state, previously associated with T-cell priming ^94^. The physical interaction and crosstalk between Vkine-activated cDC2 and Tfh ^67,95,96^ appears to promote, in the latter, the acquisition of a T-bet-driven transcriptional program, resembling the recently reported, Tpex-sustaining, Th7R module ^59,89,97^. In the presence of excess unprocessed VCAN, cDC2 swap their preferred partner to Treg or dysfunctional Tfh that fail to induce T-bet and thus fail to support Tpex ^98^ and may induce premature exhaustion ^99,100^.

Which receptor reads the matrix is a related question. VCAN has been previously reported to induce dysfunctional DC through TLR2 engagement ^101,102^, an activity expected to fall mainly on CD11b+ DC, as cDC1 do not express TLR2 ^87^; soluble biglycan, released as a damage-associated molecular pattern in autoimmunity, acts through the same TLR2/4 axis ^103^. Vkine, by contrast, acts independently of TLR2 ^39^. Proteolysis may therefore do more than attenuate a signal: by converting the parent proteoglycan into a fragment that its receptor no longer reads, it switches the matrix from one sensing pathway to another. The parent’s reader is defined; the fragment’s is not yet.

Our model ascribes CD4+-instructive functions to CD11b+ DC, such that Vkine-conditioned CD11b+ DC help determine the polarity and functionality of the resultant triad, with ICOS– ICOSL signaling flow diverted towards the Tfh1-polarized CD4+ partner and away from Treg. Chaib and colleagues independently report apposed multicellular units — tetrads, incorporating a macrophage — as a correlate of checkpoint response, and likewise find ICOS signaling to be required ^14^. They report that tetrad density did not track TLS density in their cohort and conclude that tetrads are microanatomically distinct from TLS. The convergence of independent approaches on apposed DC–T units as the functional correlate of checkpoint response is, in our view, the more notable point.

The proximal mechanism by which Vkine acts on dendritic cells remains unresolved. Activity is independent of TLR2, through which unproteolyzed VCAN signals, and of CD44, the principal hyaluronan receptor ^39^. A conventional receptor remains plausible but unidentified. Alternatively, Vkine (a large link-module-bearing fragment deposited pericellularly, unlike the compact basement-membrane matrikines) may act on the material properties of the glycocalyx rather than through a dedicated receptor ^104^. Such a mechanism has precedent in immune cells. The kinetic segregation model holds that steric exclusion of bulky surface phosphatases such as CD45 and CD43 from close-contact zones is what permits T-cell receptor triggering ^105^, and sialic acid density on myeloid cells sets the threshold for Siglec engagement ^106^: in both cases the dimensions and charge of the pericellular layer, rather than a ligand, govern whether signaling proceeds. A Vkine-decorated pericellular coat may thus constitute a distinct material state — not equivalent to the simple absence of matrix, as the stratification data above indicate — that lowers the threshold for innate sensing, consistent with the heightened DNA-sensing responsiveness we observe ^39^.

Two limitations bear emphasis. First, establishing that VCAN proteolysis is necessary for triad assembly requires cleavage-resistant-Vcan animals paired with a tumor model displaying robust endogenous Vcan-proteolysis at the site that generates Vkine; such a Vcan-proteolytic tumor model has not yet been identified. Asano et al demonstrated detectable VCAN-V1^441–442^ proteolysis by Western blot from xenograft tumor lysates derived from MDA-MB231 human breast cancer cells in addition to endothelial DPEAAE signal ^107^. They also looked at LLC and B16 melanoma syngeneic models: although they reported endothelial DPEAAE staining by IF, they did not demonstrate detectable DPEAAE signal by Western blot (as in MDA-MB231 xenografts) and, in our hands, none was found in LLC or the other models used here (Fig 1J). Rodriguez-Baena et al. reported modest anti-DPEAAE-reactive fragments by Western blot in some B16F1 tumors but noted high inter-sample heterogeneity and did not quantify the signal ^108^. It is intriguing to consider the dearth of VCAN-proteolytic mouse tumor models (at least regarding VCAN-V1^441–442^ in our screens) in the light of the fact that tumor models displaying spontaneous TLS are also scarce; whether the two phenomena are interconnected is unclear. Our conclusions therefore rest on gain-of-function, on the inverse phenotype produced by excess intact VCAN, and on human correlation, in which tumors lacking VCAN proteolysis also lack triads. This last association is not a substitute for genetic evidence, but it is the nearest equivalent available in the species of interest. Second, our germinal center data establish VCAN proteolysis as a constitutive feature of light zone architecture, restricted to the CD23⁺ FDC network and spatially coincident with the CXCL13 gradient that positions the light zone; whether proteolysis is required to establish or maintain that gradient remains to be determined.

Our data highlight prognostic and therapeutic implications of a matrix-signaling network spanning lymphoid tissue organization, wound-healing and anti-tumor immunity. Stratification according to VCAN proteolysis status can help predict benefit from checkpoint inhibition immunotherapy. VPP patients (40% of the clinical trial cohort ^41^) are best positioned as they display endogenous Vkine production with low residual VCAN. VPW patients (60% of the cohort ^41^) may benefit from precision Vkine-based immunotherapy to reverse the immunosuppression associated with low endogenous proteolysis activity. Vkine therapy, guided by the companion biomarker of endogenous VCAN proteolysis, epitomizes a new class of agents aimed at TLS induction ^11^. Moreover, our data highlight the need to reevaluate prior approaches to globally disrupt tumor stroma, an approach that has failed in late-phase trials and in some settings worsened outcomes ^109–112^, in favor of biomarker-driven, pathway-focused harnessing of stromal signals that modulate anti-tumor immunity.

## ACKNOWLEDGMENTS

This work was supported through the NIH/ National Cancer Institute (R01CA252937 to FA) and the Robert E. and Emily H. King Endowment at Rush University. We thank Cheryl Kim (LJI Flow Cytometry), Eric O’Connor (UCSD Moores Cancer Center), Leichu Liang (UCSD Moores Cancer Center), Victoria Quintana-Ribbens (UCSD Moores Cancer Center), Alejandro Rizo (UCSD Moores Cancer Center), Ricardo De-luna (UCSD Materials Research Science and Engineering Center), Art Nasamran, Daisy Chilin-Fuentes and Brin Rosenthal (UCSD CCBB), Jingting Yu (Salk Institute Genomics and Bioinformatics), Stefan Green and Ping Li (Rush Genomics), Ryan Deaton and Lucas Zecker (UIC Pathology), Balaji Ganesh (UIC Flow Cytometry), Katherine Badior and Christian Kastrup (Versiti Blood Research Institute, Milwaukee WI) for their expertise, advice and support. We thank the University of Wisconsin Translational Research Initiatives in Pathology (TRIP) Laboratory supported by the UW Department of Pathology and Laboratory Medicine and the UWCCC (P30 CA014520) for its multiplex immunohistochemistry services. We thank Suneel S. Apte (Cleveland Clinic) for the kind gift of Vkine-expressing HEK293T cells.

## AUTHOR CONTRIBUTIONS

DJL, DH, AP, GSY, EG, NL, YH, YD, MF, AG, EM, KP, PTT, SN and FA designed and/or performed experiments. KAM provided pathology expertise. GSY and SN performed bioinformatics analysis. AC, JM, JS, and DD provided crucial expertise and/or reagents. FA was overall responsible for design and conduct of the study and for securing funding. All authors reviewed, edited and/or revised the manuscript.

## DECLARATION REGARDING AI

An AI assistant (Claude, Anthropic) was used for language editing and manuscript preparation. All scientific content, data interpretation and conclusions are those of the authors, who take full responsibility for the work.

---

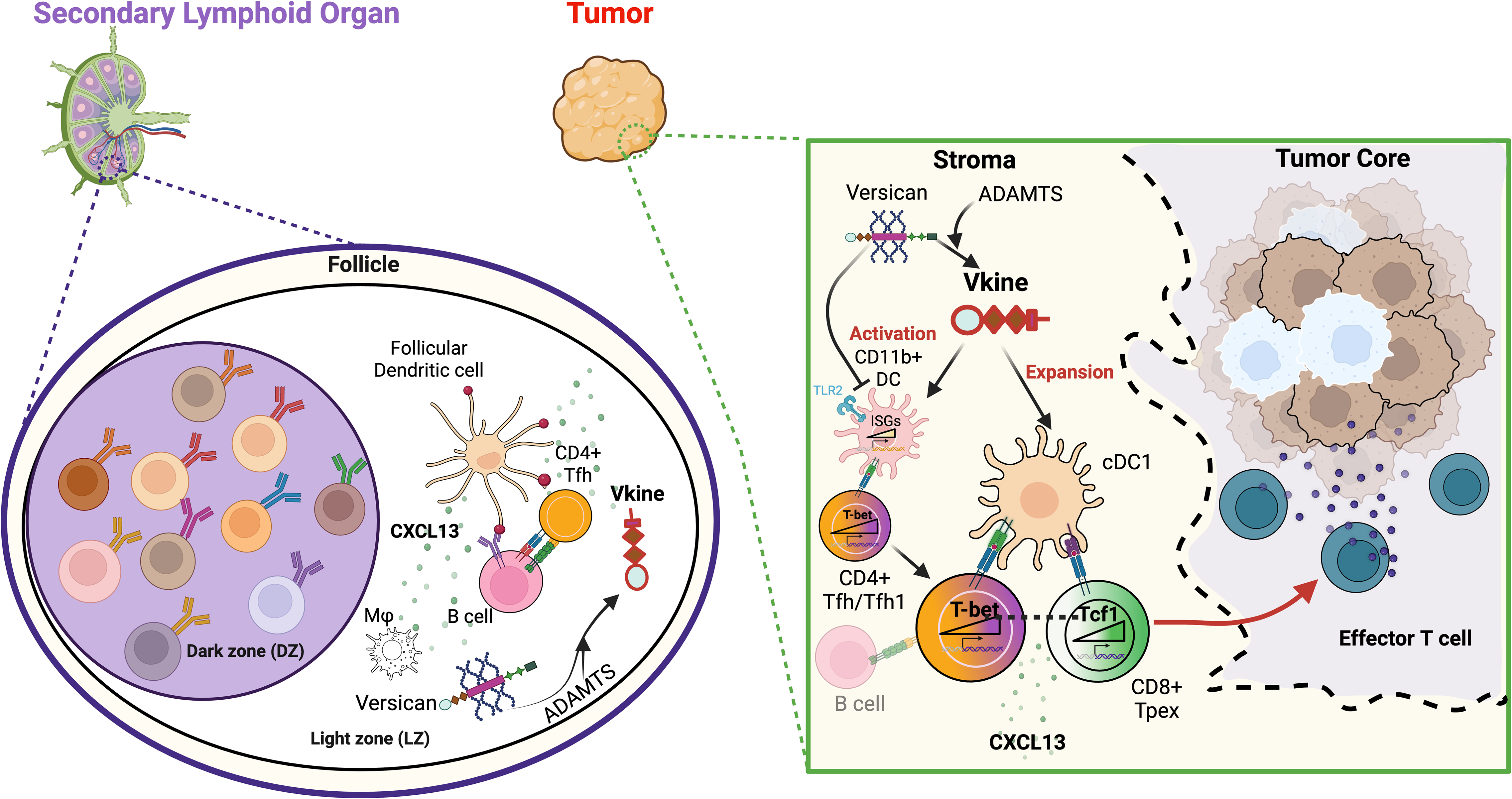

## REFERENCES

1. Teillaud, J.L., Houel, A., Panouillot, M., Riffard, C., and Dieu-Nosjean, M.C. (2024). Tertiary lymphoid structures in anticancer immunity. Nat Rev Cancer 24, 629–646. 10.1038/s41568-024-00728-0.

2. Sautes-Fridman, C., Petitprez, F., Calderaro, J., and Fridman, W.H. (2019). Tertiary lymphoid structures in the era of cancer immunotherapy. Nat Rev Cancer 19, 307–325. 10.1038/s41568-019-0144-6.

3. Schumacher, T.N., and Thommen, D.S. (2022). Tertiary lymphoid structures in cancer. Science 375, eabf9419. 10.1126/science.abf9419.

4. Ward, R.W., Kar, U., Balan, S., and Bhardwaj, N. (2026). Leveraging cDC1 biology and function for enhanced immunotherapy. J Exp Med 223. 10.1084/jem.20241199.

5. Peyraud, F., Guegan, J.P., Vanhersecke, L., Brunet, M., Teyssonneau, D., Palmieri, L.J., Bessede, A., and Italiano, A. (2025). Tertiary lymphoid structures and cancer immunotherapy: From bench to bedside. Med 6, 100546. 10.1016/j.medj.2024.10.023.

6. Mattiuz, R., Boumelha, J., Aerakis, E., Le Berichel, J., Hamon, P., Halasz, L., Vaidya, A., Soong, B.Y., Radkevich, E., Kim, H.M., et al. (2026). Dendritic cells control tertiary lymphoid structure development and maintenance in cancer. Science 393, eady1678. 10.1126/science.ady1678.

7. Helmink, B.A., Reddy, S.M., Gao, J., Zhang, S., Basar, R., Thakur, R., Yizhak, K., Sade-Feldman, M., Blando, J., Han, G., et al. (2020). B cells and tertiary lymphoid structures promote immunotherapy response. Nature 577, 549–555. 10.1038/s41586-019-1922-8.

8. Cabrita, R., Lauss, M., Sanna, A., Donia, M., Skaarup Larsen, M., Mitra, S., Johansson, I., Phung, B., Harbst, K., Vallon-Christersson, J., et al. (2020). Tertiary lymphoid structures improve immunotherapy and survival in melanoma. Nature 577, 561–565. 10.1038/s41586-019-1914-8.

9. Petitprez, F., de Reynies, A., Keung, E.Z., Chen, T.W., Sun, C.M., Calderaro, J., Jeng, Y.M., Hsiao, L.P., Lacroix, L., Bougouin, A., et al. (2020). B cells are associated with survival and immunotherapy response in sarcoma. Nature 577, 556–560. 10.1038/s41586-019-1906-8.

10. Vanhersecke, L., Brunet, M., Guegan, J.P., Rey, C., Bougouin, A., Cousin, S., Moulec, S.L., Besse, B., Loriot, Y., Larroquette, M., et al. (2021). Mature tertiary lymphoid structures predict immune checkpoint inhibitor efficacy in solid tumors independently of PD-L1 expression. Nat Cancer 2, 794–802. 10.1038/s43018-021-00232-6.

11. Johansson-Percival, A., and Ganss, R. (2021). Therapeutic Induction of Tertiary Lymphoid Structures in Cancer Through Stromal Remodeling. Front Immunol 12, 674375. 10.3389/fimmu.2021.674375.

12. Kim, H.M., Joglekar, T., Cascone, T., and Bruno, T.C. (2026). The future of tertiary lymphoid structures in cancer immunotherapy as biomarkers and therapeutic targets. Nat Cancer 7, 1021–1024. 10.1038/s43018-026-01188-1.

13. Slingerland, N., Runderkamp, E., and Thommen, D.S. (2026). Immune Niches in Cancer. Annu Rev Immunol 44, 553–582. 10.1146/annurev-immunol-082724-123843.

14. Chaib, M., Aminu, M., Herbrich, S., Arabi, M., Xuan, Y., Basi, A., Casasent, A., Macaluso, M.D., Hu, K.H., Gubin, M., et al. (2025). Macrophage-Dendritic Cell-T-Cell Tetrads Orchestrate Antitumor Immunity and Response to Checkpoint Blockade. bioRxiv. 10.64898/2025.12.24.696419.

15. Chen, J.H., Nieman, L.T., Spurrell, M., Jorgji, V., Elmelech, L., Richieri, P., Xu, K.H., Madhu, R., Parikh, M., Zamora, I., et al. (2024). Human lung cancer harbors spatially organized stem-immunity hubs associated with response to immunotherapy. Nat Immunol 25, 644–658. 10.1038/s41590-024-01792-2.

16. Jansen, C.S., Prokhnevska, N., and Kissick, H.T. (2019). The requirement for immune infiltration and organization in the tumor microenvironment for successful immunotherapy in prostate cancer. Urol Oncol 37, 543–555. 10.1016/j.urolonc.2018.10.011.

17. Jansen, C.S., Prokhnevska, N., Master, V.A., Sanda, M.G., Carlisle, J.W., Bilen, M.A., Cardenas, M., Wilkinson, S., Lake, R., Sowalsky, A.G., et al. (2019). An intra-tumoral niche maintains and differentiates stem-like CD8 T cells. Nature 576, 465–470. 10.1038/s41586-019-1836-5.

18. Papadas, A., Huang, Y., Cicala, A., Dou, Y., Fields, M., Gibbons, A., Hong, D., Lagal, D.J., Quintana, V., Rizo, A., et al. (2023). Emerging roles for tumor stroma in antigen presentation and anti-cancer immunity. Biochem Soc Trans 51, 2017–2028. 10.1042/BST20221083.

19. Devi-Marulkar, P., Fastenackels, S., Karapentiantz, P., Goc, J., Germain, C., Kaplon, H., Knockaert, S., Olive, D., Panouillot, M., Validire, P., et al. (2022). Regulatory T cells infiltrate the tumor-induced tertiary lymphoid structures and are associated with poor clinical outcome in NSCLC. Commun Biol 5, 1416. 10.1038/s42003-022-04356-y.

20. Vion, R., Roulleaux-Dugage, M., Flippot, R., Ouali, K., Rouanne, M., Clatot, F., Sellars, M., Champiat, S., Chaput, N., Massard, C., and Danlos, F.X. (2025). Induction of tertiary lymphoid structures in tumor microenvironment to improve anti-tumoral immune checkpoint blockade efficacy. Eur J Cancer 225, 115572. 10.1016/j.ejca.2025.115572.

21. Yamauchi, M., Barker, T.H., Gibbons, D.L., and Kurie, J.M. (2018). The fibrotic tumor stroma. J Clin Invest 128, 16–25. 10.1172/JCI93554.

22. Ho, W.J., Jaffee, E.M., and Zheng, L. (2020). The tumour microenvironment in pancreatic cancer - clinical challenges and opportunities. Nat Rev Clin Oncol 17, 527–540. 10.1038/s41571-020-0363-5.

23. Kung, H.C., Zheng, K.W., Zimmerman, J.W., and Zheng, L. (2025). The tumour microenvironment in pancreatic cancer - new clinical challenges, but more opportunities. Nat Rev Clin Oncol 22, 969–995. 10.1038/s41571-025-01077-z.

24. MacCarthy-Morrogh, L., and Martin, P. (2020). The hallmarks of cancer are also the hallmarks of wound healing. Sci Signal 13. 10.1126/scisignal.aay8690.

25. Wight, T.N. (2017). Provisional matrix: A role for versican and hyaluronan. Matrix Biol 60 - 61, 38-56. 10.1016/j.matbio.2016.12.001.

26. Mariathasan, S., Turley, S.J., Nickles, D., Castiglioni, A., Yuen, K., Wang, Y., Kadel, E.E., III, Koeppen, H., Astarita, J.L., Cubas, R., et al. (2018). TGFbeta attenuates tumour response to PD-L1 blockade by contributing to exclusion of T cells. Nature 554, 544–548. 10.1038/nature25501.

27. Cardoso, E.C., Lee, H., England, F.J., Cho, H., Lu, R., Varankar, S.S., Park, M.S., Rekhtman, N., Koo, B.K., Simons, B.D., et al. (2026). Early fibrotic niches establish tumour-permissive microenvironments. Nature 653, 254–264. 10.1038/s41586-026-10399-6.

28. Wight, T.N., Kang, I., Evanko, S.P., Harten, I.A., Chang, M.Y., Pearce, O.M.T., Allen, C.E., and Frevert, C.W. (2020). Versican-A Critical Extracellular Matrix Regulator of Immunity and Inflammation. Front Immunol 11, 512. 10.3389/fimmu.2020.00512.

29. Islam, S., and Watanabe, H. (2020). Versican: A Dynamic Regulator of the Extracellular Matrix. J Histochem Cytochem 68, 763–775. 10.1369/0022155420953922.

30. Papadas, A., and Asimakopoulos, F. (2020). Versican in the Tumor Microenvironment. Adv Exp Med Biol 1272, 55–72. 10.1007/978-3-030-48457-6_4.

31. Ricciardelli, C., Sakko, A.J., Ween, M.P., Russell, D.L., and Horsfall, D.J. (2009). The biological role and regulation of versican levels in cancer. Cancer Metastasis Rev 28, 233–245. 10.1007/s10555-009-9182-y.

32. Schmitt, M. (2016). Versican vs versikine: tolerance vs attack. Blood 128, 612–613. 10.1182/blood-2016-06-721092.

33. Wight, T.N., Kang, I., and Merrilees, M.J. (2014). Versican and the control of inflammation. Matrix Biol 35, 152–161. 10.1016/j.matbio.2014.01.015.

34. Wight, T.N., Kinsella, M.G., Evanko, S.P., Potter-Perigo, S., and Merrilees, M.J. (2014). Versican and the regulation of cell phenotype in disease. Biochim Biophys Acta 1840, 2441–2451. 10.1016/j.bbagen.2013.12.028.

35. Hope, C., Foulcer, S., Jagodinsky, J., Chen, S.X., Jensen, J.L., Patel, S., Leith, C., Maroulakou, I., Callander, N., Miyamoto, S., et al. (2016). Immunoregulatory roles of versican proteolysis in the myeloma microenvironment. Blood 128, 680–685. 10.1182/blood-2016-03-705780.

36. Hope, C., Emmerich, P.B., Papadas, A., Pagenkopf, A., Matkowskyj, K.A., Van De Hey, D.R., Payne, S.N., Clipson, L., Callander, N.S., Hematti, P., et al. (2017). Versican-Derived Matrikines Regulate Batf3-Dendritic Cell Differentiation and Promote T Cell Infiltration in Colorectal Cancer. J Immunol 199, 1933–1941. 10.4049/jimmunol.1700529.

37. Hirani, P., McDermott, J., Rajeeve, V., Cutillas, P.R., Jones, J.L., Pennington, D.J., Wight, T.N., Santamaria, S., Alonge, K.M., and Pearce, O.M.T. (2024). Versican Associates with Tumor Immune Phenotype and Limits T-cell Trafficking via Chondroitin Sulfate. Cancer Res Commun 4, 970–985. 10.1158/2767-9764.CRC-23-0548.

38. Emmerich, P.B., Qyli, T., Johnson, K.A., Chaudhuri, S., Clark, K.M., Verhagen, N.B., Depke, M.G., Clipson, L., Pasch, C.A., Papadas, A., et al. (2025). Stromal Versican Accumulation and Proteolysis Regulate the Infiltration of CD8(+) T Cells in Breast Cancer. Cancers (Basel) 17. 10.3390/cancers17091435.

39. Papadas, A., Deb, G., Cicala, A., Officer, A., Hope, C., Pagenkopf, A., Flietner, E., Morrow, Z.T., Emmerich, P., Wiesner, J., et al. (2022). Stromal remodeling regulates dendritic cell abundance and activity in the tumor microenvironment. Cell Rep 40, 111201. 10.1016/j.celrep.2022.111201.

40. Emmerich, P., Matkowskyj, K.A., McGregor, S., Kraus, S., Bischel, K., Qyli, T., Buehler, D., Pasch, C., Babiarz, C., Depke, M., et al. (2020). VCAN accumulation and proteolysis as predictors of T lymphocyte-excluded and permissive tumor microenvironments. Journal of Clinical Oncology 38, 3127–3127. 10.1200/JCO.2020.38.15_suppl.3127.

41. Deming, D.A., Kraus, S.G., Brand, J., Johnson, K.A., Abbott, D., Kratz, J., Turk, A.A., Emmerich, P., Carchman, E., Lubner, S.J., et al. (2026). Tumor Matrix Proteoglycan Accumulation and Processing Alter T-cell Effector Function and the Response to Immunotherapy in Patients with Oligometastatic Colorectal Cancer. Clin Cancer Res 32, 1707–1723. 10.1158/1078-0432.CCR-25-2780.

42. Maquart, F.X., Bellon, G., Pasco, S., and Monboisse, J.C. (2005). Matrikines in the regulation of extracellular matrix degradation. Biochimie 87, 353–360. 10.1016/j.biochi.2004.10.006.

43. Papadas, A., Arauz, G., Cicala, A., Wiesner, J., and Asimakopoulos, F. (2020). Versican and Versican-matrikines in Cancer Progression, Inflammation, and Immunity. J Histochem Cytochem 68, 871–885. 10.1369/0022155420937098.

44. Nandadasa, S., Foulcer, S., and Apte, S.S. (2014). The multiple, complex roles of versican and its proteolytic turnover by ADAMTS proteases during embryogenesis. Matrix Biol 35, 34–41. 10.1016/j.matbio.2014.01.005.

45. McCulloch, D.R., Nelson, C.M., Dixon, L.J., Silver, D.L., Wylie, J.D., Lindner, V., Sasaki, T., Cooley, M.A., Argraves, W.S., and Apte, S.S. (2009). ADAMTS metalloproteases generate active versican fragments that regulate interdigital web regression. Dev Cell 17, 687–698. 10.1016/j.devcel.2009.09.008.

46. Islam, S., Chuensirikulchai, K., Khummuang, S., Keratibumrungpong, T., Kongtawelert, P., Kasinrerk, W., Hatano, S., Nagamachi, A., Honda, H., and Watanabe, H. (2020). Accumulation of versican facilitates wound healing: Implication of its initial ADAMTS-cleavage site. Matrix Biol 87, 77–93. 10.1016/j.matbio.2019.10.006.

47. Nandadasa, S., Burin des Roziers, C., Koch, C., Tran-Lundmark, K., Dours-Zimmermann, M.T., Zimmermann, D.R., Valleix, S., and Apte, S.S. (2021). A new mouse mutant with cleavage-resistant versican and isoform-specific versican mutants demonstrate that proteolysis at the Glu(441)-Ala(442) peptide bond in the V1 isoform is essential for interdigital web regression. Matrix Biol Plus 10, 100064. 10.1016/j.mbplus.2021.100064.

48. Deb, G., Cicala, A., Papadas, A., and Asimakopoulos, F. (2022). Matrix proteoglycans in tumor inflammation and immunity. Am J Physiol Cell Physiol 323, C678–C693. 10.1152/ajpcell.00023.2022.

49. Mueller, C.G., Nayar, S., Campos, J., and Barone, F. (2018). Molecular and Cellular Requirements for the Assembly of Tertiary Lymphoid Structures. Adv Exp Med Biol 1060, 55–72. 10.1007/978-3-319-78127-3_4.

50. Espinosa-Carrasco, G., Chiu, E., Scrivo, A., Zumbo, P., Dave, A., Betel, D., Kang, S.W., Jang, H.J., Hellmann, M.D., Burt, B.M., et al. (2024). Intratumoral immune triads are required for immunotherapy-mediated elimination of solid tumors. Cancer Cell 42, 1202–1216 e1208. 10.1016/j.ccell.2024.05.025.

51. Minns, A.F., and Santamaria, S. (2024). Determination of Versikine Levels by Enzyme-Linked Immunosorbent Assay (ELISA). Methods Mol Biol 2747, 83–93. 10.1007/978-1-0716-3589-6_8.

52. Zheng, G.X., Terry, J.M., Belgrader, P., Ryvkin, P., Bent, Z.W., Wilson, R., Ziraldo, S.B., Wheeler, T.D., McDermott, G.P., Zhu, J., et al. (2017). Massively parallel digital transcriptional profiling of single cells. Nat Commun 8, 14049. 10.1038/ncomms14049.

53. Hao, Y., Stuart, T., Kowalski, M.H., Choudhary, S., Hoffman, P., Hartman, A., Srivastava, A., Molla, G., Madad, S., Fernandez-Granda, C., and Satija, R. (2024). Dictionary learning for integrative, multimodal and scalable single-cell analysis. Nat Biotechnol 42, 293–304. 10.1038/s41587-023-01767-y.

54. Andreatta, M., Corria-Osorio, J., Muller, S., Cubas, R., Coukos, G., and Carmona, S.J. (2021). Interpretation of T cell states from single-cell transcriptomics data using reference atlases. Nat Commun 12, 2965. 10.1038/s41467-021-23324-4.

55. Merad, M., Sathe, P., Helft, J., Miller, J., and Mortha, A. (2013). The dendritic cell lineage: ontogeny and function of dendritic cells and their subsets in the steady state and the inflamed setting. Annu Rev Immunol 31, 563–604. 10.1146/annurev-immunol-020711-074950.

56. Subramanian, A., Tamayo, P., Mootha, V.K., Mukherjee, S., Ebert, B.L., Gillette, M.A., Paulovich, A., Pomeroy, S.L., Golub, T.R., Lander, E.S., and Mesirov, J.P. (2005). Gene set enrichment analysis: a knowledge-based approach for interpreting genome-wide expression profiles. Proc Natl Acad Sci U S A 102, 15545–15550. 10.1073/pnas.0506580102.

57. Andreatta, M., and Carmona, S.J. (2021). UCell: Robust and scalable single-cell gene signature scoring. Comput Struct Biotechnol J 19, 3796–3798. 10.1016/j.csbj.2021.06.043.

58. Jin, S., Plikus, M.V., and Nie, Q. (2025). CellChat for systematic analysis of cell-cell communication from single-cell transcriptomics. Nat Protoc 20, 180–219. 10.1038/s41596-024-01045-4.

59. Takei, S., Yamasaki, S., Yamaguchi, O., Mouri, A., Shiono, A., Miura, Y., Hashimoto, K., Imai, H., Kaira, K., Ichiki, Y., et al. (2026). The CD4(+) T cell population partners with Tpex CD8(+) T cells to mediate antitumor immunity in the tumor microenvironment. Nat Commun 17 10.1038/s41467-026-71161-0.

60. Hao, Y., Hao, S., Andersen-Nissen, E., Mauck, W.M., 3rd, Zheng, S., Butler, A., Lee, M.J., Wilk, A.J., Darby, C., Zager, M., et al. (2021). Integrated analysis of multimodal single-cell data. Cell 184, 3573–3587 e3529. 10.1016/j.cell.2021.04.048.

61. Badia, I.M.P., Velez Santiago, J., Braunger, J., Geiss, C., Dimitrov, D., Muller-Dott, S., Taus, P., Dugourd, A., Holland, C.H., Ramirez Flores, R.O., and Saez-Rodriguez, J. (2022). decoupleR: ensemble of computational methods to infer biological activities from omics data. Bioinform Adv 2, vbac016. 10.1093/bioadv/vbac016.

62. Muller-Dott, S., Tsirvouli, E., Vazquez, M., Ramirez Flores, R.O., Badia, I.M.P., Fallegger, R., Turei, D., Laegreid, A., and Saez-Rodriguez, J. (2023). Expanding the coverage of regulons from high-confidence prior knowledge for accurate estimation of transcription factor activities. Nucleic Acids Res 51, 10934–10949. 10.1093/nar/gkad841.

63. Hope, C., Ollar, S.J., Heninger, E., Hebron, E., Jensen, J.L., Kim, J., Maroulakou, I., Miyamoto, S., Leith, C., Yang, D.T., et al. (2014). TPL2 kinase regulates the inflammatory milieu of the myeloma niche. Blood 123, 3305–3315. 10.1182/blood-2014-02-554071.

64. Timms, K.P., and Maurice, S.B. (2020). Context-dependent bioactivity of versican fragments. Glycobiology 30, 365–373. 10.1093/glycob/cwz090.

65. Py, B.F., Gonzalez, S.F., Long, K., Kim, M.S., Kim, Y.A., Zhu, H., Yao, J., Degauque, N., Villet, R., Ymele-Leki, P., et al. (2013). Cochlin produced by follicular dendritic cells promotes antibacterial innate immunity. Immunity 38, 1063–1072. 10.1016/j.immuni.2013.01.015.

66. Allen, C.D., Okada, T., and Cyster, J.G. (2007). Germinal-center organization and cellular dynamics. Immunity 27, 190–202. 10.1016/j.immuni.2007.07.009.

67. Gutierrez-Melo, N., and Baumjohann, D. (2023). T follicular helper cells in cancer. Trends Cancer 9, 309–325. 10.1016/j.trecan.2022.12.007.

68. Carlsen, H.S., Baekkevold, E.S., Morton, H.C., Haraldsen, G., and Brandtzaeg, P. (2004). Monocyte-like and mature macrophages produce CXCL13 (B cell-attracting chemokine 1) in inflammatory lesions with lymphoid neogenesis. Blood 104, 3021–3027. 10.1182/blood-2004-02-0701.

69. Ukita, M., Hamanishi, J., Yoshitomi, H., Yamanoi, K., Takamatsu, S., Ueda, A., Suzuki, H., Hosoe, Y., Furutake, Y., Taki, M., et al. (2022). CXCL13-producing CD4+ T cells accumulate in the early phase of tertiary lymphoid structures in ovarian cancer. JCI Insight 7. 10.1172/jci.insight.157215.

70. Gu, X., Li, D., Wu, P., Zhang, C., Cui, X., Shang, D., Ma, R., Liu, J., Sun, N., and He, J. (2024). Revisiting the CXCL13/CXCR5 axis in the tumor microenvironment in the era of single-cell omics: Implications for immunotherapy. Cancer Lett 605, 217278. 10.1016/j.canlet.2024.217278.

71. Di Pietro, A., Au, L., Crock, P., Thio, N., Pizzolla, A., Nguyen, T.N., Macdonald, S., Chalmers, H., Zhu, R., Airaghi, A., et al. (2026). Tumor-resident T cells and dendritic cells form an in situ archetype during immunotherapy response in melanoma. Nat Commun. 10.1038/s41467-026-74076-y.

72. Moon, C.Y., Belabed, M., Park, M.D., Mattiuz, R., Puleston, D., and Merad, M. (2025). Dendritic cell maturation in cancer. Nat Rev Cancer 25, 225–248. 10.1038/s41568-024-00787-3.

73. Magen, A., Hamon, P., Fiaschi, N., Soong, B.Y., Park, M.D., Mattiuz, R., Humblin, E., Troncoso, L., D’Souza, D., Dawson, T., et al. (2023). Intratumoral dendritic cell-CD4(+) T helper cell niches enable CD8(+) T cell differentiation following PD-1 blockade in hepatocellular carcinoma. Nat Med 29, 1389–1399. 10.1038/s41591-023-02345-0.

74. Lei, X., Khatri, I., de Wit, T., de Rink, I., Nieuwland, M., Kerkhoven, R., van Eenennaam, H., Sun, C., Garg, A.D., Borst, J., and Xiao, Y. (2023). CD4(+) helper T cells endow cDC1 with cancer-impeding functions in the human tumor micro-environment. Nat Commun 14, 217. 10.1038/s41467-022-35615-5.

75. Ferris, S.T., Durai, V., Wu, R., Theisen, D.J., Ward, J.P., Bern, M.D., Davidson, J.T.t., Bagadia, P., Liu, T., Briseno, C.G., et al. (2020). cDC1 prime and are licensed by CD4(+) T cells to induce anti-tumour immunity. Nature 584, 624–629. 10.1038/s41586-020-2611-3.

76. Damle, S.R., Carter, J.A., Goodsell, K.E., Pineda, J.M.B., Dickerson, L.K., Jiang, X., Mudd, J.L., Walsh, T., Kenerson, H.L., Cernak, J., et al. (2026). Intratumoral Three-Cell-Type Clusters Are a Conserved Feature of Endogenous Antitumor Immunity. Cancer Immunology Research 14, 205–218. 10.1158/2326-6066.Cir-25-0062.

77. Meiser, P., Knolle, M.A., Hirschberger, A., de Almeida, G.P., Bayerl, F., Lacher, S., Pedde, A.M., Flommersfeld, S., Honninger, J., Stark, L., et al. (2023). A distinct stimulatory cDC1 subpopulation amplifies CD8(+) T cell responses in tumors for protective anti-cancer immunity. Cancer Cell 41, 1498–1515 e1410. 10.1016/j.ccell.2023.06.008.

78. Papadas, A., Lagal, D.J., Dou, Y., Hong, D., Gibbons, A., Cicala, A., Huang, Y., Zomalan, B., Molina, E., and Asimakopoulos, F. (2024). Protocol to identify and isolate rare murine tumor-resident dendritic cell populations for low-input transcriptomic profiling. STAR Protoc 5, 103195. 10.1016/j.xpro.2024.103195.

79. Hildner, K., Edelson, B.T., Purtha, W.E., Diamond, M., Matsushita, H., Kohyama, M., Calderon, B., Schraml, B.U., Unanue, E.R., Diamond, M.S., et al. (2008). Batf3 deficiency reveals a critical role for CD8alpha+ dendritic cells in cytotoxic T cell immunity. Science 322, 1097–1100. 10.1126/science.1164206.

80. Ye, X., Waite, J.C., Dhanik, A., Gupta, N., Zhong, M., Adler, C., Malahias, E., Ni, M., Wei, Y., Gurer, C., et al. (2020). Endogenous retroviral proteins provide an immunodominant but not requisite antigen in a murine immunotherapy tumor model. Oncoimmunology 9, 1758602. 10.1080/2162402X.2020.1758602.

81. Ahmed, R., Hofmann, M., Kallies, A., Oxenius, A., Philip, M., Wherry, E.J., Ye, L., and Zehn, D. (2026). New insights into progenitor exhausted T cell populations. Nat Rev Immunol. 10.1038/s41577-026-01328-9.

82. Dahling, S., Mansilla, A.M., Knopper, K., Grafen, A., Utzschneider, D.T., Ugur, M., Whitney, P.G., Bachem, A., Arampatzi, P., Imdahl, F., et al. (2022). Type 1 conventional dendritic cells maintain and guide the differentiation of precursors of exhausted T cells in distinct cellular niches. Immunity 55, 656–670 e658. 10.1016/j.immuni.2022.03.006.

83. Taylor, M.A., Hughes, A.M., Walton, J., Coenen-Stass, A.M.L., Magiera, L., Mooney, L., Bell, S., Staniszewska, A.D., Sandin, L.C., Barry, S.T., et al. (2019). Longitudinal immune characterization of syngeneic tumor models to enable model selection for immune oncology drug discovery. J Immunother Cancer 7, 328. 10.1186/s40425-019-0794-7.

84. Chow, M.T., Ozga, A.J., Servis, R.L., Frederick, D.T., Lo, J.A., Fisher, D.E., Freeman, G.J., Boland, G.M., and Luster, A.D. (2019). Intratumoral Activity of the CXCR3 Chemokine System Is Required for the Efficacy of Anti-PD-1 Therapy. Immunity 50, 1498–1512 e1495. 10.1016/j.immuni.2019.04.010.

85. Ley, K. (2014). The second touch hypothesis: T cell activation, homing and polarization. F1000Res 3, 37. 10.12688/f1000research.3-37.v2.

86. Romano, E., Kusio-Kobialka, M., Foukas, P.G., Baumgaertner, P., Meyer, C., Ballabeni, P., Michielin, O., Weide, B., Romero, P., and Speiser, D.E. (2015). Ipilimumab-dependent cell-mediated cytotoxicity of regulatory T cells ex vivo by nonclassical monocytes in melanoma patients. Proc Natl Acad Sci U S A 112, 6140–6145. 10.1073/pnas.1417320112.

87. Maier, B., Leader, A.M., Chen, S.T., Tung, N., Chang, C., LeBerichel, J., Chudnovskiy, A., Maskey, S., Walker, L., Finnigan, J.P., et al. (2020). A conserved dendritic-cell regulatory program limits antitumour immunity. Nature 580, 257–262. 10.1038/s41586-020-2134-y.

88. Foulcer, S.J., Day, A.J., and Apte, S.S. (2015). Isolation and purification of versican and analysis of versican proteolysis. Methods Mol Biol 1229, 587–604. 10.1007/978-1-4939-1714-3_46.

89. Takei, S., Shiono, A., Yamasaki, S., Yamaguchi, O., Mouri, A., Miura, Y., Hashimoto, K., Imai, H., Kaira, K., and Kagamu, H. (2025). Th7R predicts chemo-immunotherapy response and survival in small cell lung cancer. Cancer Immunol Immunother 75, 17. 10.1007/s00262-025-04242-6.

90. Damle, S.R., Carter, J.A., Goodsell, K.E., Pineda, J.M.B., Dickerson, L.K., Jiang, X., Mudd, J.L., Walsh, T., Kenerson, H.L., Cernak, J., et al. (2026). Intratumoral Three-Cell-Type Clusters Are a Conserved Feature of Endogenous Antitumor Immunity. Cancer Immunol Res 14, 205–218. 10.1158/2326-6066.CIR-25-0062.

91. Barker, T.H., and Engler, A.J. (2017). The provisional matrix: setting the stage for tissue repair outcomes. Matrix Biol 60-61, 1-4. 10.1016/j.matbio.2017.04.003.

92. Islam, S., Jahan, N., Shahida, A., Karnan, S., and Watanabe, H. (2022). Accumulation of versican and lack of versikine ameliorate acute colitis. Matrix Biol 107, 59–76. 10.1016/j.matbio.2022.02.004.

93. McMahon, M., Ye, S., Izzard, L., Dlugolenski, D., Tripp, R.A., Bean, A.G., McCulloch, D.R., and Stambas, J. (2016). ADAMTS5 Is a Critical Regulator of Virus-Specific T Cell Immunity. PLoS Biol 14, e1002580. 10.1371/journal.pbio.1002580.

94. Duong, E., Fessenden, T.B., Lutz, E., Dinter, T., Yim, L., Blatt, S., Bhutkar, A., Wittrup, K.D., and Spranger, S. (2022). Type I interferon activates MHC class I-dressed CD11b(+) conventional dendritic cells to promote protective anti-tumor CD8(+) T cell immunity. Immunity 55, 308–323 e309. 10.1016/j.immuni.2021.10.020.

95. Cui, C., Craft, J., and Joshi, N.S. (2023). T follicular helper cells in cancer, tertiary lymphoid structures, and beyond. Semin Immunol 69, 101797. 10.1016/j.smim.2023.101797.

96. Garaud, S., Dieu-Nosjean, M.C., and Willard-Gallo, K. (2022). T follicular helper and B cell crosstalk in tertiary lymphoid structures and cancer immunotherapy. Nat Commun 13, 2259. 10.1038/s41467-022-29753-z.

97. Yanagihara, A., Yamasaki, S., Hashimoto, K., Taguchi, R., Umesaki, T., Imai, H., Kaira, K., Nitanda, H., Sakaguchi, H., Ishida, H., et al. (2023). A Th1-like CD4(+) T-cell Cluster That Predicts Disease-free Survival in Early-stage Lung Cancer. Cancer Res Commun 3, 1277–1285. 10.1158/2767-9764.CRC-23-0167.

98. Noel, G., Fontsa, M.L., Garaud, S., De Silva, P., de Wind, A., Van den Eynden, G.G., Salgado, R., Boisson, A., Locy, H., Thomas, N., et al. (2021). Functional Th1-oriented T follicular helper cells that infiltrate human breast cancer promote effective adaptive immunity. J Clin Invest 131. 10.1172/JCI139905.

99. Moreno Ayala, M.A., Campbell, T.F., Zhang, C., Dahan, N., Bockman, A., Prakash, V., Feng, L., Sher, T., and DuPage, M. (2023). CXCR3 expression in regulatory T cells drives interactions with type I dendritic cells in tumors to restrict CD8(+) T cell antitumor immunity. Immunity 56, 1613–1630 e1615. 10.1016/j.immuni.2023.06.003.

100. Zagorulya, M., Yim, L., Morgan, D.M., Edwards, A., Torres-Mejia, E., Momin, N., McCreery, C.V., Zamora, I.L., Horton, B.L., Fox, J.G., et al. (2023). Tissue-specific abundance of interferon-gamma drives regulatory T cells to restrain DC1-mediated priming of cytotoxic T cells against lung cancer. Immunity 56, 386–405 e310. 10.1016/j.immuni.2023.01.010.

101. Tang, M., Diao, J., and Cattral, M.S. (2017). Molecular mechanisms involved in dendritic cell dysfunction in cancer. Cell Mol Life Sci 74, 761–776. 10.1007/s00018-016-2317-8.

102. Tang, M., Diao, J., Gu, H., Khatri, I., Zhao, J., and Cattral, M.S. (2015). Toll-like Receptor 2 Activation Promotes Tumor Dendritic Cell Dysfunction by Regulating IL-6 and IL-10 Receptor Signaling. Cell Rep 13, 2851–2864. 10.1016/j.celrep.2015.11.053.

103. Moreth, K., Brodbeck, R., Babelova, A., Gretz, N., Spieker, T., Zeng-Brouwers, J., Pfeilschifter, J., Young, M.F., Schaefer, R.M., and Schaefer, L. (2010). The proteoglycan biglycan regulates expression of the B cell chemoattractant CXCL13 and aggravates murine lupus nephritis. J Clin Invest 120, 4251–4272. 10.1172/JCI42213.

104. Watanabe, H. (2023). Versican and versikine: the dynamism of the extracellular matrix. Proteoglycan Research 1, e13.

105. Davis, S.J., and van der Merwe, P.A. (2006). The kinetic-segregation model: TCR triggering and beyond. Nat Immunol 7, 803–809. 10.1038/ni1369.

106. Crocker, P.R., Paulson, J.C., and Varki, A. (2007). Siglecs and their roles in the immune system. Nat Rev Immunol 7, 255–266. 10.1038/nri2056.

107. Asano, K., Nelson, C.M., Nandadasa, S., Aramaki-Hattori, N., Lindner, D.J., Alban, T., Inagaki, J., Ohtsuki, T., Oohashi, T., Apte, S.S., and Hirohata, S. (2017). Stromal Versican Regulates Tumor Growth by Promoting Angiogenesis. Sci Rep 7, 17225. 10.1038/s41598-017-17613-6.

108. Rodriguez-Baena, F.J., Redondo-Garcia, S., Peris-Torres, C., Martino-Echarri, E., Fernandez-Rodriguez, R., Plaza-Calonge, M.D.C., Anderson, P., and Rodriguez-Manzaneque, J.C. (2018). ADAMTS1 protease is required for a balanced immune cell repertoire and tumour inflammatory response. Sci Rep 8, 13103. 10.1038/s41598-018-31288-7.

109. Wolters, A., Bijlsma, M., and Prakash, J. (2026). Precision targeting of stromal states in pancreatic cancer: a clinical perspective. Trends Cancer. 10.1016/j.trecan.2026.07.003.

110. Sahai, E., Astsaturov, I., Cukierman, E., DeNardo, D.G., Egeblad, M., Evans, R.M., Fearon, D., Greten, F.R., Hingorani, S.R., Hunter, T., et al. (2020). A framework for advancing our understanding of cancer-associated fibroblasts. Nat Rev Cancer 20, 174–186. 10.1038/s41568-019-0238-1.

111. Catenacci, D.V., Junttila, M.R., Karrison, T., Bahary, N., Horiba, M.N., Nattam, S.R., Marsh, R., Wallace, J., Kozloff, M., Rajdev, L., et al. (2015). Randomized Phase Ib/II Study of Gemcitabine Plus Placebo or Vismodegib, a Hedgehog Pathway Inhibitor, in Patients With Metastatic Pancreatic Cancer. J Clin Oncol 33, 4284–4292. 10.1200/JCO.2015.62.8719.

112. Van Cutsem, E., Tempero, M.A., Sigal, D., Oh, D.Y., Fazio, N., Macarulla, T., Hitre, E., Hammel, P., Hendifar, A.E., Bates, S.E., et al. (2020). Randomized Phase III Trial of Pegvorhyaluronidase Alfa With Nab-Paclitaxel Plus Gemcitabine for Patients With Hyaluronan-High Metastatic Pancreatic Adenocarcinoma. J Clin Oncol 38, 3185–3194. 10.1200/JCO.20.00590.

