## Supplementary material for "A matrikine organizes the dendritic cell–T cell triads that build tertiary lymphoid structures": Supp. Fig. S

Supplementary Figure S1

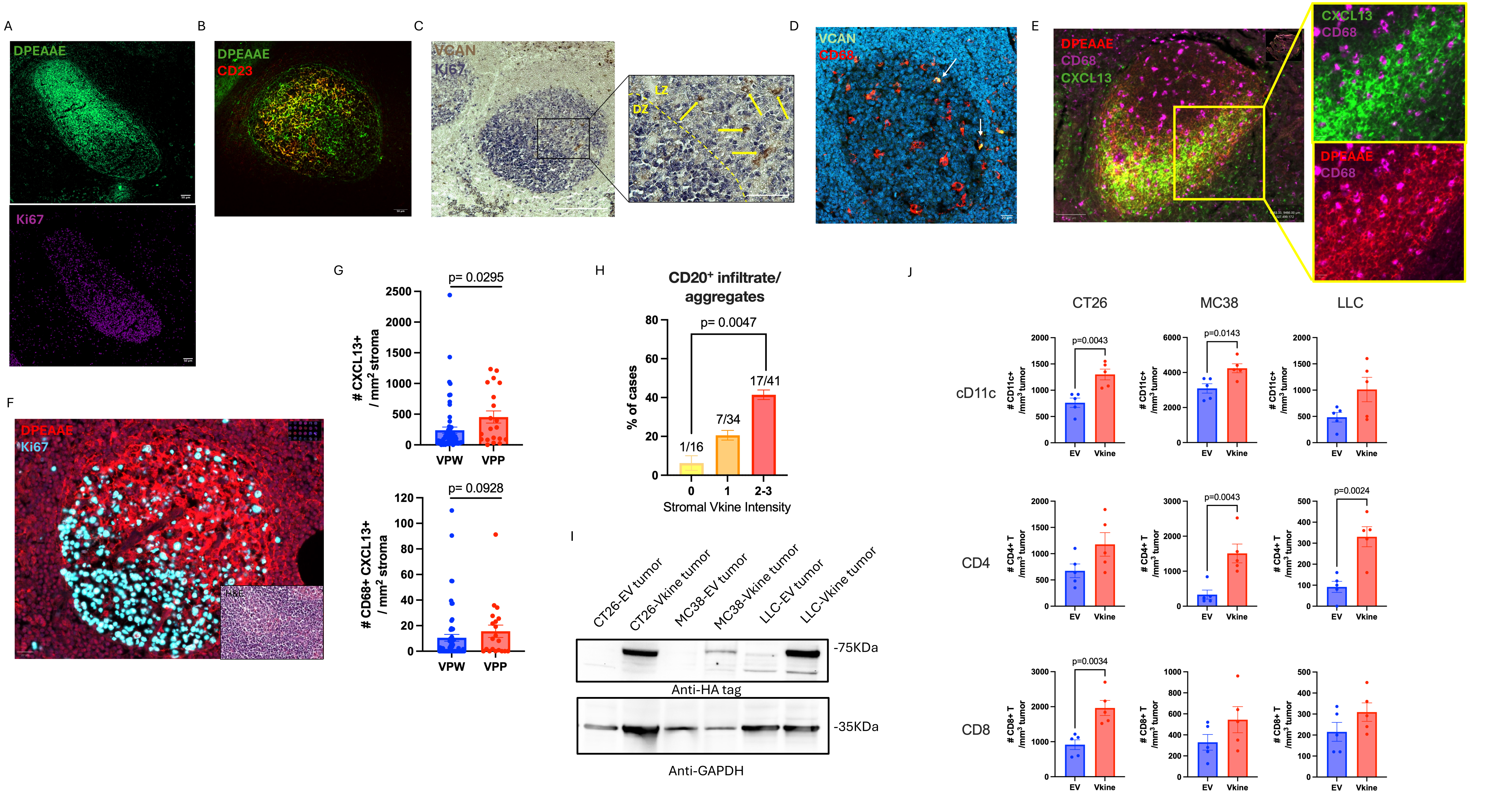

Supplementary Figure S2

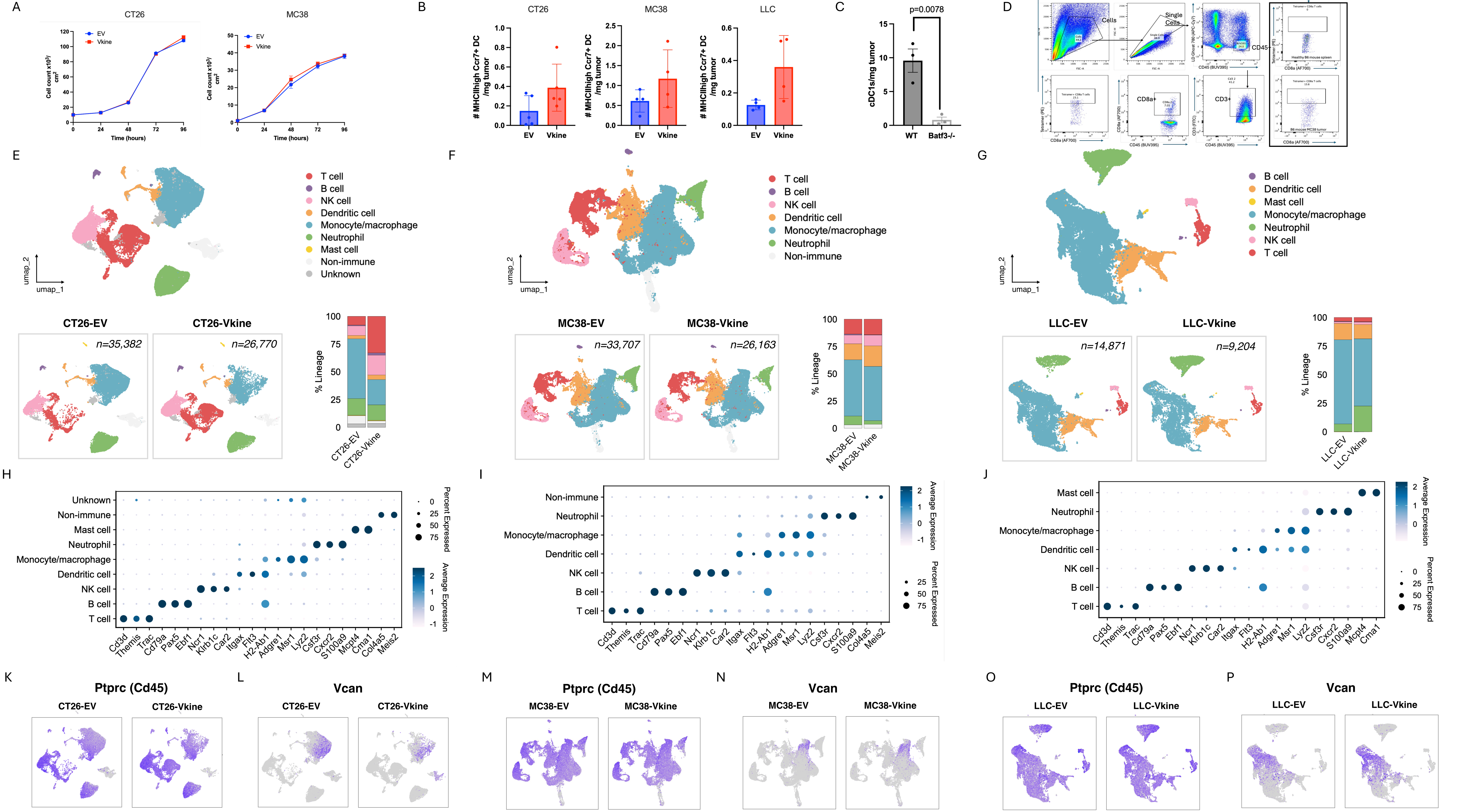

Supplementary Figure S3

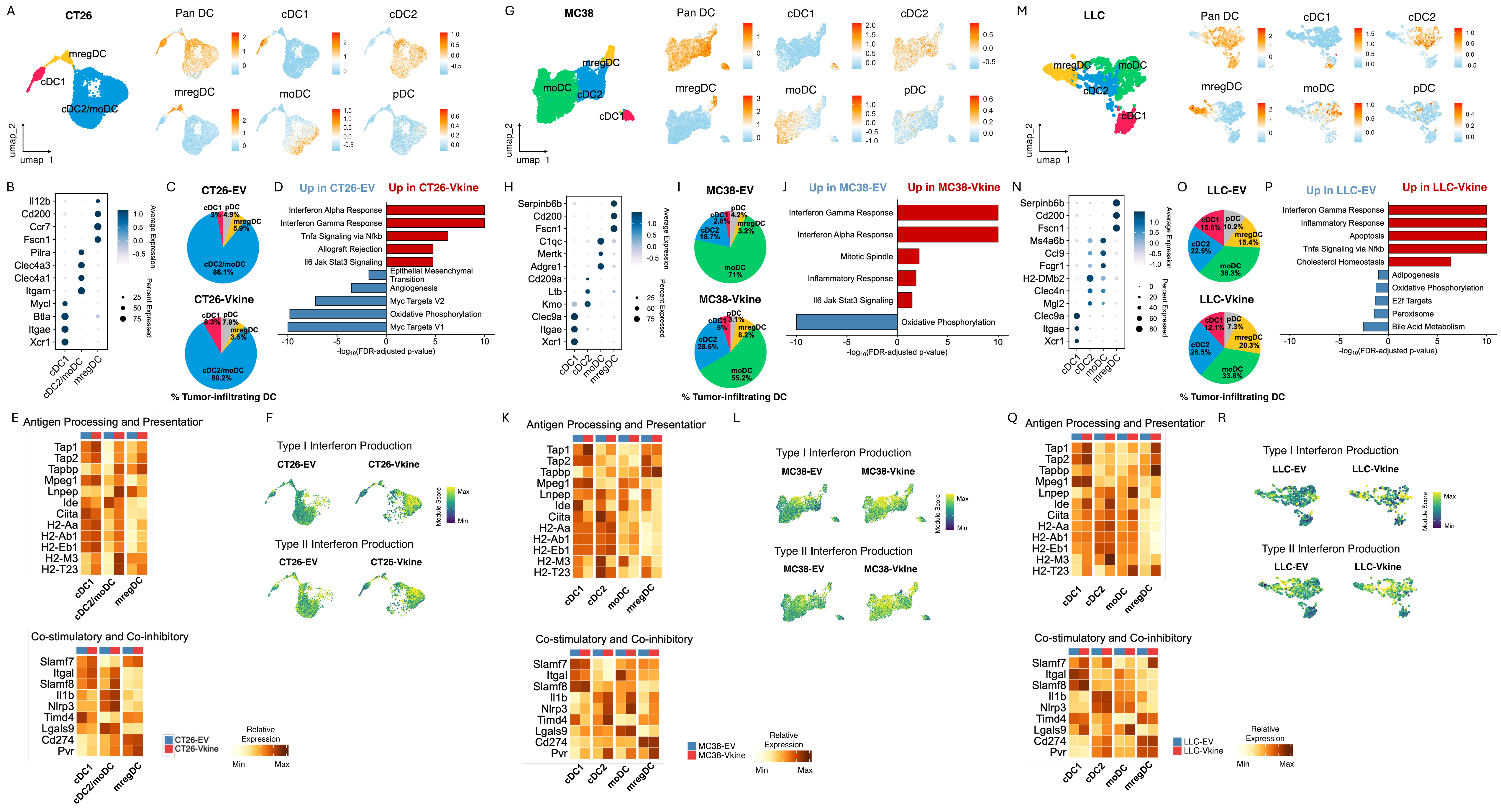

Supplementary Figure S4

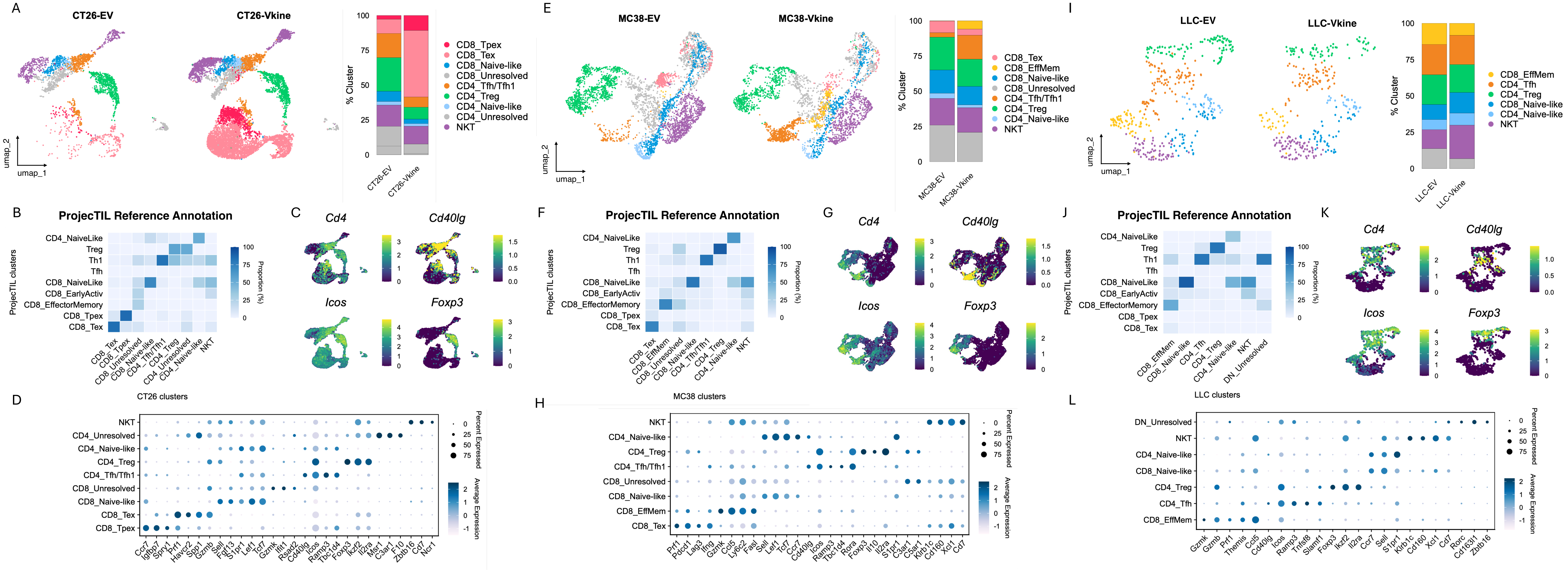

Supplementary Figure S5

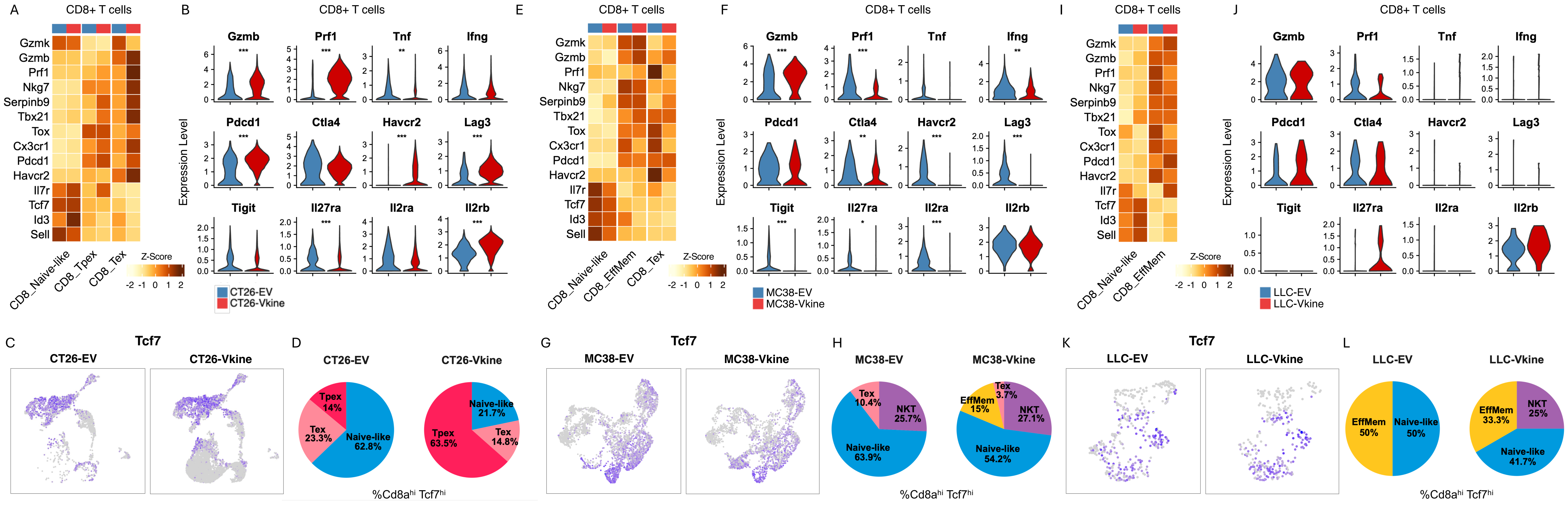

Supplementary Figure S6

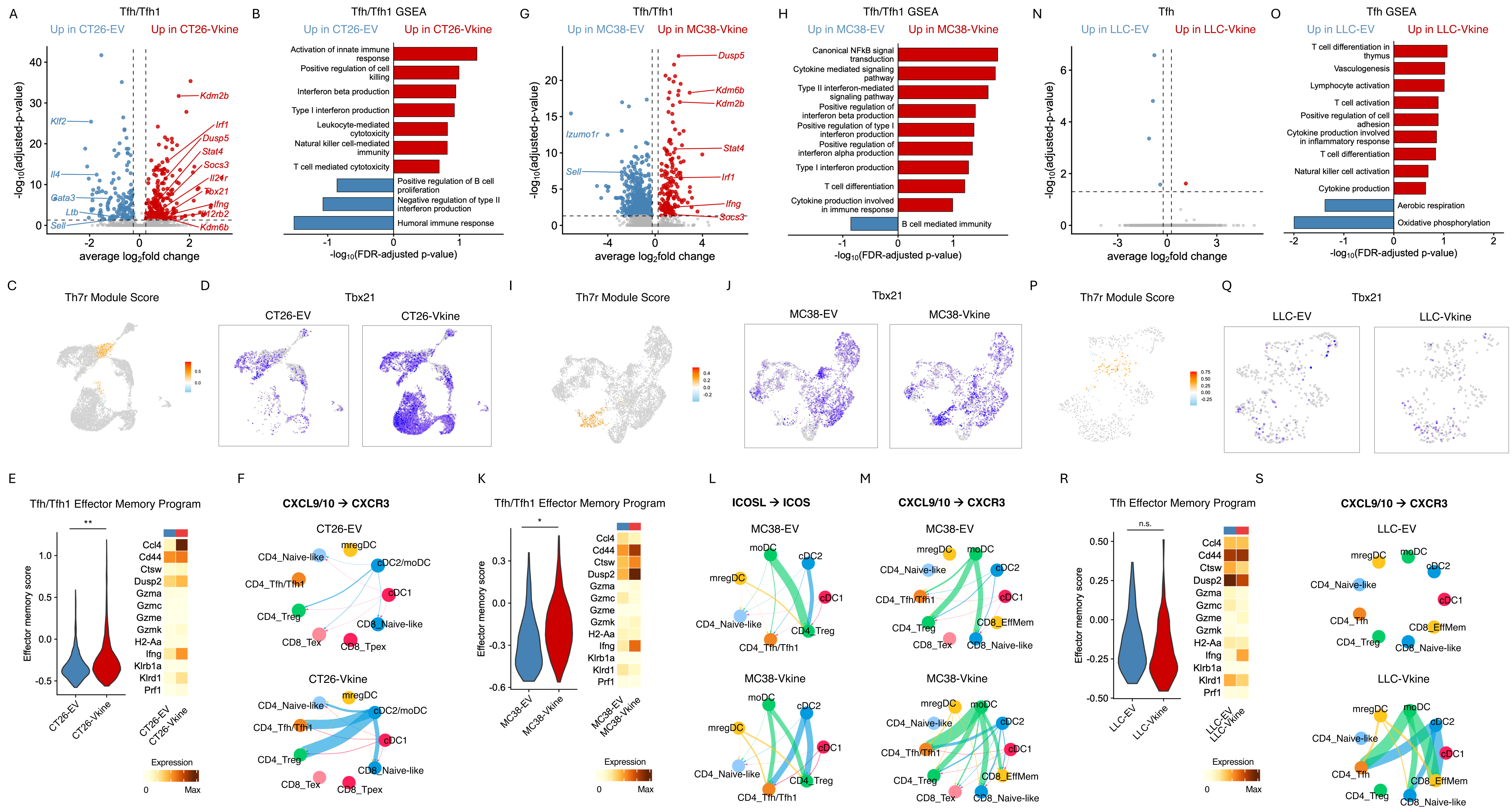

Supplementary Figure S7

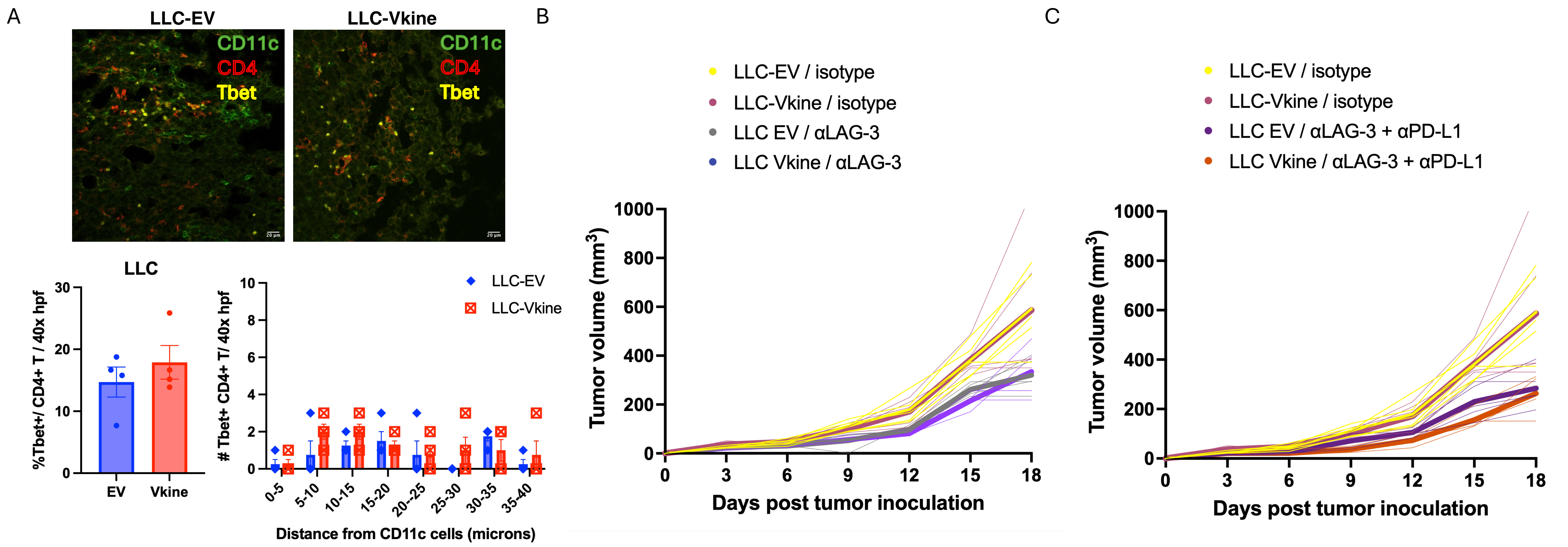

Supplementary Figure S8

A

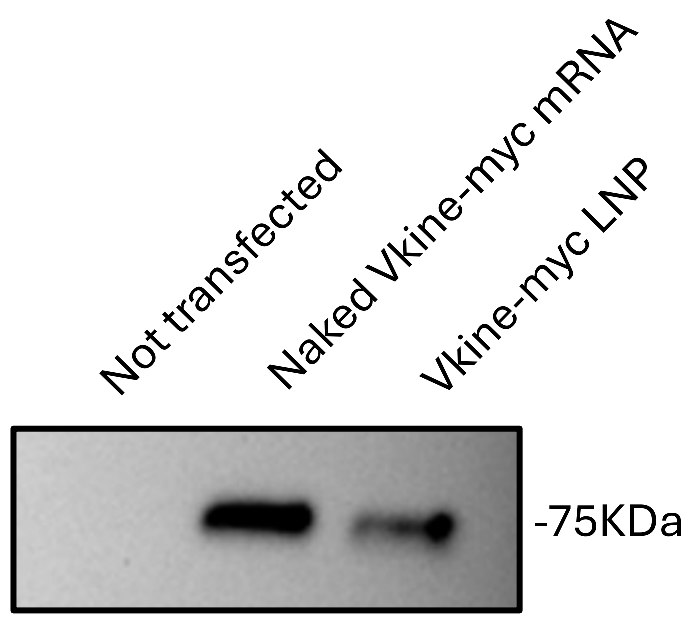


B

| | | | Size (d.nm): | % Number: | St Dev (d.n... |
|---|---|---|---|---|---|
| Z-Average (d.nm): | 85.15 | Peak 1: | 64.52 | 100.0 | 15.52 |
| PdI: | 0.082 | Peak 2: | 0.000 | 0.0 | 0.000 |
| Intercept: | 0.933 | Peak 3: | 0.000 | 0.0 | 0.000 |
| Result quality : | Good | | | | |

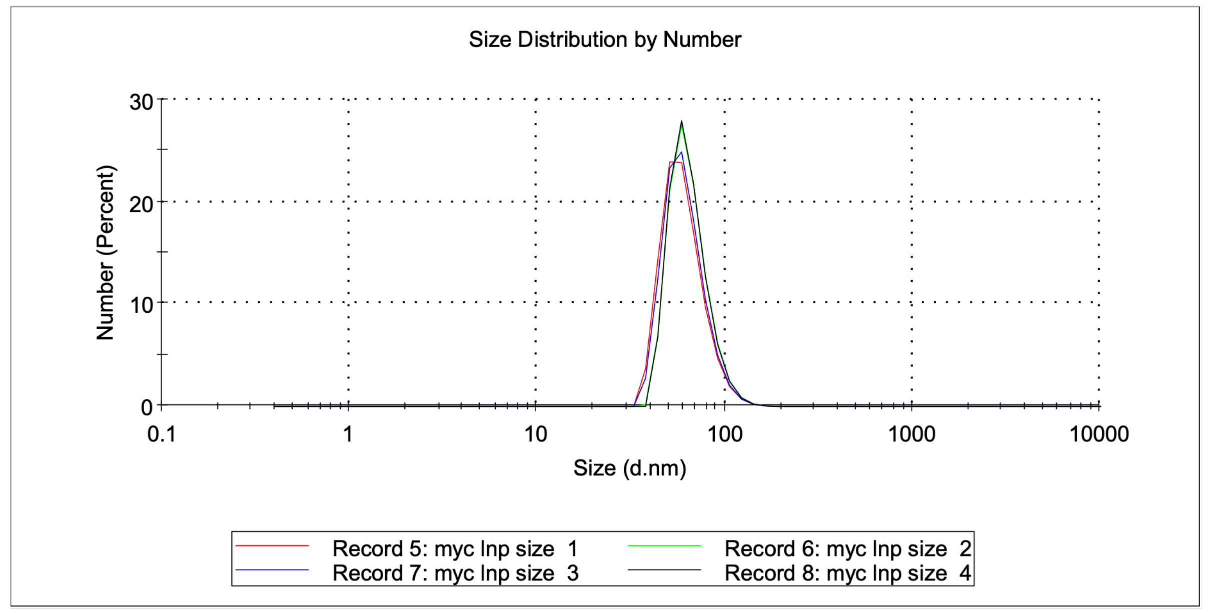


| | | | Mean (mV) | Area (%) | St Dev (mV) |
|---|---|---|---|---|---|
| Zeta Potential (mV): | 2.74 | Peak 1: | 2.74 | 100.0 | 8.48 |
| Zeta Deviation (mV): | 8.48 | Peak 2: | 0.00 | 0.0 | 0.00 |
| Conductivity (mS/cm): | 0.423 | Peak 3: | 0.00 | 0.0 | 0.00 |
| Result quality : | Good | | | | |

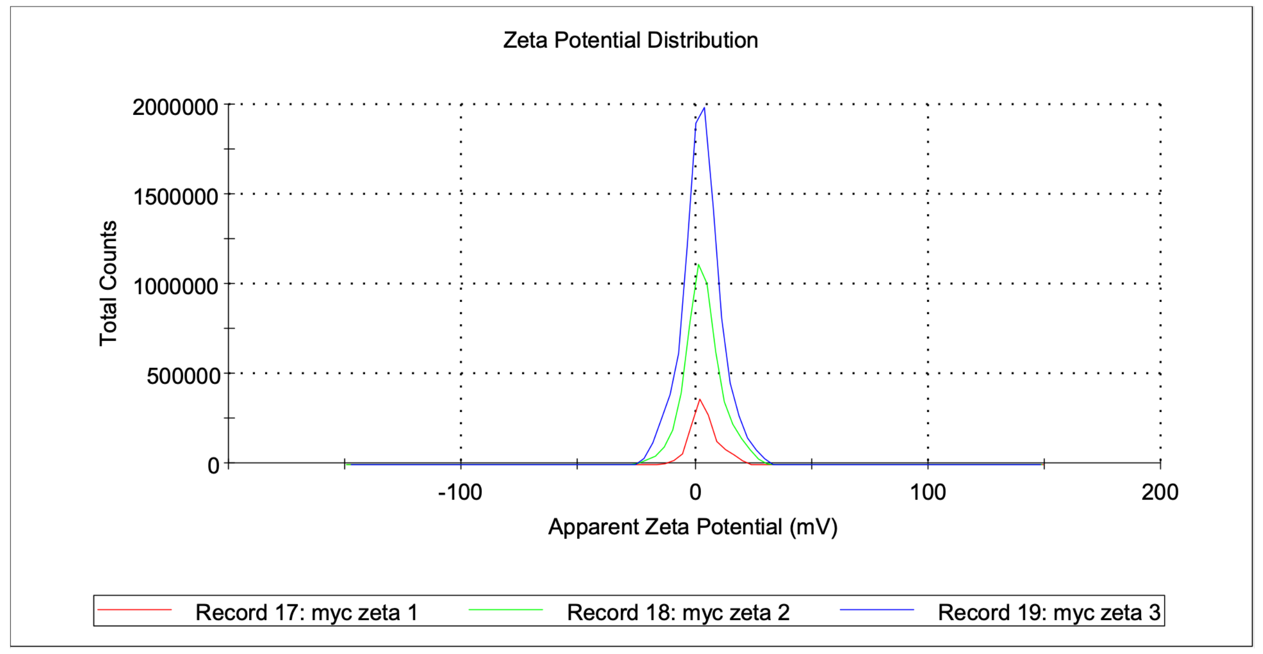


C

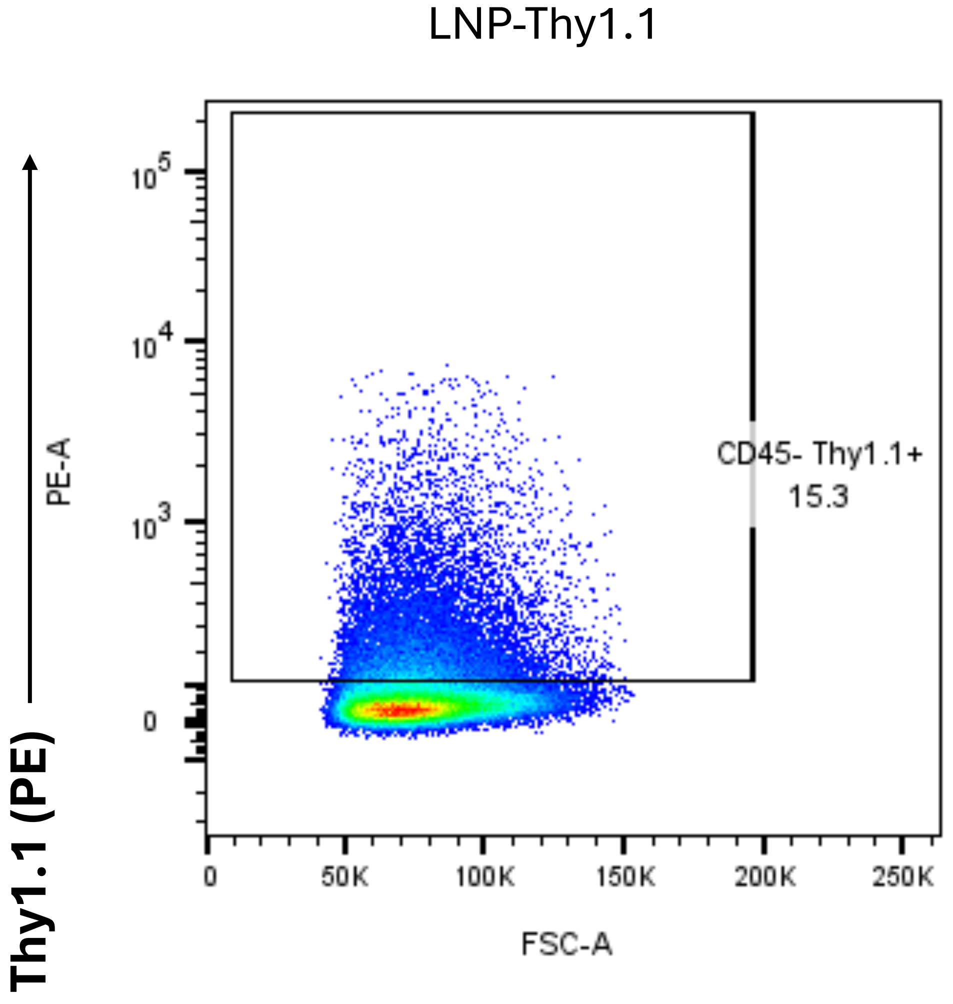


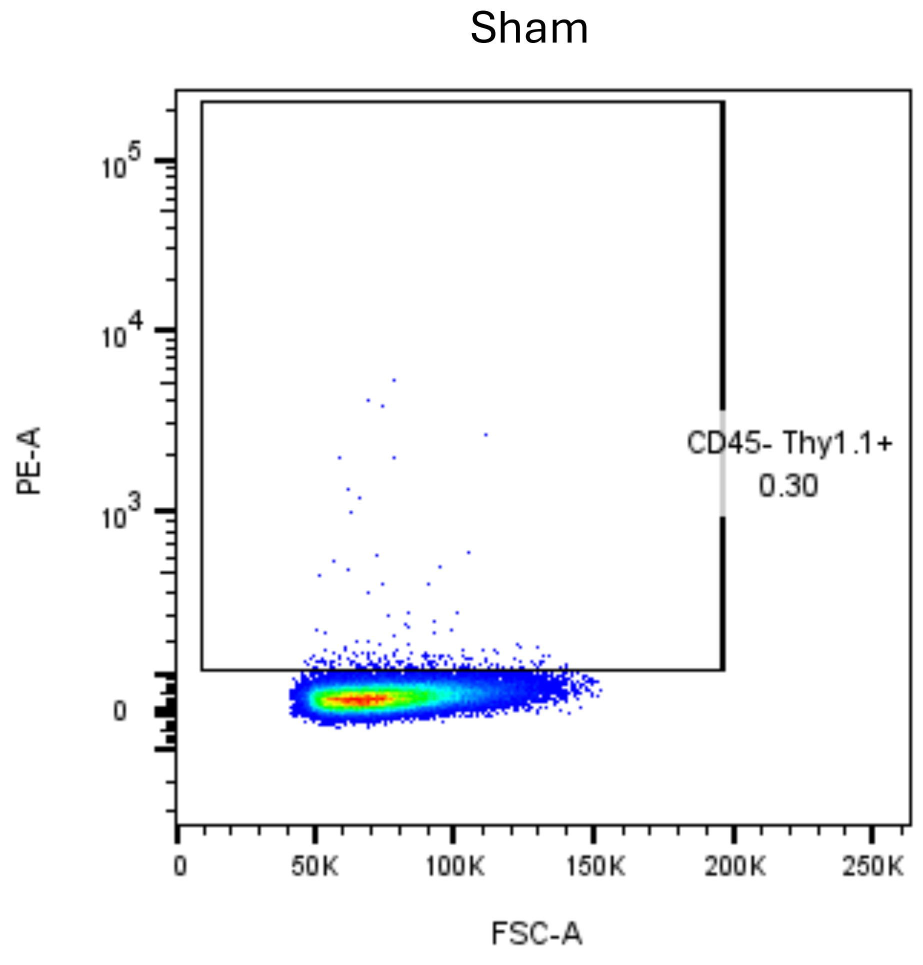

Supplementary Figure S9

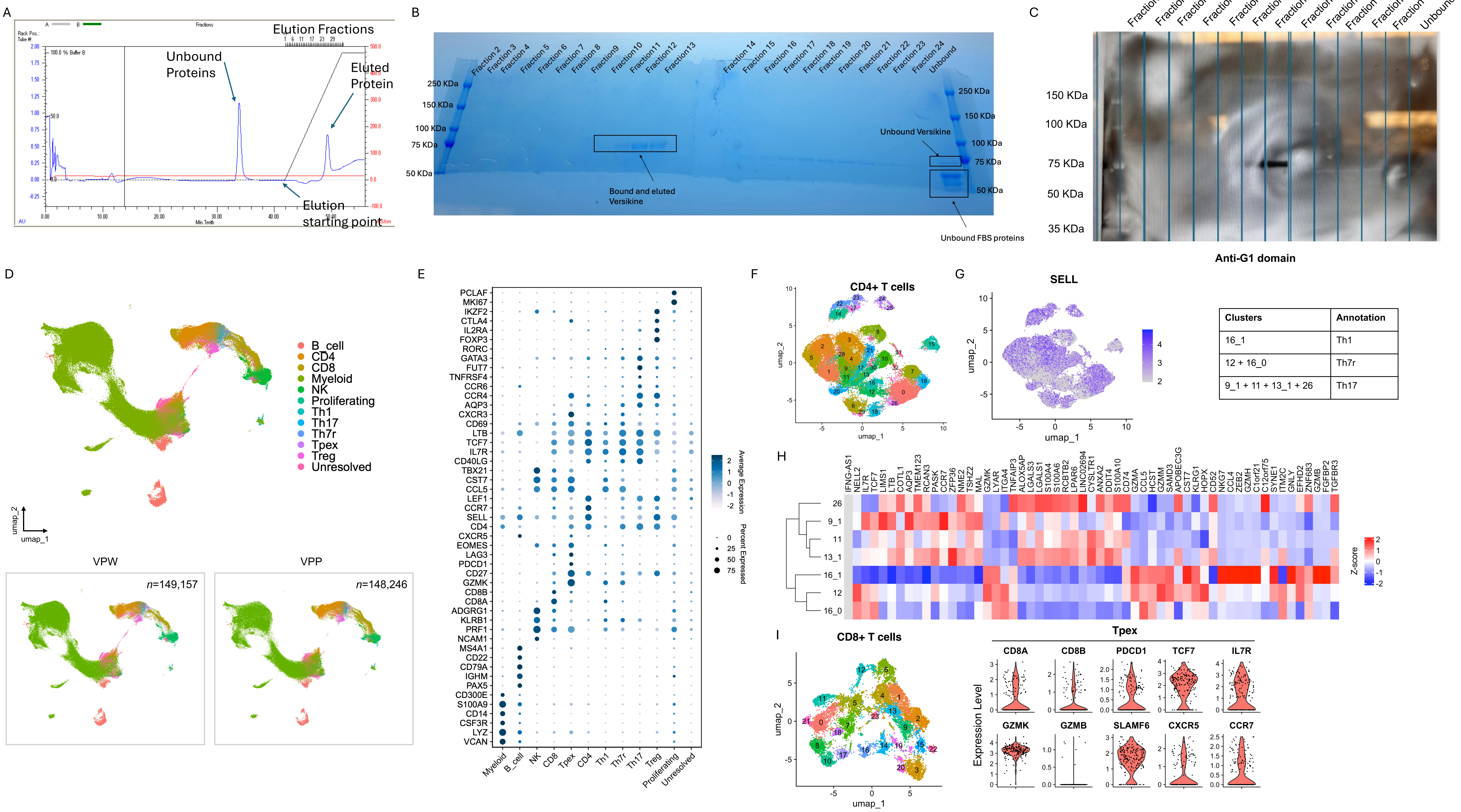


| Clusters | Annotation |
|---|---|
| 16_1 | Th1 |
| 12 + 16_0 | Th7r |
| 9_1 + 11 + 13_1 + 26 | Th17 |
