## Supplementary material for "A matrikine organizes the dendritic cell–T cell triads that build tertiary lymphoid structures": Supp. Fig. S

**SUPPLEMENTARY FIGURE LEGENDS (Lagal et al., 2026)**

**Supp. Fig. S1.**

A: Human tonsillar tissue IF using fluorescently-labeled anti-DPEAAE and anti-Ki67 (proliferation marker) antibodies demonstrate polarization of VCAN proteolysis within the non-proliferative light zone (LZ) of germinal centers (GC). 20X objective: scalebar 50 μm. B: DPEAAE staining overlaps with the extent of the follicular dendritic network in the LZ, highlighted by CD23 staining. 20X objective: scalebar 50 μm. C: Immunohistochemistry for intact VCAN demonstrates producer cells located mostly (albeit not exclusively) in the GC LZ. 10X objective: scalebar 220 μm; 40X objective: scalebar 90 μm. D: IF staining for CD68 and intact VCAN demonstrates VCAN production in a subset of tingible-body macrophages. 40X objective: scalebar 20 μm. E: IF staining demonstrates colocalization of the DPEAAE signal and CXCL13 gradient in the LZ of the GC, with tingible body macrophage distribution across the GC demarcated by CD68 staining. 40X objective: scalebar 100 μm; 40X objective: scalebar 20 μm. F: Ki67 and DPEAAE staining in a reactive human lymph node (inset, H&E). 40X objective: scalebar 20 μm. G: Density of total CXCL13^+^ cells/ mm^2^ stromal area (top) and CD68^+^CXCL13^+^ macrophages/mm^2^ stroma surface area (bottom) across VPP (n= 20 patients) and VPW (n= 61 patients) NSCLC patient samples. H: B cell aggregate frequency increases with stromal Vkine intensity. Proportion of tissue microarray cases containing CD20⁺ B cell infiltrate/aggregates by stromal Vkine (DPEAAE) staining intensity, with intensities 2 and 3 combined (n = 16, 34 and 41 cases respectively). Numerators and denominators are shown above each bar; error bars are SEM. I: Immunoblot for HA detection (HA-tagged Vkine) in total tumor lysates of EV and Vkine-expressing CT26, MC38 and LLC tumors. J: Density of CD11c, CD4^+^ T and CD8^+^ T cells across EV and Vkine-expressing CT26, MC38 and LLC tumors. Cell counts per area were quantified by HALO image analysis across individual 25 different optical slices 1 μm thick. Total cell numbers were extrapolated to mm^3^ tumor volume. Data are representative as mean ± SEM. Statistics were performed using unpaired Student’s t test unless otherwise stated. In G, statistics by Wilcoxon rank sum test with continuity correction. In H, exact Cochran–Armitage test for trend across the three ordered strata. Statistical significance is provided as the exact p value or denoted by an asterisk, *p < 0.05; **p < 0.01; ***p < 0.001, ****p < 0.0001. p values less than 0.05 were considered statistically significant.

**Supp Fig. S2.**

A: In vitro growth rates of CT26-EV, CT26-Vkine, MC38-EV and MC38-Vkine cell lines (n= 4 samples). B: Density of Ccr7+ DC across tumor models (n= 4 mice). C: Loss of flow cytometry-delineated cDC1 in tumors implanted in *Batf3*-null recipients (n= 3 mice). D: Flow cytometry gating strategy to delineate specific CD8 T cells against MC38-intrinsic MuLV-derived tumor antigen (p15E). E/F/G. CD45^+^ cells were sorted from tumors established with CT26 (E), MC38 (F), and LLC (G) cells engineered to express Vkine or EV. UMAPs showing 62,152 cells from CT26 (E), 59,870 cells from MC38 (F), and 24,075 cells from LLC (G) captured by scRNA-seq (Chromium GEM-X Single Cell 3’ Gene Expression v4, 10x Genomics). Main cell populations are indicated. Column charts indicate percentage of each cell population between conditions. H/I/J: Dot plots showing relative expression of lineage markers used for broad cell type annotation. Dot size represents percentage of cells expressing feature within cluster. K-P: Feature plots showing log2-transformed expression of Ptprc (Cd45) (K/M/O) and Vcan (L/N/P) across tumor models. Data are representative as mean ± SEM. Statistics were performed using unpaired Student’s t test. Statistical significance is provided as the exact p value or denoted by an asterisk, *p < 0.05; **p < 0.01; ***p < 0.001, ****p < 0.0001. p values less than 0.05 were considered statistically significant.

**Supp. Fig. S3.**

A/G/M: Feature plots showing module scores for a pan-DC gene signature (*Itgax, Flt3, H2-Ab1*), which was used to identify DCs for downstream analysis, and feature plots displaying module scores for canonical markers of DC subsets. Genes used for each module are described in the Methods. B/H/N: Dot plots showing selected genes from the top 25 *de novo* markers per DC subpopulation. C/I/O: Pie charts showing DC subpopulation composition as a percent of total tumor-infiltrating DCs (TIDC) in EV- and Vkine-conditions across tumor models. D/J/P: GSEA of Hallmark pathways in Vkine compared to EV in total DC across tumor models. Top terms with FDR-adjusted p-value < 0.05 are shown. E/K/Q: Heatmaps displaying scaled expression of genes across DC subpopulations, grouped into functional categories. F/L/R: Feature plots showing module scores for Type I interferon production (left) and Type II interferon production (right) gene signatures in EV- and Vkine-conditions across tumor models. Gene sets were obtained from Gene Ontology Biological Processes.

**Supp. Fig. S4.**

A/E/I: UMAPs showing T cell subpopulations after reclustering, across each cell line. Column charts demonstrate percent of each subpopulation across conditions. B/F/J: Confusion matrices showing percent overlap between our annotations and predicted cell types by ProjecTILs mouse T cell reference atlas. C/G/K: Feature plots showing log2-transformed expression of *Cd4, Icos, Cd40lg,* and *Foxp3*. D/H/L: Dot plots depicting selected genes from the top 25 *de novo* markers per T cell subpopulation.

**Supp. Fig. S5.**

A/E/I: Heatmaps displaying scaled expression of genes across selected CD8^+^ T subsets. B/F/J: Violin plots showing expression of effector molecules (*Gzmb, Prf1, Tnf, Ifng*), checkpoint receptors (*Pdcd1/PD-1, Ctla4, Havcr2/TIM-3, Lag3, Tigit*), and interleukin receptors (*Il27ra, Il2ra, and Il2rb*) across CD8^+^ effector populations in EV- and Vkine-conditions across tumor models. C/G/K: Feature plots showing log2-transformed expression of Tcf7 in EV- and Vkine-conditions across tumor models***.*** D/H/L: Pie charts showing the composition of the Tcf7^hi^ Cd8a^+^ fraction in EV- and Vkine-conditions across tumor models. ***p_adj < 0.001; **p_adj < 0.01; *p_adj < 0.05.

**Supp. Fig. S6.**

A/G/N: Volcano plots depicting significant differentially expressed genes in the Tfh/Tfh1 population across tumor models. Red indicates upregulated genes in Vkine and blue represents downregulated genes. Highlighted genes are related to the Th1 regulon, including *Kdm6b/JMJD3, Ifng, Il12rb2, Il21r, and Stat4.* B/H/O: GSEA showing top T cell-related terms enriched in EV and Vkine. Terms were downloaded from Gene Ontology Biological Processes. FDR-adjusted p-values are shown on the x-axis. C/I/P: Feature plots showing Th7r module score in the Sell^low^ CD4+ compartment. D/J/Q: Feature plots showing log2-transformed expression of *Tbx21*/T-bet across conditions. E/K/R: Violin plots showing effector memory scores in the Tfh/Tfh1 population using a published gene set from Zhang et al., *Cancer Cell* 2021. Heatmaps showing average expression of genes in EV and Vkine conditions across tumor models. FDR-adjusted p-values were determined from GSEA and indicated by asterisks. F/L/M/S: Circle plots depicting CXCL9-CXCR3 and CXCL10-CXCR3 signaling (F/M/S) and ICOSL-ICOS (L) from DC subsets to T cell subsets. Arc color indicates sender population identity; thickness indicates relative signaling strength between populations; arrowheads indicate signaling direction. ***p*_adj < 0.01; **p*_adj < 0.05.

**Supp. Fig. S7.**

A: Immunofluorescence analysis of LLC tumors demonstrates no increase in CD4^+^T-bet^+^ T cells in Vkine-replete conditions (40X objective: scalebar 20μm). HALO analysis shows no physical proximity between CD11c DC and CD4^+^T-bet^+^ T cells in either LLC-EV or LLC-Vkine conditions. B: Anti-LAG-3 immune checkpoint inhibition in LLC-EV and LLC-Vkine models. Tumor growth curves shown as spider plots (thin lines) and composite (thick lines). C: Combination anti-PDL1/anti-LAG3 immunotherapy. Data are individual growth curves or mean, statistics by Mann-Whitney test. Statistical significance is provided as the exact p value or denoted by an asterisk, *p < 0.05; **p < 0.01; ***p < 0.001, ****p < 0.0001. p values less than 0.05 were considered statistically significant.

**Supp. Fig. S8.**

A: Efficient translation of Vkine-encoding mRNA delivered in complex with lipid-based transfection reagent (middle lane) or encapsulated in LNP (right lane). B: Dynamic light scattering shows average diameter of LNP (top). Electrophoretic light scattering shows average zeta potential of LNP (bottom). C: MC38 tumors were injected with LNP-Thy1.1 mRNA and dissociated for flow cytometry 24 hours later. Flow cytometric detection of surface Thy1.1 demonstrates efficient uptake of LNP and translation of synthetic Thy1.1 mRNA.

**Supp. Fig. S9.**

A: Representative chromatograph of recombinant Versikine purification through nickel-based immobilized metal affinity chromatography (IMAC). Elution was performed using a 5-column volume linear gradient of DMEM containing 500mM imidazole. Fractions containing isolated Versikine were identified based on UV absorbance spectra and validated through SDS-PAGE. B: SDS-PAGE of the different fractions obtained after protein purification, gel coomassie staining revealed the total protein content in each fraction. Vkine (75 KDa) was found in fractions 11-12-13 without the presence of unpurified proteins. Unpurified fraction (unbound) showed Vkine and the presence of additional protein from FBS. 11, 12 and 13 fractions were used to concentrate recombinant Vkine. C: Anti-VCAN (G1 domain) western-blotting of the different fractions obtained after protein purification. Antibody reveled Vkine (75kDa) in fractions 11-12-13. 11, 12 and 13 fractions were used to concentrate recombinant Vkine. D: UMAP showing 297,403 CD45+ cells captured by scRNA-seq (GEO accession number: GSE316301) from primary colon tumors of patients with the VPW (*n* = 5 pre-treatment, *n* = 7 post-treatment) or VPP (*n* = 6 pre-treatment, *n* = 6 post-treatment) phenotype. Cell populations are indicated by color. E: Dot plot showing canonical genes selected from the top 50 *de novo* markers for each population. F: The CD4+ T cells were reclustered into 32 *de novo* clusters for downstream analysis. G: Feature plot showing log2-transformed expression of SELL (*CD62L*). H: Heatmap showing expression of 58-gene signature for Th1, Th7r, and Th17 cells, applied to the SELL^low^ CD4+ T cell clusters. Signature was derived from Takei et al. *Nat Commun* 2026. I: The CD8+ T cells were reclustered. Violin plots depicting log2-transformed expression for the Tpex population (cluster 5_2): *CD8A, CD8B, PDCD1, TCF7, IL7R, GZMK, SLAMF6, CXCR5, CCR7,* with the absence of *GZMB*.
