## Supplementary material for "A matrikine organizes the dendritic cell–T cell triads that build tertiary lymphoid structures": Supp. Table S1

Supplementary Table S1.

Antibodies.

| Antibody | Company | Clone | Catalog Number | Dilution | Technique |
| --- | --- | --- | --- | --- | --- |
| Alexa Fluor® 700 anti-mouse CD8a Antibody | Biolegend | 53-6.7 | 100730 | 1:100 | Flow cytometry |
| Alexa Fluor® 700 anti-mouse CD3 Antibody | BioLegend | 17A2 | 100216 | 1:100 | Flow cytometry |
| Alexa Fluor® 700 anti-mouse I-A/I-E | BioLegend | M5/114.15.2 | 107622 | 1:100 | Flow cytometry |
| APC anti-mouse CCR7 | BioLegend | 4B12 | 120108 | 1:100 | Flow cytometry |
| APC anti-mouse CD8a Antibody | BioLegend | 53-6.7 | 100712 | 1:100 | Flow cytometry |
| Biotin-conjugated Donkey Anti-Mouse IgG | Jackson immunoresearch | Polyclonal | 715-065-151 | 1:1000 | Immunohistochemistry |
| Biotin-conjugated Donkey Anti-Rabbit IgG | Jackson immunoresearch | Polyclonal | 711-065-152 | 1:1000 | Immunohistochemistry |
| Biotin-conjugated F(ab')₂ Fragment | Jackson Laboratories | Polyclonal | 715-066-151 | 1:1000 | Immunohistochemistry |
| Brilliant Violet 605™ anti-mouse/human CD11b | BioLegend | M1-70 | 101257 | 1:100 | Flow cytometry |
| BUV395 anti-mouse CD45 | Thermo Fisher Scientific | 30F-F11 | 564279 | 1:100 | Flow cytometry |
| c-Myc | Cell signalling | D84C12 | 5605S | 1:3000 | Western blot |
| CD11c | Thermo Fisher Scientific | N418 | 53-0114-82 | 1:100 | Fresh-Frozen  Immunofluorescence |
| CD23 | Sigma-Aldrich | 1B12 | 123M-17 | 1:4 | FFPE Immunoflorescence |
| CD4 | Abcam | EPR19514 | ab183685 | 1:100 | Fresh-Frozen  Immunofluorescence |
| CD68 | Thermo Fisher Scientific | KP1 | 14-0688-82 | 1:100 | FFPE Immunoflorescence |
| CD8a | Thermo Fisher Scientiic | 53-6.7 | 14-0081-82 | 1:100 | Fresh-Frozen  Immunofluorescence |
| CXCL13 | R&D Systems | Polyclonal | AF801 | 1:75 | FFPE Immunoflorescence |
| Donkey anti-Goat IgG AF488 | Thermo Fisher Scientific | Polyclonal | A11055 | 1:250 | Immunohistochemistry and immunofluorescence |
| Donkey anti-Mouse IgG AF647 | Thermo Fisher Scientific | Polyclonal | A31571 | 1:250 | Immunohistochemistry and immunofluorescence |
| Donkey anti-Rabbit IgG AF594 | Thermo Fisher Scientific | Polyclonal | A21207 | 1:250 | Immunohistochemistry and immunofluorescence |
| FITC anti-mouse CD3 | BioLegend | 17A2 | 100204 | 1:100 | Flow cytometry |
| FOXP3 | Thermo Fisher Scientific | FJK-16s | 14-5773-82 | 1:100 | Immunofluorescence |
| GAPDH | Cell Sgnaling | 14C10 | 2118S | 1:3000 | Western blot |
| Goat anti-Chicken IgY AF488 | Thermo Fisher Scientific | Polyclonal | A11039 | 1:250 | Immunohistochemistry and immunofluorescence |
| Goat anti-Mouse IgG AF647 | Thermo Fisher Scientific | Polyclonal | A21235 | 1:250 | Immunohistochemistry and immunofluorescence |
| Goat anti-Rabbit IgG AF568 | Thermo Fisher Scientific | Polyclonal | A11011 | 1:250 | Immunohistochemistry and immunofluorescence |
| HA-Tag | Cell signaling  Technologies | C29F4 | 3724S | 1:1000 | Western blot |
| HRP- conjugate Anti-mouse IgG | Cell signaling | Polyclonal | 7076P2 | 1:2000 (WB)-1:500 (E) | Western blot and ELISA |
| HRP- conjugate Anti-Rabbit IgG | Cell signaling | Polyclonal | 7074P2 | 1:2000 | Western blot |
| InVivoMAb anti-mouse CD8α | BioXCell | 2.43 | BE0061 | 150μg | *In vivo* administration |
| InVivoMAb anti-mouse CTLA-4 | BioXCell | 9H10 | BE0131 | 100μg | *In vivo* administration |
| InVivoMAb anti-mouse IL-4 | BioXCell | 11B11 | BE0045 | 100μg | *In vivo* administration |
| InVivoMAb anti-mouse PD-L1 | BioXCell | 10F.9G2 | BE0101 | 100-200μg | *In vivo* administration |
| Ki67 | Thermo Fisher Scientific | Polyclonal | PA5-143573 | 1:100 | FFPE Immunoflorescence |
| Ki67 | Cell Signaling | 8D5 | 9449 | 1:800 | Immunohistochemistry |
| MuLV p15E Tetramer (KSPWFTTL) | MBL Life Science |  | TS-M507-1 | 1:10 | Flow cytometry |
| Pacific Blue™ anti-mouse CD11c | BioLegend | N418 | 117322 | 1:100 | Flow cytometry |
| PE anti-mouse/human CD45R/B220 | BioLegend | RA3-6B2 | 103208 | 1:100 | Flow cytometry |
| PE/Cyanine7 anti-mouse CD45 | BioLegend | 30F-F11 | 103114 | 1:100 | Flow cytometry |
| PE/Cyanine7 anti-mouse PD-L1 | BioLegend | 10F.9G2 | 124313 | 1:100 | Flow cytometry |
| PerCP/Cyanine5.5 anti-mouse/rat XCR1 | BioLegend | ZET | 148208 | 1:100 | Flow cytometry |
| Rat IgG2b Isotype Control | Sigma-Aldrich | RTG2B1-2 | SAB4702123 | 100-200μg | *In vivo* administration |
| T-bet/TBX21 | Cell signaling | E4I2K | 97135S | 1:250 | Fresh-Frozen  Immunofluorescence |
| TCF1/TCF7 | Cell signalling | C63D9 | 2203S | 1:250 | Fresh-Frozen  Immunofluorescence |
| TruStain FcX™ PLUS (anti-mouse CD16/32) | Biolegend | S17011E | 156604 | 1:50 | Flow cytometry |
| VCAN | Sigma-Aldrich | Polyclonal | HPA004726 | 1:100 | FFPE Immunoflorescence |
| VCAN (G1-domain) | DSHB | 12C5 | AB_528503 | 1:100 (WB) and 1:33 (E) | Western blot and ELISA |
| Versican V0, V1 (DPEAAE) | Thermo Fisher Scientific | Polyclonal | PA1-1748A | 1:100 | FFPE Immunoflorescence,Western blot and ELISA |
